# Cross-presentation of citrullinated antigens drives cytotoxic CD8^+^ T cell responses in rheumatoid arthritis

**DOI:** 10.64898/2026.06.03.729882

**Authors:** Jae-Seung Moon, Mengrui Zhang, Laura S. van Dam, Eun Kyung Song, Orr Sharpe, Julie A. Carman, Daniel C. Ramirez, Melanie H. Smith, Laura T. Donlin, Maureen C. Howard, Mark M. Davis, William H. Robinson

## Abstract

Rheumatoid arthritis (RA) is an autoimmune synovitis marked by anti-citrullinated protein antibodies (ACPAs) and infiltration of the synovium by activated immune cells. In ACPA-positive RA, CD8⁺ T cells are elevated in both the blood and synovium, and can be activated by MHC class I-restricted citrullinated autoantigens to mediate cytotoxic effector function. However, the mechanisms underlying the activation of cytotoxic CD8⁺ T cells in RA remain poorly understood. Here, single-cell transcriptomic and T cell receptor repertoire analysis of RA blood and synovial T cells revealed shared clonally expanded cytotoxic CD8⁺ T cell programs, with synovial enrichment of activated effector and proliferating populations and increased frequencies of *GZMB*⁺*IFNG*⁺ CD8⁺ T cells. We demonstrated that RA-associated oral bacteria stimulate neutrophil extracellular trap (NET) formation, leading to the peptidyl arginine deiminases (PAD)-dependent generation of extracellular citrullinated bacterial and host proteins. We further demonstrated that these antigens can be cross-presented to CD8⁺ T cells via HLA class I molecules expressed by monocyte-derived dendritic cells (MoDCs) and autoreactive ACPA-expressing B cells. Toll-like receptor (TLR) signaling, particularly TLR4 activation by citrullinated antigens, enhanced cross-presentation of citrullinated antigens and promoted CD8⁺ T cell activation and clonal expansion. In turn, citrullinated antigens stimulated autoreactive B cells to produce IL-8, which recruited CXCR1/2⁺ cytotoxic CD8⁺ T cells and amplified B cell-CD8⁺ T cell interactions. These findings reveal a mechanistic pathway linking microbial triggers, antigen presentation, and cytotoxic CD8⁺ T cell responses that may drive joint destruction in RA.

**One Sentence Summary:** TLR4-driven cross-presentation of citrullinated antigens by dendritic cells and autoreactive B cells promotes activation of CD8^+^ T cells in rheumatoid arthritis.

## INTRODUCTION

Rheumatoid arthritis (RA) is a chronic autoimmune disease characterized by synovial inflammation and joint destruction (*1, 2*). The presence of anti-citrullinated protein antibodies (ACPAs) (*3*) is a biomarker for RA with diagnostic utility. ACPAs recognize citrullinated epitopes generated through peptidyl arginine deiminase (PAD)-mediated conversion of peptidyl-arginine to peptidyl-citrulline in host and microbial proteins (*4, 5*). Neutrophil extracellular trap (NET) formation is one of the cellular processes that generates citrullinated proteins. Multiple risk factors are linked to ACPA^+^ RA, including periodontal disease, smoking, and HLA-DRB1 alleles that enhances presentation of citrullinated peptides to CD4^+^ T cells (*6–8*). Additionally, polymorphisms in MHC class I, such as the HLA-B*08, have been associated with ACPA^+^ RA (*9*), supporting a role for CD8^+^ T cells in RA pathogenesis. Indeed, RA blood and synovium frequently contain increased numbers of CD8^+^ T cells with effector phenotypes, and some studies have demonstrated their cytotoxic effector function (*10–14*). We recently identified that citrullinated autoantigens presented by host MHC class I can drive clonal expansion of CD8^+^ T cells from ACPA^+^ RA patients, leading to expression of cytotoxic mediators and killing of target cells (*15*). These findings suggest that autoreactive CD8⁺ T cells may directly contribute to synovial tissue destruction in RA.

Activation of conventional CD8^+^ T cells involves cross-presentation of exogenous antigens via MHC class I by professional antigen-presenting cells (APCs) such as dendritic cells (DC) (*16, 17*) and B cells (*18–20*). Typically, DCs are considered the most potent APCs for cytotoxic T lymphocyte (CTL) activation (*21*) as they possess specialized machinery for routing and loading exogenous antigens onto MHC class I molecules (*22, 23*). B cells can also cross-present exogenous antigen, however resting B cells tend to induce CTL tolerance (*24*), whereas activated B cells stimulate CTL responses by cross-priming (*25*). Further, autoreactive B cell receptors (BCRs) can internalize specific autoantigens, process them, and present peptides via MHC class I molecules to activate CD8⁺ T cells (*26*). Interestingly, TLR4 signaling promotes cross-presentation (*27*), and citrullinated proteins such as histone H3, vimentin, and fibrinogen can directly activate TLR4 pathway (*28–30*), suggesting that APCs in ACPA⁺ RA may efficiently cross-present these antigens via MHC class I.

The oral cavity of RA patients with periodontitis provides a source of citrullinated microbial antigens generated by host neutrophil PADs during NETosis (*31*). Citrullinated antigens are specifically recognized by the BCRs of ACPA-expressing B cells. Recently, isolation of ACPA-expressing B cells from RA patients using citrullinated peptide tetramers has revealed a relatively high-frequency population (up to ∼1 in 500 B cells), predominantly exhibiting a class-switched memory phenotype (CD20^+^CD27^+^IgD^−^), with some citrullinated antigen-specific plasmablasts (*32*). Kristyanto et al. (*33*) showed that ACPA-positive B cells displayed an activated, proliferative memory phenotype and reduced expression the inhibitory receptor CD32, suggesting heightened antigen-presenting potential. Based on these findings, we hypothesized that ACPA-expressing B cells in RA patients could act as cross-presenting APCs to autoreactive CD8^+^ T cells.

To investigate mechanisms of autoreactive CD8^+^ T cell activation and potential role of periodontal infections in activating them in RA, here we characterized CD8⁺ T cell states across RA blood and synovium using single-cell RNA sequencing (scRNA-seq) with T cell receptor (TCR) repertoire analyses, and identified shared clonally expanded cytotoxic CD8⁺ T cell programs, with synovial enrichment of activated effector and proliferating populations. Building on these observations, we demonstrated that RA-associated oral bacteria induce neutrophil NETosis, generating extracellular citrullinated antigens that can be cross-presented to CD8^+^ T cells by monocyte-derived dendritic cells (MoDCs) and ACPA-expressing B cells. Further, we showed that pharmacological inhibition of TLR4 or IRAK4 impairs cross-priming of cytotoxic CD8^+^ T cells by both MoDCs and ACPA-expressing B cells. Moreover, autoreactive B cells in response to citrullinated antigens secreted IL-8 to recruit CXCR1/2^+^ CD8^+^ T cells, forming B cell-CD8^+^ T cell clusters also observed in RA synovium. Together, these findings provide a mechanistic basis for the long-recognized association between periodontitis and RA, implicating citrullinated microbial antigens in the activation of innate and adaptive immune pathways that drive joint tissue destruction.

## RESULTS

### Activated, clonally expanded CD8⁺ T cell programs are shared across RA blood and synovium

To define disease-relevant CD8⁺ T cell states in RA and their relationship across blood and synovium, we performed scRNA-seq with TCR repertoire profiling of T cells isolated from paired RA synovium and blood samples, along with additional RA blood samples. Unsupervised clustering identified multiple CD8^+^ T cell populations, including cytotoxic, activated effector, interferon-stimulated genes (ISG)^hi^, and proliferating populations (Fig. 1A and fig. S1A and B). Notably, synovial T cells were enriched for activated effector and proliferating CD8⁺ T cell populations compared with blood (Fig. 1B and fig. S1C), consistent with ongoing immune stimulation within the joint microenvironment. In addition, the synovium contained an increased fraction of CD8⁺ T cells expressing *GZMB* and/or *IFNG*, a transcriptional phenotype corresponding to a CD8⁺ T cell population that we previously observed to expand upon stimulation with citrullinated antigens in RA PBMCs (*15*) (Fig. 1C and fig. S1D).

**Fig. 1.**
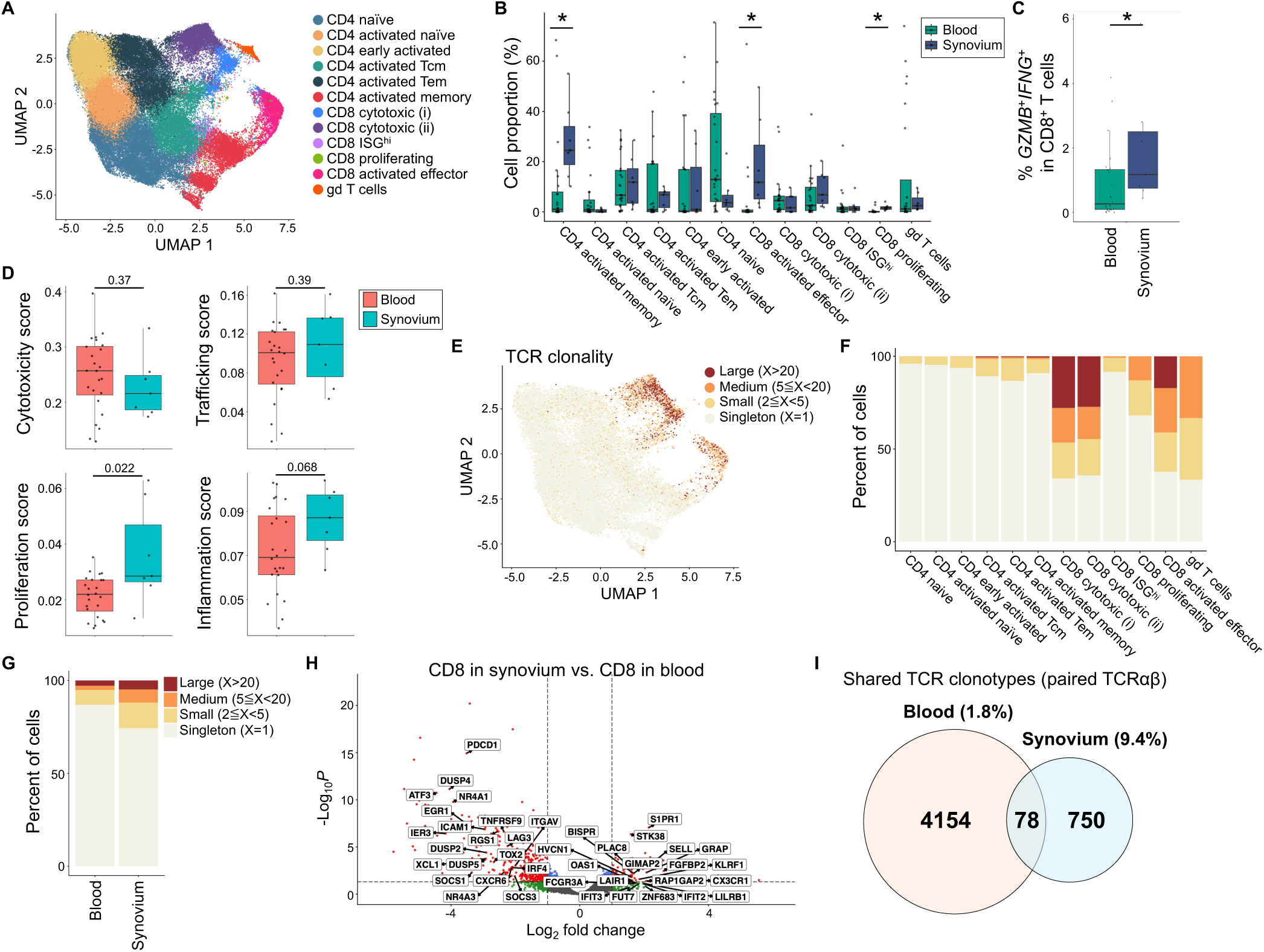
RA blood and synovium share clonally-expanded cytotoxic CD8⁺ T cell programs, with enrichment of activated and proliferating states in the synovium. **(A)** UMAP plot of integrated T cells from RA blood and synovium scRNA-seq datasets (*n* = 94,903 cells; 24 blood samples, 7 synovium samples from ACPA^+^ RA patients). Cell clusters were annotated based on canonical markers. **(B)** Proportion of T cell clusters in RA blood versus synovium. **(C)** Percentage of *GZMB*⁺*IFNG*⁺ CD8⁺ T cells in RA blood and synovium, defined as cells within CD8⁺ T cell clusters with *GZMB* > 0 and *IFNG* > 0. **(D)** Box plots showing cytotoxicity, trafficking, proliferation, and inflammation scores across RA blood and synovial CD8⁺ T cells. **(E)** UMAP plot of T cells integrated with TCR clonality based on paired TCRαβ sequences (n = 44,481 paired TCRαβ). Color indicates clonotype size as defined by the frequency of each unique paired TCRαβ sequence in the dataset. **(F)** Bar plot showing TCR clonality across all clusters. **(G)** Comparison of TCR clonality between RA blood and synovium. **(H)** Volcano plot of differentially expressed genes comparing expanded CD8⁺ T cell clones in synovium versus blood. Selected significant genes are highlighted. **(I)** Venn diagram showing overlap of unique paired TCRαβ clonotypes in CD8⁺ T cells from RA blood and synovium. Numbers denote blood-only (4,154), shared (78), and synovium-only (750) clonotypes. Statistical analysis was determined using pseudo-bulk level Bayes linear model in Limma (B-D). Data are plotted as means ± SD. *P < 0.05. ISG; Interferon-stimulated genes, Tcm; central memory T cells, Tem; effector memory T cells.

To compare functional gene programs of CD8⁺ T cells between blood and synovium, we calculated module scores using defined cytotoxicity, trafficking, proliferation, and inflammation signatures (*15*). Cytotoxicity and trafficking scores were comparable between blood and synovium, whereas proliferation and inflammation scores were increased in synovial CD8⁺ T cells (Fig. 1D). These data suggest that while cytotoxic programs of CD8⁺ T cell are broadly similar between blood and synovium, synovial CD8⁺ T cells exhibit features of sustained activation and local expansion within an inflammatory tissue environment.

To evaluate whether the activated CD8⁺ T cell states in RA reflect antigen-driven responses, we analyzed paired TCRαβ sequences to quantify clonal expansion. Clonal expansion was concentrated within cytotoxic CD8⁺ T cell populations (Fig. 1, E and F), with modestly increased clonality in the synovium as compared with blood (Fig. 1G). Consistent with compartment-specific state adaptation, expanded synovial CD8⁺ T cell clones showed increased expression of a sustained activation-associated program (e.g., *NR4A1*/*NR4A3*, *EGR1*, *ATF3*, *DUSP* family) together with elevated inhibitory receptors (e.g., *PDCD1*, *LAG3*) and tissue-associated features such as *CXCR6* and *XCL1*, whereas expanded blood CD8⁺ T cell clones were enriched for a circulating effector program (e.g., *CX3CR1*, *S1PR1*, *FGFBP2*, *KLRF1* and interferon-stimulated genes) (Fig. 1H). Together, these data identify shared clonally expanded cytotoxic CD8⁺ T cell programs across RA blood and synovium, with the synovium enriched for activated effector and proliferating states and increased transcriptional evidence of cytotoxic activation.

To quantify the extent of shared TCR repertoires across compartments, we identified 78 shared TCRαβ clonotypes between blood and synovium across 5 RA patients (9.4% of synovial clonotypes and 1.8% of blood clonotypes) (Fig. 1I). To examine how clonally expanded CD8⁺ T cells are related across RA blood and synovium and whether shared clones adopt distinct transcriptional programs, we performed TCR lineage analysis and visualized expanded clonotype families in a TCR lineage tree. This analysis revealed largely expanded clone families (lineages 1–5) that were shared between blood and synovium and were predominantly distributed among CD8 activated effector, cytotoxic, and proliferating clusters (fig S2). Notably, although cells within each lineage shared identical TCRαβ clonotypes, they frequently occupied different transcriptional clusters, indicating that shared clonal lineages can adopt distinct gene programs across microenvironments and activation states. Consistent with functional diversification, expression of cytotoxic effector genes varied across lineages: lineage 1 exhibited high expression of both *GZMB* and *GZMK*, lineages 2 and 5 were characterized by preferential *GZMK* expression, and lineages 3 and 4 showed preferential *GZMB* expression (fig. S2). Together, these data highlight that shared, expanded CD8⁺ T cell lineages in RA span multiple functional states, with lineage-specific granzyme programs and state-dependent transcriptional programs across blood and synovium.

Overall, these results indicate that RA blood and synovium share clonally expanded cytotoxic CD8⁺ T cell programs, with enrichment of activated effector and proliferating states in the synovium.

### RA-associated oral bacteria provide a source of citrullinated proteins via NETosis

We next investigated potential sources of citrullinated antigens and the processing pathways that enable CD8⁺ T cell activation in ACPA⁺ RA. The oral cavity of ACPA⁺ RA patients with periodontitis has been identified as a source of citrullinated commensal bacteria, predominantly the family of *Streptococcal* bacteria (*31, 34*). To investigate whether citrullinated oral bacterial proteins activate CD8^+^ T cells in ACPA^+^ RA blood, PBMCs from ACPA⁺ RA patients were stimulated with native or *in vitro* PAD4-citrullinated bacterial lysates from species previously linked to RA flares, including *Streptococcus gordonii* (SG), *S. oralis* (SO), *S. parasanguinis* (SP), and *Fusobacterium nucleatum* (FN) (*31*). Citrullinated bacterial lysates increased the frequency of granzyme B (GZMB)^+^IFNγ^+^ or CD107a^+^ CD8^+^ T cells compared to native lysates, with significant responses observed for citrullinated SP and SG lysates (Fig. 2A). In contrast, healthy control CD8^+^ T cells did not respond to the citrullinated bacterial lysates (fig. S3). Importantly, native lysates from these Gram-positive bacteria elicited minimal activation, unlike LPS stimulation, ruling out contributions from non-specific bacterial contaminants. These results demonstrate that citrullinated bacterial antigens derived from RA flare-associated oral bacteria can induce cytotoxic CD8^+^ T cell responses. For all subsequent experiments, *S. parasanguinis* (SP) was used as a model oral bacterium.

**Fig. 2.**
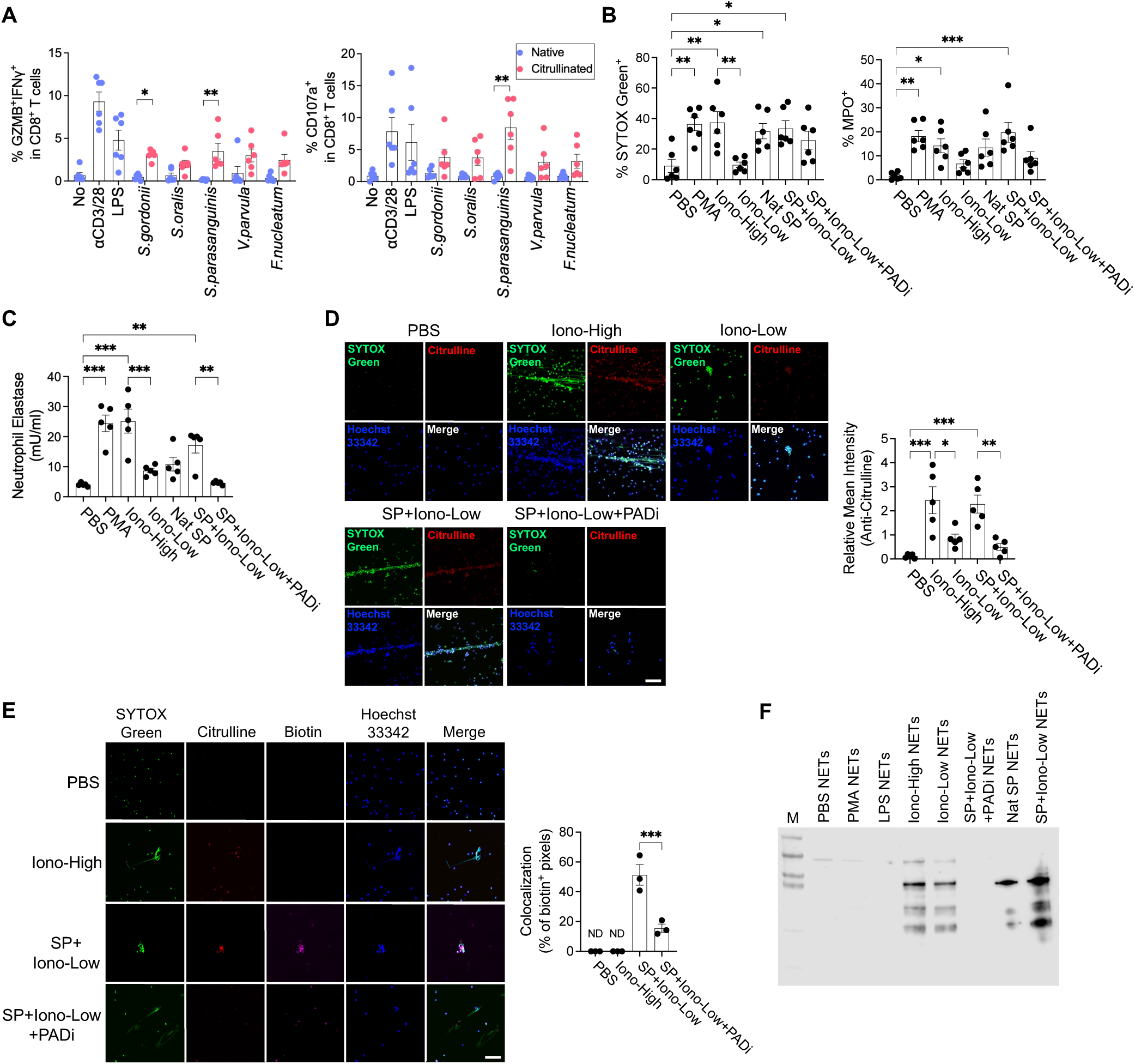
Oral bacteria-induced neutrophil NETosis generates citrullinated antigens in a PAD-dependent manner. (**A**) Proportion of GZMB^+^IFNγ^+^ (*left*) or CD107a^+^ (*right*) CD8^+^ T cells from ACPA^+^ RA PBMCs (*n* = 6) stimulated by native (*blue*) or citrullinated (*red*) oral bacterial lysates for 16 hr. (**B**) Frequencies of SYTOX Green^+^ (*left*) or MPO^+^ (*right*) population in ACPA^+^ RA blood neutrophils (*n* = 6) stimulated with each condition. (**C**) The level of neutrophil elastase secreted by ACPA^+^ RA neutrophils with each stimulation (*n* = 5). (**D** and **E**) ACPA^+^ RA neutrophils were treated with or without ionomycin high (1 μM) or low (0.01 μM) concentration, native *S. parasanguinis* (SP) lysates, or PAD inhibitor (PADi) Cl-amidine for 4 hr, and visualized to detect total citrullinated proteins (D), or biotinylated bacterial proteins (E) in the NETs with confocal microscopy. Relative mean intensity of anti-modified citrulline (D, *n* = 5) or percentage of colocalization (% of biotin^+^ pixels) (E, *n* = 3) was determined with image J. Scale bars, 50 μm. (**F**) Immunoblot of anti-citrullinated peptide antibody (ACPA) in NETs induced by each condition. SG, *Streptococcus gordonii;* SO; *Streptococcus oralis;* SP; *Streptococcus parasanguinis;* VP, *Veillonella parvula;* FN, *Fusobacterium nucleatum.* Statistical analysis was determined using two-way ANOVA with Tukey’s test (A), or one-way ANOVA with Tukey’s multiple comparisons test (B-D). At least three independent experiments were performed. Data are plotted as means ± SEM. *P < 0.05. **P < 0.01, ***P < 0.001.

Having established that citrullinated antigens drive these T cell responses, we next sought to identify the physiological source of citrullinated proteins in the RA environment. To this end, we investigated whether *S. parasanguinis* could induce NET formation containing citrullinated proteins. Neutrophils from ACPA^+^ RA blood were stimulated with phorbol myristate acetate (PMA), high-dose (1 μM, Iono-High) or low-dose (10 nM, Iono-Low) ionomycin, native *S. parasanguinis* lysates, or *S. parasanguinis* lysates plus Iono-Low with or without the PAD inhibitor (PADi) Cl-amidine. Ionomycin, a calcium ionophore, was used to activate PAD enzymes to promote NETosis-mediated hypercitrullination. NET formation was quantified using SYTOX Green and myeloperoxidase (MPO) staining, and further confirmed by elastase release, a key mediator of NET formation through nuclear translocation. Strong NET formation was observed following PMA, Iono-High, or SP+Iono-Low stimulation, while PAD inhibition markedly reduced NETosis and elastase activity (Fig. 2, B and C). Immunofluorescence confirmed that NETs induced by high-dose ionomycin or SP+Iono-Low contain citrullinated proteins, while PAD inhibition prevented protein hypercitrullination (Fig. 2D). Biotinylated bacterial proteins co-localized with anti-citrulline antibody signals in SP+Iono-Low-induced NETs (∼50%), suggesting PAD-dependent citrullination of bacterial proteins as previously described (*31*) (Fig. 2E). Additionally, human recombinant ACPAs bound to these NETs, whereas ACPAs did not bind to NETs in the presence of PAD inhibitor, indicating that the NETs generated in response to native *S. parasanguinis* lysates with ionomycin expose citrullinated human and bacterial proteins in a PAD-dependent manner (Fig. 2F and fig. S4). Notably, low-dose ionomycin alone elicited minimal NETosis and weak citrullination, suggesting that *S. parasanguinis* components are primary drivers of NET formation, while ionomycin primarily facilitates the calcium influx required for PAD-dependent hypercitrullination. (Fig. 2, B-D and F). Together, these results suggest that RA-associated oral bacteria, such as *S. parasanguinis*, can induce NETosis, thereby providing a source of citrullinated proteins that may stimulate both adaptive and innate immune pathways.

### Citrullinated proteins stimulate TLR4 signaling in dendritic cells

Activation of TLR4 by microbial products triggers intracellular pathways that can enhance antigen uptake, processing, and cross-presentation of external antigens on MHC class I through modulation of endosomal compartments (*27*). NETs induced by micro-organisms contain extruded neutrophil DNA, PAD enzymes, and a mixture of citrullinated host and bacterial proteins. We investigated whether NET products including citrullinated proteins stimulate TLR4 signaling in MoDCs from RA PBMCs. Confocal microscopy showed that MoDCs phagocytosed both citrullinated host proteins (e.g., histone H3) and biotinylated bacterial proteins from SP+Iono-Low NETs, whereas citrullinated proteins were absent in the PAD inhibitor-treated NETs (Fig. 3A).

**Fig. 3.**
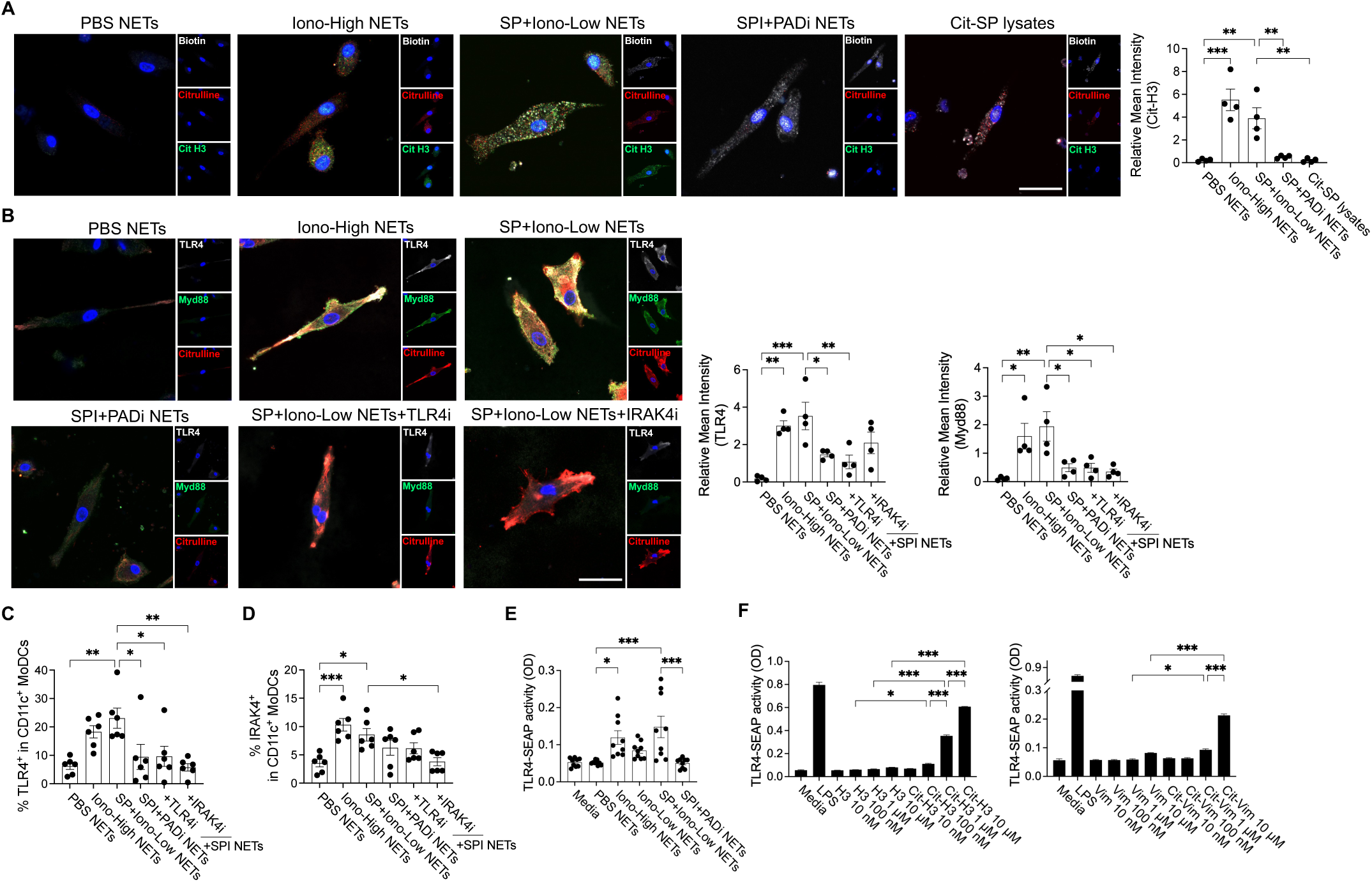
Citrullinated oral bacteria-induced NET components stimulate TLR4 signaling in dendritic cells. (**A** to **D**) Monocyte-derived dendritic cells (MoDCs) were treated with NET supernatants formed by PBS, ionomycin high dose (1 μM, Iono-High), native *S. Parasanguinis* (SP) lysates with low dose of ionomycin (0.01 μM), or SPI (native *S. parasanguinis* lysates with low-dose ionomycin) with PAD inhibitor (PADi) or intact citrullinated *S. parasanguinis* lysates, or SPI NETs in the presence or absence of TLR4 inhibitor TAK-242, or IRAK4 inhibitor zimlovisertib for 12 hr, and visualized to detect phagocytosed citrullinated proteins (A), or the level of TLR4 and MyD88 (B) in MoDCs by confocal microscopy. Relative mean intensity of cit-H3 (A, *right*, *n* = 4), TLR4, or MyD88 (B, *n* = 4) was determined with image J. Scale bars, 10 μm. The expression level of TLR4 (C, *n* = 6) or IRAK4 (D, *n* = 6) in CD11c^+^ MoDCs was assessed by flow cytometry. (**E** and **F**) HEK-Blue-hTLR4 cells were treated with each NET supernatant (E, *n* = 9) or native (Nat) or citrullinated (Cit) human proteins in different concentrations (F, *n* = 3) for 24 hr. The level of secreted embryonic alkaline phosphatase (SEAP) activity was measured in the supernatant. SPI; *S. parasanguinis* lysates with low-dose ionomycin. Statistical analysis was determined using one-way ANOVA with Tukey’s multiple comparisons test (B-F). Data are plotted as means ± SEM. *P < 0.05. **P < 0.01, ***P < 0.001.

We next measured TLR4, MyD88, or IRAK4 expression in MoDCs following incubation with the various NETs. To block TLR4 signaling in MoDCs, we pre-treated the TLR4 inhibitor TAK-242 or the IRAK4 inhibitor zimlovisertib. TLR4 and MyD88 expressions were significantly increased in MoDCs treated with the Iono-High or SP+Iono-Low NETs, but not with the PAD inhibitor-treated NETs (Fig. 3, B and C). Further, inhibition of either TLR4 or IRAK4 reduced TLR4 expression in response to the SP+Iono-Low NETs, while IRAK4 inhibition selectively downregulated IRAK4 expression (Fig. 3, B to D). These findings demonstrate that citrullinated proteins in NETs activates TLR4 signaling in MoDCs.

To further examine whether citrullinated proteins in the NETs directly activate TLR4 signaling, we utilized HEK-Blue hTLR cells, which express human TLRs and an NF-κB-inducible secreted alkaline phosphatase (SEAP) reporter. SEAP is produced upon cognate TLR ligand stimulation and can be measured by color change of culture medium. TLR4-expressing HEK-Blue cells showed a significant increase in SEAP production upon stimulation with the Iono-High or SP+Iono-Low NETs, compared to the PBS or PAD inhibitor-treated NETs (Fig. 3E). Since *S. parasanguinis* is a Gram-positive bacterium that has a pool of TLR2 ligands, the SP+Iono-Low NETs also induced SEAP production in HEK-Blue hTLR2 cells; however, this effect was not significantly different from the PAD inhibitor-treated NETs (fig. S5). We further confirmed that citrullinated histone H3 and vimentin dose-dependently increased TLR4-dependent SEAP activity compared to their native forms (Fig. 3F).

Together, these findings demonstrate that *S. parasanguinis*-induced NET-derived citrullinated proteins engage TLR4 signaling in MoDCs, providing a pathway for enhanced cross-presentation.

### Dendritic cells cross-present citrullinated antigens to activate and induce proliferation of cytotoxic CD8^+^ T cells in a TLR4-dependent manner

Since citrullinated proteins are internalized by MoDCs and activated TLR4 signaling, we investigated whether MoDCs cross-present citrullinated antigens in a TLR4-dependent manner. MoDCs were pre-treated with TLR4 or IRAK4 inhibitors, loaded with NETs or citrullinated human proteins, and then cocultured with autologous CD8^+^ T cells (Fig. 4A). SP+Iono-Low NET-loaded MoDCs markedly increased the frequency of GZMB^+^IFNγ^+^ or CD107a^+^ CD8^+^ T cells compared to PBS or PAD inhibitor-treated NETs (Fig. 4B). This activation was abrogated by pharmacological inhibition of TLR4 or IRAK4, similar to the blockade of HLA class I-dependent antigen presentation, indicating this activation is antigen- and TLR4-dependent (Fig. 4B). CD8^+^ T cell proliferation was also enhanced by the SP+Iono-Low NETs and was reduced by TLR4, IRAK4, or HLA class I inhibition (Fig. 4C). It is of note that a baseline proliferation was observed in control conditions due to the low-dose IL-2 supplementation required for T cell survival; thus, the specific response to SP+Iono Low NETs represents a significant increase over this background. Together with the observed HLA class I dependence, these findings reflect that the response is driven by NET-derived immunostimulatory components rather than residual ionophore effects in T cell activation. Thus, these results suggest following internalization by MoDCs, NET-derived citrullinated antigens are cross-presented to CD8^+^ T cells in an HLA class I-restricted manner, amplified by TLR4 signaling.

**Fig. 4.**
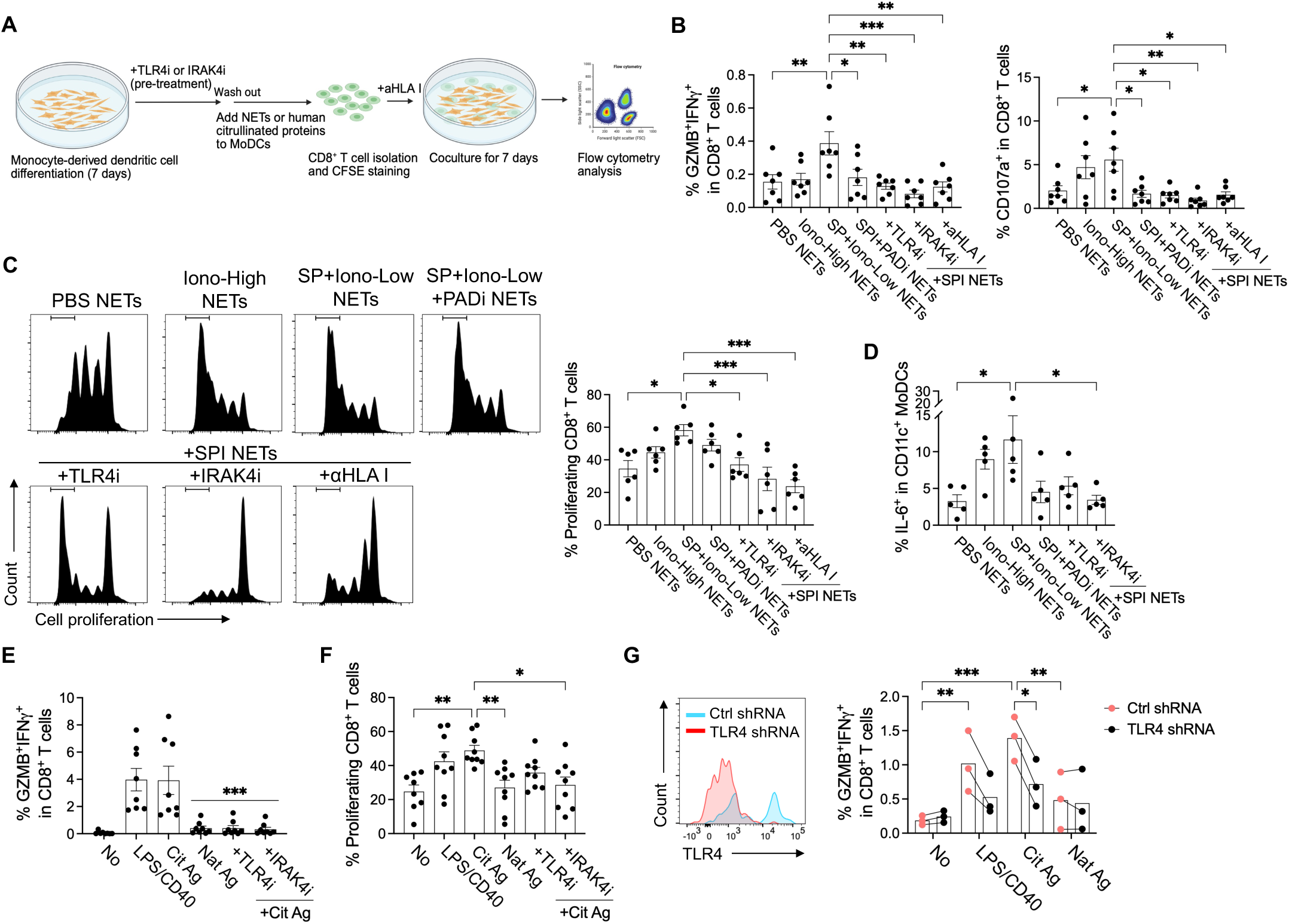
Cytotoxic CD8^+^ T cells are activated by cross-presentation of citrullinated proteins by dendritic cells in a TLR4-dependent manner. (**A**) Experimental schematic of coculture assay of in vitro MoDCs and CD8^+^ T cells from ACPA^+^ RA PBMCs in presence of NETs or human citrullinated proteins with or without TLR4 or IRAK4 inhibitor, or anti-HLA class I antibody. Figure created with BioRender. (**B**) Quantification of GZMB^+^IFNγ^+^ (*left*, *n* = 7), or CD107a^+^ (*right*, *n* = 7) CD8^+^ T cells in the coculture assay measured by flow cytometry. (**C**) Representative flow cytometry histograms (*left*), and frequency (*right*) of proliferating CD8^+^ T cells gating with cell proliferation dye eF450 (*n* = 6). (**D**) Percentage of IL-6-expressing CD11c^+^ MoDCs in the coculture assay (*n* = 5). (**E** and **F**) Human native- or citrullinated proteins were incubated with MoDCs in the presence or absence of inhibitors and cocultured with CFSE-stained CD8^+^ T cells. LPS and anti-CD40 antibody were used as a positive control to activate MoDCs. Proportion of GZMB^+^IFNγ^+^ (E), or proliferating (F) CD8^+^ T cells was measured using flow cytometry. * indicates versus Cit Ag (E). *n* = 8. (**G**) Representative histogram showing *TLR4* shRNA knockdown efficiency in MoDCs (*left*) or quantification of GZMB^+^IFNγ^+^ CD8^+^ T cells (*right*) in the coculture of control (*red*) or *TLR4* (*black*) shRNA-treated MoDCs with autologous CD8^+^ T cells. Each dot represents an independent sample (*n* = 3). SPI; *S. parasanguinis* lysates with low-dose ionomycin. Statistical analysis was determined using one-way ANOVA with Tukey’s multiple comparisons test (B-F), or two-way ANOVA with Fisher’s LSD test (G). Data are plotted as means ± SEM. *P < 0.05. **P < 0.01, ***P < 0.001.

Stimulation with SP+Iono-Low NETs also induced upregulation of IL-6 and antigen-presentation markers CD80 and HLA-DR in MoDCs, an effect which was attenuated by TLR4 or IRAK4 inhibition (Fig. 4D and fig. S6). Further, citrullinated histone H3 and vimentin induced CD8⁺ T cell activation and proliferation comparable to LPS+anti-CD40 antibody, while TLR4 or IRAK4 inhibitors suppressed the responses, indicating that these responses are antigen-specific and driven by TLR4 signaling (Fig. 4, E and F). To explore whether *TLR4* deficiency in MoDCs could influence cross-presentation of citrullinated antigens, we performed a coculture assay with *TLR4*-deficient MoDCs using *TLR4* shRNA-containing viral particles and autologous CD8^+^ T cells. Knockdown of *TLR4* in MoDCs impaired cross-presentation of citrullinated antigens, leading to reduced CD8^+^ T cell activation compared to control shRNA-transfected MoDCs (Fig. 4G).

To explore underlying mechanism, MoDCs were pre-treated with inhibitors targeting antigen-processing pathways prior to citrullinated antigen loading. The increase in IFNγ^+^ or CD69^+^CD107a^+^ CD8^+^ T cells was markedly reduced by brefeldin A, primaquine, chloroquine, or lactacystin, implicating Golgi transport, endosomal trafficking, lysosomal degradation, and proteasomal processing (fig. S7, A and B). This result reveals the broad array of intracellular signal transduction events involved in this response. We next focused on how dendritic cells internalize citrullinated antigens. DCs are efficient at taking up external proteins through phagocytosis or Fc receptor-mediated endocytosis of immune complexes. We observed ACPA immune complexes (ACPA 66 or 74) with His-tagged citrullinated proteins enhance antigen internalization into MoDCs compared to citrullinated antigen alone or isotype control immune complexes (fig. S8). Blocking Fc gamma receptor inhibited ACPA immune complex-mediated antigen uptake, indicating that ACPAs facilitate internalization and subsequent cross-presentation of citrullinated antigens (*35*).

Together, these findings demonstrate that citrullinated antigens can be cross-presented by MoDCs to activate and expand CD8^+^ T cells, with responses strongly augmented by TLR4 signaling.

### Citrullinated antigen-specific B cells exhibit a superior cross-presenting function to activate CD8^+^ T cells

Since studies have shown that B cells can internalize external antigens via antigen-specific BCRs, and cross-present them to activate CD8^+^ T cells in autoimmunity or chronic infection (*26, 36–38*), we evaluated whether membrane ACPA-expressing B cells contribute to the cross-presentation of citrullinated antigens. To this end, we cocultured B cells and autologous CD8^+^ T cells from ACPA^+^ RA patients with NET preparations under this study in the presence or absence of TLR or IRAK inhibitors. Coculture with SP+Iono-Low NET-loaded B cells increased IFNγ-expressing CD8^+^ T cells, which was substantially decreased by IRAK4 inhibition, and partially by TLR4i or TLR7i (fig. S9A). SP+Iono-Low NETs also increased the proportion of IL-6- and CD86-expressing CD19^+^ B cells in an activation response that was substantially decreased by IRAK4 inhibition, and partially by the TLR4 inhibitor (fig. S9, B and C). Interestingly, inhibition of Bruton’s tyrosine kinase (BTK), a BCR signaling molecule, potently suppressed SP+Iono Low NETs-induced B cell activation and subsequent CD8^+^ T cell activation, suggesting BCR engagement by citrullinated proteins (fig. S9, A-C). To further evaluate the antigen-presenting role of B cells, we evaluated their phenotypes upon antigen stimulation. While native antigens showed minimal effect comparable to unstimulated controls, co-stimulatory molecules CD80 and CD86 were significantly upregulated on B cells in response to citrullinated human antigens (Fig. 5A). This activation was blocked by inhibition of TLR4, IRAK4, or BTK (Fig. 5A). Further, B cells loaded with citrullinated antigens upregulated the antigen presentation and activation markers HLA-DR, HLA class I, CD40, and pSyk (Tyr348), while TLR4, IRAK4 or BTK inhibition abrogated these responses (fig. S10, A to D). Thus, citrullinated antigens have a dual stimulatory effect to induce broad TLR4-mediated activation and antigen-specific BCR signaling to drive B cell activation and antigen-presentation.

**Fig. 5.**
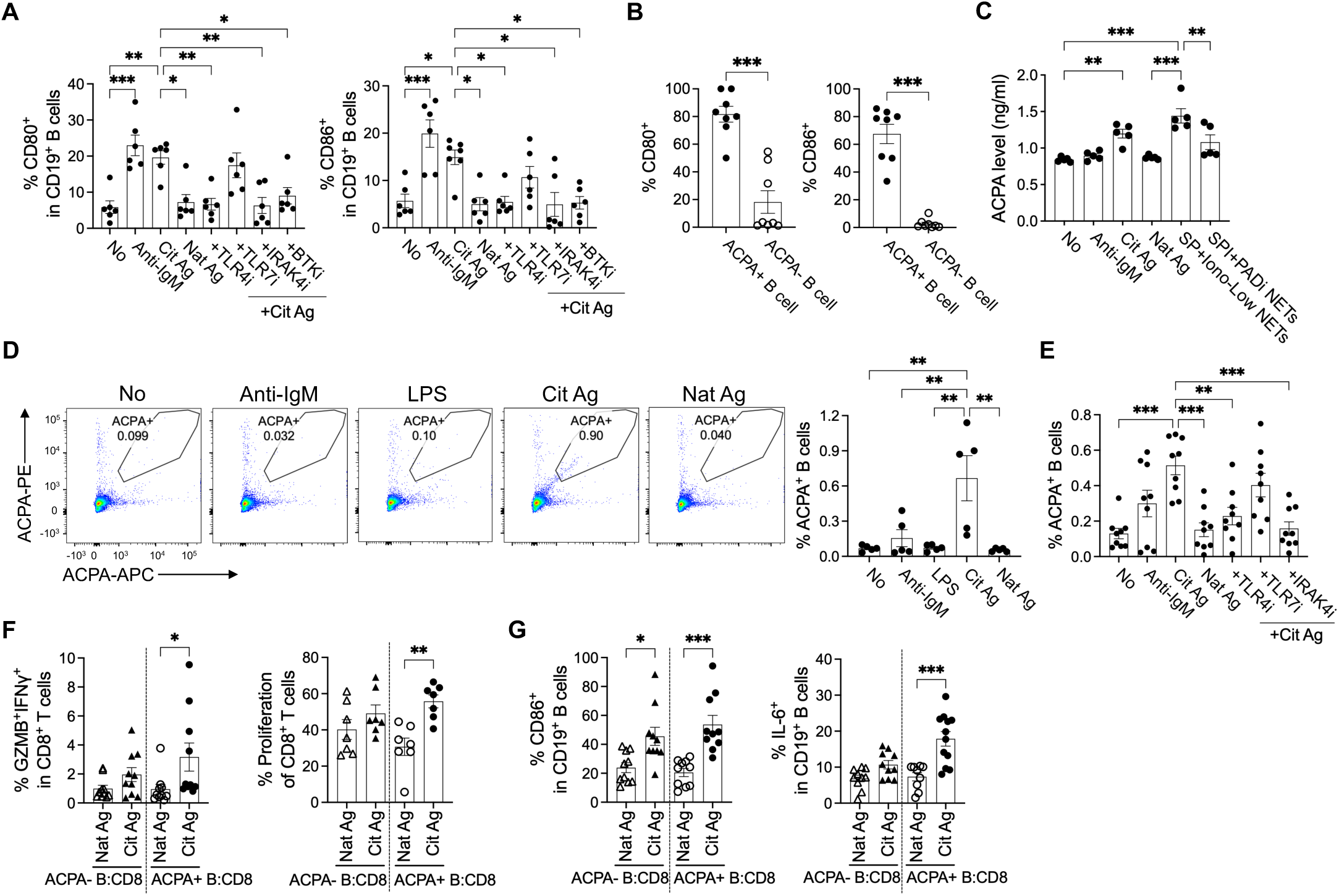
ACPA^+^ B cells can cross-present citrullinated antigens to activate CD8^+^ T cells. (**A**) Pan CD19^+^ B cells from ACPA^+^ RA PBMCs were treated with anti-IgM antibody, human native or citrullinated proteins, with or without the pretreatment of TLR4, TLR7, or IRAK4 inhibitor for 16 hr. Quantification of CD80 (*left*) or CD86 (*right*) in CD19^+^ B cells was measured using flow cytometry. *n* = 6. (**B**) Frequencies of CD80 (*left*), and CD86 (*right*) in ACPA-tetramer^+^ or - tetramer^−^ B cells. *n* = 8. (**C**) The secreted level of ACPA in the supernatants of pan B cells in response to each stimulation was quantified using ACPA ELISA. *n* = 5. (**D** and **E**) Percentage of ACPA^+^ B cells in pan B cells stimulated with anti-IgM antibody, LPS, or human native or citrullinated proteins in the presence or absence of each inhibitor. D; *n* = 5, E; *n* = 9. (**F**) Quantification of GZMB^+^IFNγ^+^ (*left, n* = 10) or proliferating (*right, n* = 7) CD8^+^ T cells in the coculture of ACPA^+^ or ACPA^−^ B cells with cell proliferation dye-stained CD8^+^ T cells with human native- or citrullinated proteins. (**G**) Frequencies of CD86^+^ (*left*) or IL-6^+^ (*right*) CD19^+^ B cells in the coculture of FACS-sorted ACPA^+^ or ACPA^−^ B cells with CD8^+^ T cells. *n* = 10. Statistical analysis was determined using one-way ANOVA with Tukey’s multiple comparisons test (A, C-G), or Student’s unpaired t-test (B). Data are plotted as means ± SEM. *P < 0.05. **P < 0.01, ***P < 0.001.

We next explored which B cell subpopulation(s) exhibit a higher capacity for recognizing and cross-presenting citrullinated antigens. Given that ACPAs specifically capture citrullinated antigens, we examined whether membrane ACPA-expressing B cells exhibit enhanced antigen-presenting ability. Circulating RA-associated ACPA^+^ B cells are predominantly found in the population of switched memory B cells at a frequency of 0.03% +/− 0.04% (*32, 39*). To detect and characterize them in ACPA^+^ RA blood using flow cytometry, we used a citrullinated peptide-streptavidin tetramer labeled with dual fluorophores (PE and APC) (fig. S11A and table S1). These tetramers specifically detected B cells in ACPA^+^ RA blood, and the sorted tetramer^+^ B cells secreted substantial amounts of ACPA (fig. S12, A and B). We observed that the population of ACPA^+^ B cells exhibited increased frequencies of the CD19^+^IgD^−^CD27^+^ switched memory (SwMe) and CD19^+^IgD^+^CD27^+^ non-switched memory (NSM) subsets, along with a reduced proportion of the CD19^+^IgD^−^CD27^−^ double-negative (DN) and CD19^+^IgD^+^CD27^−^ naïve (N) subsets, as compared to ACPA-negative B cells (*40*) (fig. S11B). Notably, over 10% of ACPA^+^ B cells were CD19^+^CD20^−^CD27^+^CD38^hi^ plasmablasts, a population nearly absent in ACPA^−^ B cells (fig. S11C). ACPA^+^ B cells expressed higher levels of CD80 and CD86 compared to ACPA-negative B cells, suggesting that ACPA^+^ B cells have a higher capacity for antigen-presentation than ACPA-negative B cells (Fig. 5B). We further showed the expression levels of TLR2, TLR4 and TLR7 on ACPA^+^ B cells was 80%, 100%, and 50%, respectively, as compared to less than 5% on ACPA^−^ B cells (fig. S11, D to F). Strikingly, the level of TLR4 was substantially higher on the surface of ACPA^+^ B cells as compared to CD27^+^ memory B cells (fig. S11G). Functionally, ACPA^+^ B cells highly secreted ACPAs in response to citrullinated antigens and SP+Iono-NETs, an effect abolished by PAD inhibition in the NET conditions (Fig. 5C). In addition to ACPA secretion, citrullinated antigens also induced expansion of ACPA^+^ B cells, while native proteins had no effect. (Fig. 5D). Furthermore, neither LPS alone nor BCR crosslinking with anti-IgM induced this expansion, supporting concurrent antigen-specific BCR engagement and TLR4 signaling in response to citrullinated antigens. This expansion was reduced by TLR4 or IRAK4 inhibition, and not by TLR7 inhibition, confirming that TLR4 signaling is required for B cell activation and expansion in response to citrullinated antigens (Fig. 5E). Further, pitstop 2, which inhibits receptor-mediated endocytosis, significantly reduced endocytosis of His-tagged citrullinated proteins by B cells, suggesting that the antigen uptake is an active receptor-mediated process rather than non-specific binding (fig. S13A). Consistent with dual engagement of BCR and TLR4, these citrullinated autoantigens were preferentially internalized by ACPA^+^ B cells or age-associated B cells (ABCs), B cell subsets that have been demonstrated to express ACPA in RA (fig. S13B).

We next assessed the possibility of cross-priming function of ACPA^+^ B cells using a coculture assay with ACPA^+^ or ACPA^−^ B cells and autologous CD8^+^ T cells from ACPA^+^ RA PBMCs. Citrullinated antigen-loaded ACPA^+^ B cells significantly increased GZMB^+^IFNγ^+^ and proliferating CD8^+^ T cells as compared to native protein-loaded ACPA^+^ B cells (Fig. 5F). In contrast, ACPA^−^ B cells did not induce statistically significant changes, although a minor upward trend was observed likely due to residual B cells reactive to other citrullinated epitopes not targeted in our sorting strategy (Fig. 5F). Consistent with our bulk B cell results, citrullinated antigens induced ACPA^+^ B cell activation expressing CD86 and IL-6, which has been shown to drive autoimmune germinal center formation (*41*), as compared to native proteins in the coculture assay (Fig. 5G).

Together, these data demonstrate that autoreactive ACPA-expressing B cells are highly responsive to citrullinated antigens that induce BCR and TLR4-mediated B cell activation and proliferation, and these B cells cross-present to and activate antigen-specific CD8^+^ T cells to enhance their cytotoxic functions.

### B cell-mediated cross-primed CD8^+^ T cells are clonally expanded and highly cytotoxic

Using single-cell RNA (scRNA) and VDJ repertoire sequencing, we analyzed the transcriptome and clonality of ACPA^+^ RA CD8^+^ T cells and B cells in our coculture system to explore whether these CD8^+^ T cells undergo clonal expansion (Fig. 6A). Unsupervised clustering identified 7 distinct CD8^+^ T cell clusters, including naïve/memory, *CCR6*^+^ or *GZMB*^+^*GZMK*^+^ cytotoxic, *GATA3*^+^ inflammatory, *MKI67*^+^ or *PCNA*^+^ proliferating, along with a small fraction of inflammatory *IL4*^+^*IL17*^+^ CD4 (Fig. 6B and fig. S14). Integration of transcriptome and TCR repertoire data revealed significant clonal expansion in cytotoxic and proliferating CD8^+^ T cells, with a higher frequency of expanded clonal lineages observed in response to citrullinated antigens as compared to native proteins (Fig. 6, C and D). Notably, IRAK4 inhibition reduced these clonal expansions, supporting a role for TLR signaling in B cell-mediated cross-presentation (Fig. 6D). Among the top clonally expanded clones, specific TCR clonotypes exhibited marked expansion upon citrullinated antigen stimulation, supporting antigen-specific TCR-mediated recognition (Fig. 6E).

**Fig. 6.**
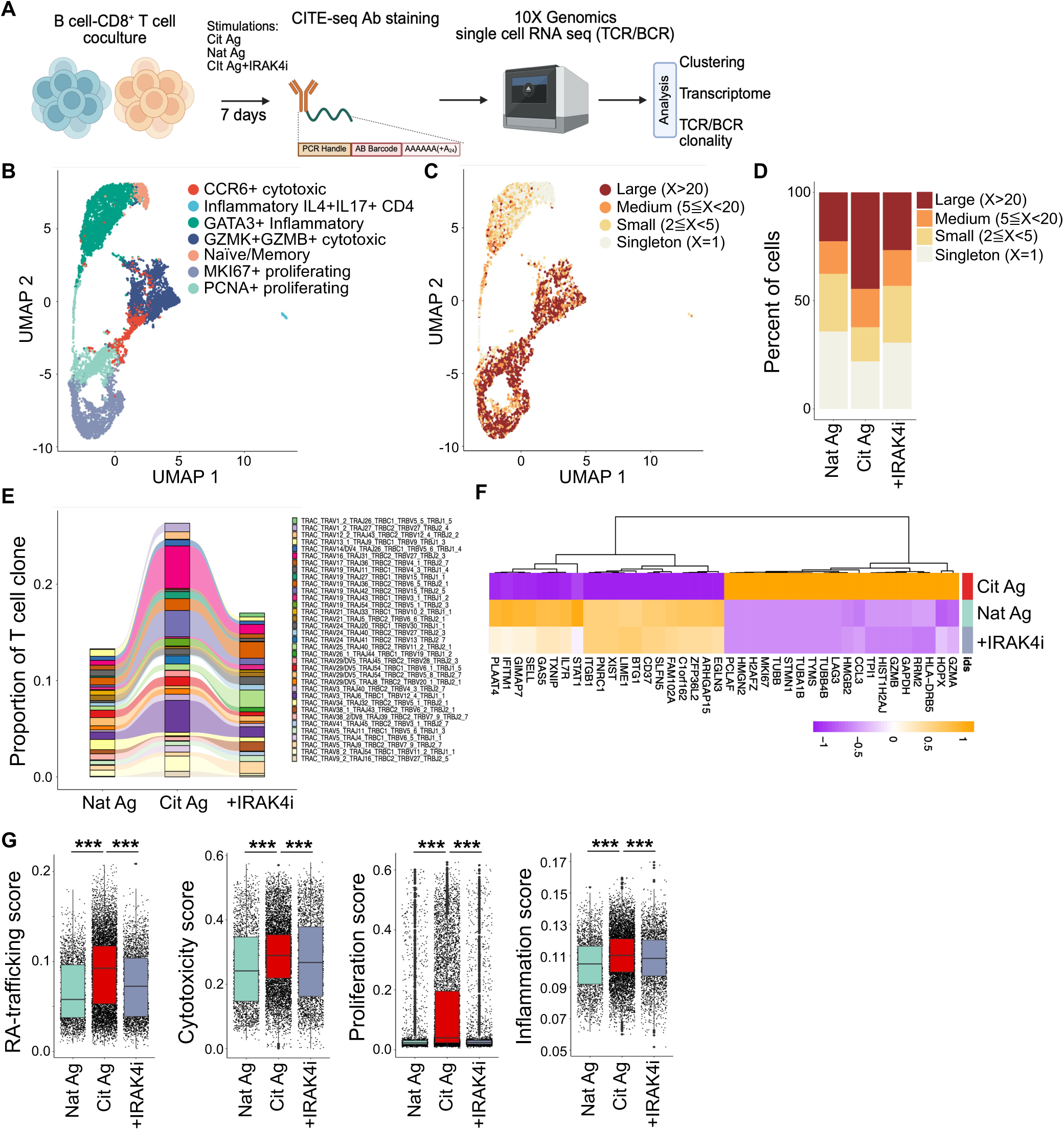
B cells induce clonal expansion of cytotoxic CD8^+^ T cells in response to citrullinated antigens. (**A**) Experimental scheme depicting single-cell RNA and VDJ sequencing of cocultured cells using magnetic sorted B cells stimulated by native- or citrullinated proteins with or without IRAK4 inhibitor and autologous CD8^+^ T cells. These cells were stained with CITE-seq antibodies and processed using the 10X Genomics Chromium platform. Figure created with BioRender. (**B**) UMAP plot of CD8^+^ T cells (*n* = 12,051) in the coculture assay with B cells and CD8^+^ T cells from 2 ACPA^+^ RA PBMCs. (**C**) UMAP plot of CD8^+^ T cells integrated with TCR clonality (*n* = 8,669 paired TCRαβ sequences). Color indicates the groups by the frequency of clonotypes in total cells. X represents the frequency of each clonotype defined by its unique paired TCRαβ sequence. Large-expanded clones (X>20), Medium-expanded clones (20>X≧5), Small expanded clones (5>X≧2), and Singletons (X=1). (**D**) Proportion of clonal lineages based on clonal size in stimulation with Nat Ag, Cit Ag, or Cit Ag with IRAK4 inhibitor. (**E**) Alluvial plot representing comparison of top 35 T cell clones from TCR repertoire in CD8^+^ T cells cocultured with B cells. (**F**) Heatmap showing expression level of differentially expressed genes between CD8^+^ T cells from Cit Ag vs. Nat Ag vs. Cit Ag with IRAK4i stimulations. (**G**) Box plots representing cytotoxicity, RA-trafficking, proliferation, and inflammation scores in CD8^+^ T cells stimulated by native or citrullinated proteins with or without IRAK4i. Each dot represents each cell. Statistical analysis was calculated with Wilcoxon rank-sum test (Mann-Whitney). The boxplot shows the median as a line in the middle of the box, with the first and third quartiles forming the box’s boundaries, while the whiskers extend to the minimum and maximum values in the dataset. ***P < 0.001.

Differential gene expression analysis revealed upregulation of cytotoxic (*GZMA*, *GZMB*) and proliferative (*MKI67*) genes in CD8^+^ T cells stimulated with citrullinated antigens, whereas cells exposed to native proteins or IRAK4 inhibition displayed naïve/memory-like gene signatures (Fig. 6F, fig. S15, A and B, and table S2 and S3). Consistently, these citrullinated antigen-stimulated CD8^+^ T cells exhibited higher gene scores of cytotoxicity, cell trafficking, proliferation, and inflammation (Fig. 6G).

We next analyzed the B cell transcriptome and BCR sequence data to characterize B cell responses to citrullinated antigen stimulation. Cluster analysis identified 5 distinct B cell clusters, including ABCs, germinal center B cells, activated naïve, plasmablasts, and proliferating plasmablasts (fig. S16, A and B). Citrullinated antigen stimulation increased expression levels of TLR4 and CD80 and CD86 on the B cell surface (fig. S16C). Moreover, citrullinated antigen stimulation induced expansion of proliferating plasmablasts and ABCs, which exhibited increased BCR clonal expansions as compared to other subsets (fig. S16, D to G). Together, these results demonstrate that B cells cross-present citrullinated antigens to activate cytotoxic CD8⁺ T cells, leading to antigen-driven clonal expansion.

### B cells recruit CD8^+^ T cells to facilitate contact-dependent antigen presentation *in vitro* and in RA synovium

To investigate whether B cell-mediated CD8^+^ T cell activation requires cell-cell interaction, we performed transwell migration assays with 0.4 or 5 μm pore inserts. 0.4 μm transwells prevent CD8^+^ T cell migration into the bottom well, while 5 μm pore transwells allow migration and subsequent contact between B cells and CD8^+^ T cells in the bottom well (Fig. 7A). B cells from ACPA^+^ RA blood in the bottom chamber were stimulated with either anti-IgM antibody, native, or citrullinated proteins, and autologous CD8^+^ T cells were loaded into the 0.4 or 5 μm transwell insert. CD8^+^ T cells migrated through the 5 μm pores and upregulated GZMB and IFNγ in response to B cells stimulated with citrullinated antigens or anti-IgM antibody, but not native proteins (Fig. 7B). In contrast, non-migrated CD8^+^ T cells in the insert did not respond to citrullinated antigen stimulation. B cells were also activated in response to citrullinated antigen stimulation (Fig. 7C). CD8^+^ T cells in the 0.4 μm insert were not activated, suggesting that B cell-mediated CD8^+^ T cell activation requires direct cell-cell interaction rather than soluble factors alone (Fig. 7D). B cells also showed reduced activation in the absence of interaction with CD8^+^ T cells in the 0.4 μm transwell, indicating bidirectional crosstalk between B cells and CD8^+^ T cells (Fig. 7E).

**Fig. 7.**
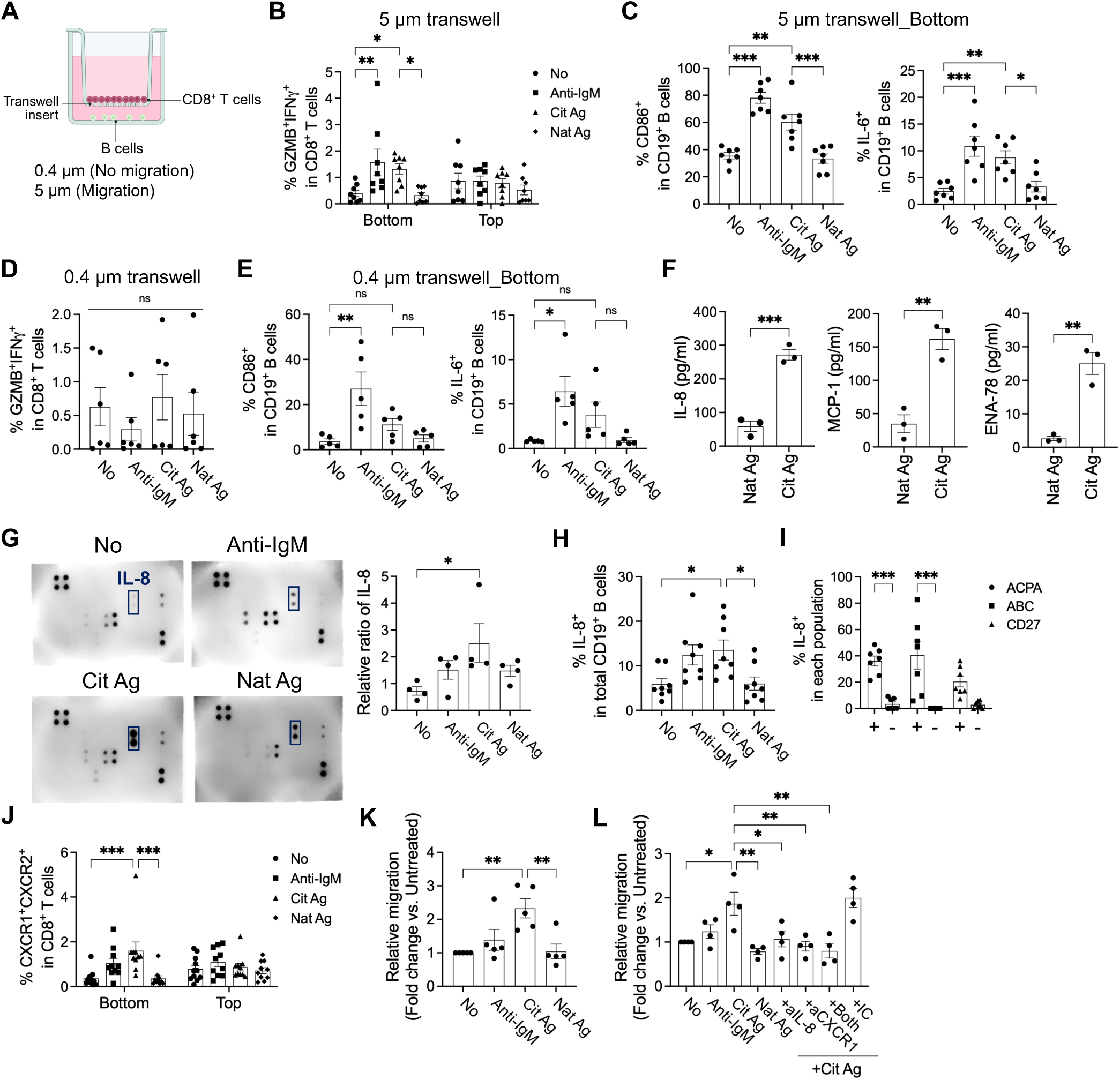
Citrullinated antigens stimulate ACPA^+^ RA B cells to secrete chemokines that attract CD8^+^ T cells to facilitate antigen presentation. (**A**) Experimental scheme representing the transwell assay with B cells in the bottom well and CD8^+^ T cells in the transwell insert. Figure created with BioRender. (**B** and **C**) Quantification of GZMB^+^IFNγ^+^ CD8^+^ T cells in the top (transwell insert) or bottom well (B, *n* = 8), or CD86^+^ (C, *left, n* = 7), or IL-6^+^ (C, *right, n* = 7) CD19^+^ B cells in 5 μm transwell assay with CD8^+^ T cells and B cells stimulated by anti-IgM antibody, native or citrullinated proteins for 7 days. (**D** and **E**) Percentage of GZMB^+^IFNγ^+^ CD8^+^ T cells in the transwell insert (D, *n* = 6), or CD86^+^ (E, *left, n* = 5), or IL-6^+^ (E, *right, n* = 5) CD19^+^ B cells in the bottom well of 0.4 μm pore transwell. (**F**) Bar graph showing the level of secreted chemokines in the cell culture supernatant of ACPA^+^ RA B cells stimulated by native or citrullinated proteins for 24 hr measured by Luminex. *n* = 3. (**G**) Representative immunoblots (*left*) and quantification (*right, n* = 4) showing human chemokine array with the cell culture supernatants of ACPA^+^ RA B cells stimulated by anti-IgM antibody, native or citrullinated proteins. Relative ratio was calculated based on the positive control of each membrane by Image J. The box represents a signal of anti-IL-8 antibody. (**H**) Quantification of IL-8^+^ CD19^+^ B cells in response to anti-IgM antibody, native or citrullinated proteins measured by flow cytometry. *n* = 8. (**I**) Percentage of IL-8-expressing B cells in each population comparing between ACPA^+^ vs. ACPA^−^ B cells, age-associated B cells (ABC) vs. non-ABC, or CD27^+^ vs. CD27^−^ B cells from ACPA^+^ RA patients. *n* = 7. (**J**) Pan B cells were treated with anti-IgM antibody, native or citrullinated proteins in the bottom well, and CD8^+^ T cells were seeded onto the transwell insert. Proportion of CXCR1^+^CXCR2^+^ CD8^+^ T cells in the insert (unmigrated) or bottom (migrated) of the transwell. *n* = 10. (**K**) The ratio of migrated CD8^+^ T cells from the transwell insert into the bottom well incubated with the culture supernatants of B cells. Untreated B cell supernatants were used as a standard. *n* = 5. (**L**) Bar graph representing the migrated cell ratio of CD8^+^ T cells from the transwell insert into the bottom well in the presence of B cells with or without each stimulation or the blocking antibodies. *n* = 4. IC; isotype control. Statistical analysis was determined using one-way ANOVA with Tukey’s multiple comparisons test (C-E, G-I, K, and L), two-way ANOVA with Tukey’s (B) or Sidak’s (J) multiple comparisons test, or unpaired t-test (F). Data are plotted as means ± SEM. *P < 0.05. **P < 0.01, ***P < 0.001. ns; not significant.

To explore how B cells facilitate CD8^+^ T cell migration, we analyzed chemokine secretion in bulk B cells stimulated with citrullinated antigens. By Luminex or chemokine array, we showed increased secretion of IL-8 in response to citrullinated proteins as compared with native proteins (Fig. 7, F and G). We further confirmed that the frequency of IL-8-expressing B cells is increased following citrullinated antigen stimulation using flow cytometry, and >40% of autoreactive B cells, including ACPA-expressing B cells or ABCs, express IL-8, which is consistent with previous findings (*33*) (Fig. 7, H and I).

We next asked whether CXCR1/2-expressing CD8^+^ T cells, receptors for IL-8, are recruited by citrullinated antigen-stimulated B cells. Migration assays revealed that CD8^+^ T cells expressing CXCR1 alone or CXCR1/CXCR2 were preferentially recruited by citrullinated antigen-stimulated B cells (Fig. 7J and fig. S17A). CXCR5^+^ or CX3CR1^+^ CD8^+^ T cells, reported in germinal center or RA synovium (*42, 43*), were also enriched among the migrated populations (fig. S17, B and C). Consistent with previous findings (*44, 45*), we showed CXCR1^+^CXCR2^+^ CD8^+^ T cells from ACPA^+^ RA blood exhibited enhanced cytotoxic potential, expressing GZMB, IFNγ, perforin, or granulysin (GNLY) as compared to double-negative CD8^+^ T cells (fig. S18, A and B). Further, this migration was enhanced by supernatants from citrullinated antigen-stimulated B cells (Fig. 7K and fig. S17D) and reduced by dual blockade of IL-8 and CXCR1, but not by isotype control (Fig. 7L and fig. S17E). These findings suggest that citrullinated antigens stimulate B cells to produce IL-8, which recruits CXCR1^+^ or CXCR2^+^ CD8^+^ T cells to facilitate B cell-CD8^+^ T cell interactions.

To investigate whether these interactions occur in RA synovium, we performed immunofluorescence on ACPA^+^ RA synovial tissues. We detected lymphocytic aggregates of B cells and CD8^+^ T cells, suggesting close spatial interaction that could support antigen presentation in ACPA^+^ RA synovium (Fig. 8A). While germinal center formation was not histologically detected in these samples, we observed these interactions in the lymphocyte aggregates across three patient samples, suggesting that these B cell - T cell interactions are occurring in the B cell aggregates that have been reported in a subset of RA synovium (*46–48*). Further, subsets of CD8^+^ T cells or B cells expressing CXCR1 or IL-8, respectively, were present in RA synovium (Fig. 8, B and C).

**Fig. 8.**
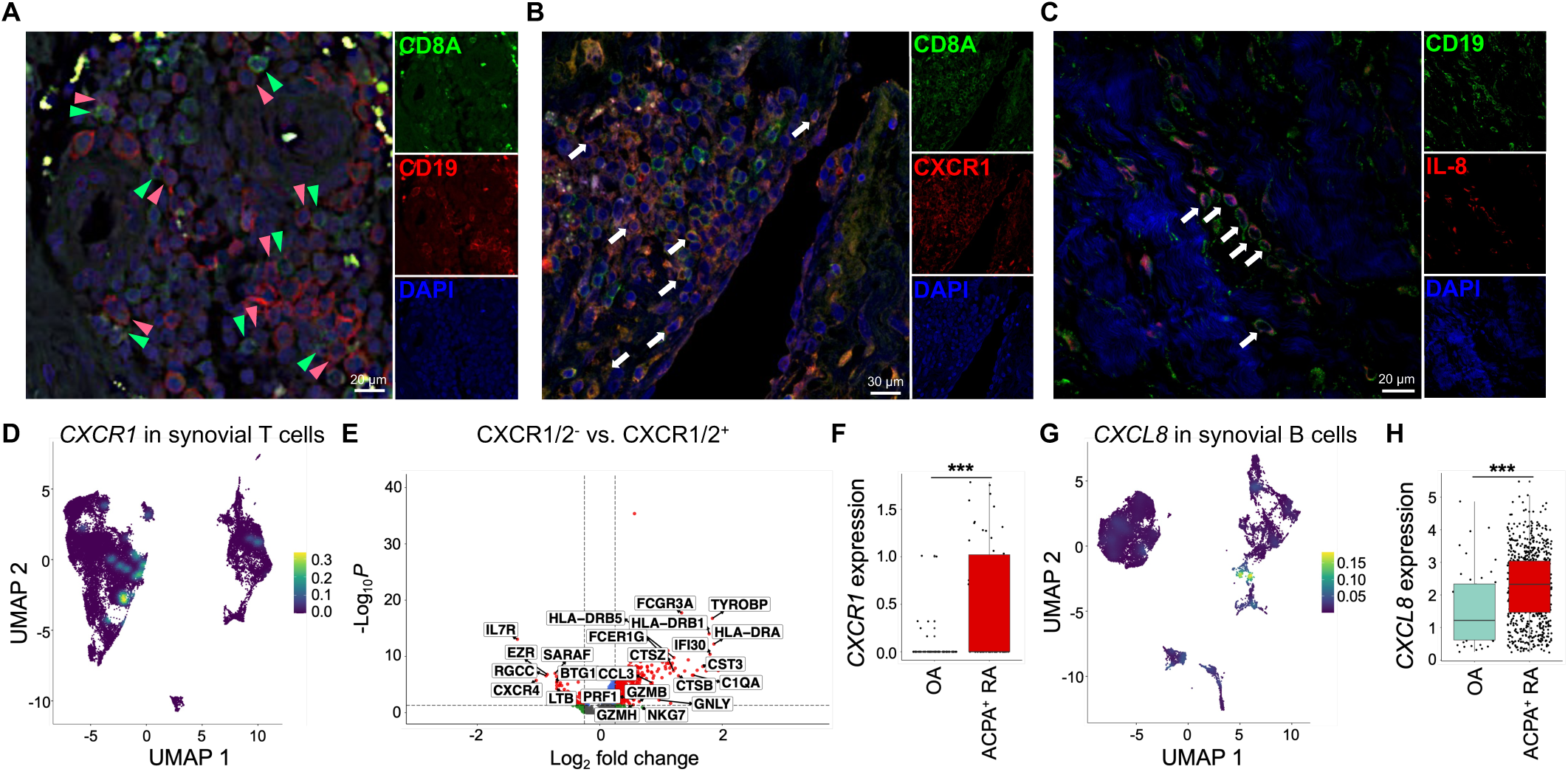
CXCR1^+^CD8^+^ T cells and IL-8^+^ CD19^+^ B cells are present in ACPA^+^ RA synovium. (**A** to **C**) Representative immunofluorescence images for CD8/CD19 (A), CD8/CXCR1 (B), or CD19/IL-8 (C) on formalin-fixed tissue sections from ACPA^+^ RA synovium. Arrows show CD8^+^ T cell (*green*) and B cells (*red*) in (A), CXCR1^+^CD8^+^ T cells in (B), or IL-8^+^CD19^+^ B cells in (C). Results are representative of three independent patient samples. Scale bars are indicated in each figure. (**D** to **F**) Single-cell RNA sequencing of CD3^+^ T cells (*n* = 44,032) isolated from ACPA^+^ RA or OA synovium (*n* = 5). Density plot of *CXCR1* in all CD3^+^ T cells (D). Volcano plot of differentially expressed genes comparing *CXCR1/2*^+^ CD8^+^ T cells with *CXCR1/2*^−^ CD8^+^ T cells (E). Expression level of *CXCR1* in all CD3^+^ T cells from OA and ACPA^+^ RA synovium (F). (**G**) UMAP density plot of *CXCL8* level in B cells (*n* = 27,013) from OA and ACPA^+^ RA synovium (*n* =5). (**H**) Box plot representing expression level of *CXCL8* in plasmablasts from OA and ACPA^+^ RA synovium. Statistical analysis was calculated with Wilcoxon rank-sum test. ***P < 0.001.

We next analyzed the transcriptomes of CD8^+^ T cells and B cells in ACPA^+^ RA or osteoarthritis (OA) synovium samples using scRNA sequencing. *CXCR1*^+^ cells were enriched in cytotoxic CD8^+^ T cells highly expressing cytotoxic mediators, including *GZMB*, *GZMK*, *IFNG*, or *GNLY* (Fig. 8D an fig. S19A). Differential gene expression analysis confirmed upregulation of cytotoxic (*GZMB, GNLY, PRF1*) and activation (*HLA-DR*) genes in *CXCR1*^+^ or *CXCR2*^+^ CD8^+^ T cells compared with *CXCR1*^−^*CXCR2*^−^ CD8^+^ T cells (Fig. 8E). *CXCR1* expression in synovial T cells was significantly increased in ACPA^+^ RA as compared to OA synovium (Fig. 8F), suggesting association with RA autoimmunity. Moreover, *CXCL8*, a gene for IL-8, was highly expressed in *CD27*^+^*CD38*^+^ B cells (Fig. 8G and fig. S19B), with levels markedly higher in plasmablasts from ACPA^+^ RA compared with OA synovium (Fig. 8H).

Together, these results suggest a potential mechanism whereby B cell-driven IL-8 promotes recruitment of cytotoxic CD8^+^ T cells, thereby amplifying synovial inflammation and tissue destruction in RA.

## DISCUSSION

There is increasing interest in the role of CD8^+^ T cells in autoimmune diseases including RA (*49–52*). CD8^+^ T cells can directly kill target cells, release cytokines that damage tissues, and may also suppress autoimmune responses by eliminating self-reactive cells or functioning as regulatory T cells (*53*). In our single-cell transcriptomic analysis with paired TCRαβ repertoire profiling of RA blood and synovial T cells, clonal expansion is preferentially enriched in cytotoxic CD8⁺ T cell populations across both compartments. We identify TCR clonotypes present in both blood and synovium, and observe that cells sharing an identical TCR frequently map to distinct transcriptional states rather than a single cluster. In addition, the synovium contains an increased fraction of *GZMB*⁺*IFNG*⁺ CD8⁺ T cells, consistent with the citrullinated antigen–responsive CD8⁺ T cell population we previously observed to expand in RA PBMCs (*15*). Building on this context, we demonstrate that MoDCs and ACPA-expressing B cells from RA patients cross-present MHC class I-restricted citrullinated antigens to activate autoreactive CD8^+^ T cells. We further demonstrate that citrullinated antigens activate TLR4, thereby enhancing cross-presentation and promoting cytotoxic CD8⁺ T cell activation. In parallel, autoreactive B cells secrete IL-8, which recruits CXCR1/2⁺ cytotoxic CD8⁺ T cells, reinforcing B cell-CD8⁺ T cell interactions within the synovium. Together, these findings reveal a mechanistic pathway linking microbial triggers, antigen presentation, and CD8⁺ T cell responses that may drive synovial tissue destruction in ACPA⁺ RA.

TLR4 is known to play an important role in the pathogenesis of RA (*54, 55*), and citrullinated proteins such as histone H3, vimentin, and fibrinogen can directly activate TLR4 (*28–30*). This activation promotes cross-presentation of exogenous antigens (*27*) by limiting phago-lysosome fusion, delaying antigen degradation, and prolonging MHC class I loading (*56*). Although *S. parasanguinis* is a Gram-positive bacterium that primarily engages TLR2, the observed activation by citrullinated bacterial lysates was significantly attenuated by TLR4 inhibition. Importantly, native lysates failed to induce comparable CD8^+^ T cell cytotoxicity. Furthermore, we confirmed in parallel experiments that the inclusion of polymyxin B (an LPS inhibitor) in the cell culture did not alter T cell responses to citrullinated antigens, excluding endotoxin contamination as a confounding factor. This indicates that the TLR4-dependent CD8^+^ T cell response is driven by citrullinated antigens rather than non-specific microbial products. Consequently, the IRAK4 inhibitor, a suppressor of all TLR signaling, induced a more profound suppression than the TLR4 inhibitor, suggesting that multiple TLRs likely contribute to this process (*57–59*). Effective B cell cross-presentation of external antigens is known to require B cell activation (*60*). Since TLR4 can activate naïve B cells, part of the TLR4-mediated augmentation may reflect expansion of the pool of activated antigen-presenting B cells. The TLR-induced cross-presentation mechanisms identified herein in RA may also be active in other autoimmune diseases in which CD8^+^ T cell responses mediates tissue injury, including systemic lupus erythematosus (SLE), myositis, type I diabetes, multiple sclerosis, and vitiligo (*61, 62*). In SLE, nuclear autoantigens are complexed with RNA or DNA, and in polymyositis, tRNA synthetase autoantigens are similarly associated with RNA, resulting in immunologically active autoantigens that stimulate TLRs and would thereby promote cross-presentation that activates cytotoxic CD8^+^ T cell responses contributing to autoimmune-mediated tissue destruction.

Our single-cell transcriptome and VDJ repertoire analysis provides a comprehensive view of the cellular dynamics governing this interaction. On the B cell side, citrullinated antigen stimulation induced a distinct differentiation trajectory toward Age-associated B cells and proliferating plasmablasts, which exhibited pronounced BCR clonal expansion and upregulation of TLR4 and costimulatory molecules. This transcriptomic signature suggests that citrullinated antigens, via co-engagement of the BCR and TLR4, license B cells into potent antigen-presenting cells. Concomitantly, citrullinated antigens promoted robust clonal expansion of cytotoxic CD8^+^ T cells compared with the modest expansion induced by native proteins, whereas IRAK4 inhibition markedly attenuated this response. These data suggest that while specific TCR engagement by citrullinated antigens provides the primary stimulus, B cell-intrinsic TLR signaling is also required to drive efficient cross-presentation and subsequent CD8^+^ T cell expansion.

We showed that activation of CD8^+^ T cells from RA patients by ACPA-expressing B cells presenting citrullinated antigens results in robust IL-8 secretion by B cells. Previous studies have shown IL-8 secretion by antigen-activated B cells to facilitate T cell recruitment into germinal centers (*63*), or by ACPA-expressing memory B cells in blood or synovial fluid to induce neutrophil migration (*33*). We observed that IL-8-secreting ACPA^+^ B cells recruit CXCR1/2^+^ CD8^+^ T cells, and these CD8^+^ T cells produce high levels of cytotoxic mediators, as previously described (*44, 45, 64*). IL-8 has also been known to mediate neutrophil migration, which may contribute to NET formation in response to ACPA immune complexes, thereby replenishing the pool of citrullinated antigens. Consistent with the proposed amplification loop, synovial T cell profiling reveals increased representation of activated effector and proliferating CD8⁺ T cell states and a higher frequency of *GZMB*⁺*IFNG*⁺ CD8⁺ T cells relative to blood. Thus, these findings support a pathogenic feedback loop that sustains CD8^+^ T cell activation and tissue damage in ACPA^+^ RA.

Histologic analyses of ACPA^+^ RA synovium used in this study revealed interstitial lymphocyte infiltrates but no distinct germinal center structures. This suggests that while CD8^+^ T cells and B cells interact within the synovium in ectopic lymphoid structures (ELS), these do not necessarily form bona fide germinal centers (GCs). Previous studies have demonstrated that CD8^+^ T cells as well as CD4^+^ T cells are essential for ELS formation in RA synovitis, and that these structures elicit antigen-specific responses (*46–48*). This implies that in RA synovium, antigen-specific B cells may activate autoreactive CD8^+^ T cells. Future studies will be needed to examine whether B cell - CD8^+^ T cell interactions in RA synovium directly induce cytotoxic function of autoreactive CD8^+^ T cells that mediate tissue destruction, using systems that resemble *in vivo* germinal centers, such as RA synovium organoids (*65*).

Synovial biopsies from RA patients revealed that a significant proportion of synovial CD8^+^ T cells express CD69 and CD103, markers of tissue-resident memory T (T_RM_) cells (*12*). These long-lived cells remain in previously inflamed joints during remission, and do not recirculate to other joints. In murine models of RA, site-specific depletion of CD8^+^ T_RM_-like cells during remission curtailed arthritis recurrence, suggesting an essential role in RA pathogenesis (*13*). However, the mechanisms by which cytotoxic CD8^+^ T cells are activated in ACPA^+^ RA have remained unclear. While most studies of CD8^+^ T_RM_ cells have focused on pathogen immunity, there is increasing evidence that CD8^+^ T_RM_ cells can also respond to self-antigens and mediate autoimmunity (*66–68*). Numerous studies investigating the mechanism of B cell depletion therapy in animal models of autoimmune disease have implicated elimination or suppression of autoreactive CD8^+^ T cell responses as a key mediator of efficacy (*69*) e.g. in type 1 diabetes (*70*), multiple sclerosis (*71*), and immune-mediated thrombocytopenia (*72*).

Therefore, our study provides a potential mechanism by which CD8^+^ T_RM_-like cells are sustained and activated via B cell-mediated cross-presentation of autoantigens in RA.

Consistent with heterogeneous tissue-associated CD8⁺ T cell states, TCR lineage analysis identified multiple, large expanded clonal families shared between blood and synovium that distribute across activated effector, cytotoxic, and proliferating clusters (fig. S2). Notably, despite shared TCR clonotypes, these lineages exhibit distinct transcriptional programs and granzyme patterns, with some lineages preferentially expressing *GZMK* and/or *GZMB*. These findings suggest that clonally expanded CD8⁺ T cells can diversify into multiple functional states within the synovial microenvironment, providing a cellular substrate for sustained cytotoxic potential in the setting of antigen presentation.

A limitation of this study involves the restricted spectrum of citrullinated antigens recognized by ACPAs included in our analysis. We utilized citrullinated H3 and vimentin as representative antigens, which have been previously shown to activate GZMB^+^ CD8^+^ T cells (*15*). However, the repertoire of citrullinated antigens in RA is vast. Consequently, our sorting strategy based on this specific antigen subset implies that the ACPA^−^ B cell fraction may still contain B cells reactive to other citrullinated epitopes not included in our probe cocktail. This likely accounts for the minor, non-significant increase of CD8^+^ T cell activation observed in the ACPA^−^ control group. Furthermore, different citrullinated antigens may possess distinct TLR-stimulatory activities and varying binding affinities for MHC class I alleles, potentially resulting in variable levels of CD8^+^ T cell activation. Future studies are needed to define which citrullinated antigens and epitopes most potently mediate APC cross-presentation and pathogenic CD8^+^ T cell activation in RA. Another limitation is that while most RA patients were sampled in an active disease state, approximately 40% were receiving immunosuppressive therapy at the time of sample collection. This could have affected the *in vitro* function of immune cells, possibly leading to underestimation of B cell and T cell responses. A further limitation is that synovial single-cell profiling in this study focuses on T cells, enabling high-resolution definition of CD8⁺ T cell states and clonality but not directly assess antigen-processing or TLR signaling programs within synovial antigen-presenting cells *in vivo*.

In summary, our study identifies a mechanism by which citrullinated antigens drive cytotoxic CD8^+^ T cell activation through cross-presentation by MoDCs and ACPA^+^ B cells, with IL-8 produced by autoreactive B cells promoting B cell - CD8^+^ T cell interactions in ACPA^+^ RA (fig. S20). These findings provide insight into RA immunopathogenesis and suggest that targeting pathways regulating cross-presentation or cytotoxic CD8^+^ T cell activation may offer therapeutic benefit in RA.

## MATERIALS AND METHODS

### Study design

This study was aimed to define how citrullinated antigens promote cytotoxic CD8⁺ T cell activation in ACPA⁺ RA through TLR-dependent cross-presentation. We analyzed RA blood and synovial T cells using single-cell RNA sequencing with paired TCR repertoire profiling to characterize CD8⁺ T cell states and clonal expansion. We performed mechanistic studies using primary human cells from ACPA⁺ RA or healthy donors. These included PBMC stimulation with native versus PAD4-citrullinated oral bacterial lysates or human proteins, neutrophil NETosis assays, and coculture systems using monocyte-derived dendritic cells or ACPA-expressing B cells with autologous CD8⁺ T cells to assess cross-presentation-dependent CD8⁺ T cell activation. We tested pharmacologic inhibition of TLR4 or IRAK4 where indicated and included endotoxin-control conditions when appropriate. We quantified IL-8 production upon antigen stimulation and tested CXCR1/2-dependent recruitment of cytotoxic CD8⁺ T cells using transwell migration assays. Sample sizes were determined by sample availability and prior experience, and no formal statistical method was used to predetermine sample size. Randomization and blinding were not performed. Data exclusion was limited to technical failures such as low cell yield. Experiments were repeated independently across donors as indicated in the figure legends.

### Human subject research

Clinical samples were collected from all human subjects in the present study. The study protocols were approved by the Institutional Review Boards (IRB) of the VA Palo Alto Health Care System and Stanford University, and written informed consent was obtained from all subjects prior to blood collection (IRB3780, table S4). Cryopreserved viable and formalin-fixed ACPA^+^ RA synovial tissue samples were obtained from Hospital for Special Surgery (HSS IRB 2014-233). All patients had active disease (Disease Activity Score >2.6) at the time of blood or synovial sample collection. ∼40% of patients received at least one of the immunosuppressives before sample collection. We excluded patients who had previously received B cell depletion therapy.

### Sample preparation and immune cell isolation

Whole blood was collected in heparin blood collection tubes (BD Biosciences). Peripheral blood mononuclear cells (PBMCs) were isolated via Ficoll-Paque density gradient centrifugation (Sigma Aldrich) using Leucosep tubes (Greiner Bio-One) for all experiments in this study. The isolated cells were cryopreserved in Recovery™ Cell Culture Freezing Medium (Thermo Fisher Scientific). Synovial tissues preserved in Cryostor CS10 (Sigma-Aldrich) were sourced from the Hospital for Special Surgery for ACPA^+^ RA samples or VA Palo Alto Health Care System for OA samples. The tissues were thawed completely at 37°C and then rinsed with RPMI 1640 (Corning Life Science) containing 10% FBS (ATCC) and 1% glutamine (Gibco). The tissue samples were finely chopped and subjected to enzymatic digestion using 100 μg/mL Liberase TL (Roche) and 100 μg/mL DNase I (Roche) in RPMI 1640 for 1 hr at 37°C with gentle inversion to break them down into single-cell suspensions. The resulting cell suspensions were passed through a 70 μm cell strainer and washed with cold RPMI 1640 supplemented with 10% FBS and 1% glutamine. These cells were then prepared for antibody staining and subsequent 10X single-cell RNA sequencing. CD8^+^ T cells, CD19^+^ B cells, or CD14^+^ monocytes from PBMCs were isolated using the EasySep Human CD8^+^ T cell Isolation kit, Human Pan-B cell Enrichment kit, or Human CD14 Positive Selection kit II (Stem Cell Technologies) according to the manufacturer’s instructions. Neutrophils were isolated from freshly collected whole blood from ACPA^+^ RA patients in the presence of 1 mM EDTA using the EasySep Direct Human Neutrophil Isolation kit (Stem Cell Technologies) and used immediately to minimize cell damage or death.

### Single-cell transcriptome and VDJ repertoire sequencing

For single-cell RNA and VDJ sequencing of CD8^+^ T cells or B cells from CD8^+^ T cell and B cell coculture assay, CD8^+^ T cells and pan B cells were isolated from ACPA^+^ RA patients and cocultured under the conditions described in the T cell proliferation assay section. After the coculture for 7 days, B cells and CD8^+^ T cells were stained with Human TruStain FcX and then CITE-seq antibodies (table S5) to detect protein expression on cell surface, and TotalSeq-C Hashtags (BioLegend) to distinguish each stimulation group in the analysis. For OA and ACPA^+^ RA synovium analysis, synovial cells were isolated as previously described and pan B cells or CD3^+^ T cells were isolated using the EasySep isolation kit, respectively. Single-cell suspensions were counted and loaded onto a Chromium Controller (10X Genomics), followed by library preparation for gene expression and TCR-seq with Chromium Next GEM Single Cell 5’ v2 (Dual Index) with Feature Barcode technology (10X Genomics) according to the manufacturer’s protocol. The cDNA library was pooled and sequenced using Illumina NovaSeq 6000 and the data were processed using Cell Ranger (v9.0.0) and aligned to the GRCh38 reference genome.

### Bioinformatic analysis of scRNA, scTCR-seq and scBCR-seq data

Downstream analysis of scRNA-seq data was performed in R using multiple Bioconductor packages (*73, 74*) and custom scripts, and included additional data from previously analyzed synovium samples with matched blood (accession PRJNA900189). Hashed libraries were demultiplexed through DropletUtils package (*75*). Data were integrated using scMerge package (*76*) and combined into a SingleCellExperiment (*74*) object. For quality control, cells were filtered through a flexible median absolute deviation (MAD) threshold using outlier detection function in Scuttle (*77*). Additionally, cells were further filtered out with >10% of mitochondrial genes, total UMI counts per cell <800 and >50,000 and total number of genes per cell <500 and >7,000. The filtered UMI count matrix was normalized and log-transformed through a deconvolution normalization strategy using the scran package (*78*). Principal component analysis (PCA) was performed on the top 1,500 high variable genes (HVGs) and the first 50 principal components (PCs) were used to perform Uniform Manifold Approximation and Projection (UMAP) for visualization. The Mutual nearest neighbors (MNNs) was performed for sample/batch correction and integration using batchelor package (*79*). Cell clustering was performed by *k*-nearest neighbor graph method (*k* = 30) using Louvain algorithm (*80*). CD8^+^ T cells were further defined based on the gene expression level of *CD8A*, and re-clustered with a range of k-nearest neighbors. Similarly, B cells were further defined based on the gene expression of CD19 and re-clustered. Top differentially expressed genes (DEGs) in each cluster were calculated using Wilcoxon rank-sum test with Bonferroni correction (*q* < 0.05). Each cluster was manually annotated based on the expression of canonical markers and DEGs. The gene expression levels of cytotoxicity, RA synovium trafficking, proliferation and inflammation-related genes, as described in the previous study (*15*), were scored using UCell package (*81*). The cell UMAP plots and heatmap plots were generated using dittoSeq package (*82*). The gene expression kernel density plots were generated using Nebulosa package (*83*).

For TCR and BCR repertoire analysis, both TCR and BCR contigs were annotated using the Cellranger vdj pipeline. BCR contigs were further re-annotated to IMGT human VDJ database using Change-O package (*84*). The TCR clonotypes were defined by the TCR alpha-and beta-chain with same gene arrangement. Similarly, the BCR clonotypes were defined by both BCR heavy and light chain with same gene arrangement, length of CDR3 sequences and similarity of CDR3 sequences (threshold: 75%). The size of clonal expansion was calculated by the count and proportion of each clonotype among total clonotypes in a patient. We define the levels of the clonal expansion for both TCR and BCR based on the clonal size (X) of each clonotype; Large-expanded clones (X>20), Medium-expanded clones (20>X ≧5), Small expanded clones (5>X≧2), and Singletons (X=1). The TCR and BCR clonotype data were further annotated and integrated into gene expression-based clusters to define gene expression profiles of clonally expanded cells. The TCR and BCR repertoire analysis was performed in R using immunarch, Dowser (*85*) and custom scripts. Alluvial plot for TCR was generated based on the top 20 expanded clones. For TCR clonal lineage tree analysis, TCR clonotypes shared between blood and synovial tissue were identified, and the full-length nucleotide sequence alignments of paired TRα and TRβ chains were concatenated per cell to produce a single representative sequence. These sequences were multiply aligned using MUSCLE (*86*). A maximum-likelihood phylogenetic tree was then inferred with RAxML (*87*). The resulting tree was visualized with ggtree, with outer annotation rings indicating tissue origin (blood vs. synovium), T cell subtype, clonal size category, and single-cell expression of *GZMK* and *GZMB*.

### Bacterial lysates preparation

As previously described (*31*), all bacteria strains were aerobically (*Streptococcus gordonii*, *Streptococcus oralis*, or *Fusobacterium nucleatum*) or anaerobically (*Streptococcus parasanguinis* or *Veillonella parvula*) grown to mid-logarithmic phase at 0.4-0.9 optical density at 600 nm (OD_600_) in America Type Culture Collection (ATCC) - recommended media and conditions. After the culture, the bacteria were pelleted by centrifugation at 6,000 g for 10 min, washed twice with phosphate-buffered saline (PBS, Corning), and the pellets were stored at - 20 °C. The frozen bacterial pellet was resuspended in B-PER lysis solution (Thermo Fisher Scientific) supplemented with 1× Halt protease inhibitor (Thermo Fisher Scientific), 0.05 μg/ml lysostaphin (Sigma Aldrich), and 40 μg/ml lysozyme (Sigma Aldrich). The mixture was incubated at 37°C with rotation for 45 min. Following incubation, samples were centrifuged at 20,000 g for 20 min at 4°C, and the resulting supernatants were collected. The supernatants were then buffer-exchanged into 1× Tris-buffered saline (TBS) overnight at 4°C and quantified using the Bicinchoninic Acid (BCA) assay (Thermo Fisher Scientific). For citrullination of bacterial lysates, the lysates were adjusted to a final concentration of 1 mg/ml in citrullination buffer (100 mM Tris, 10 mM CaCl₂, and 5 mM DTT) and subjected to *in vitro* citrullination by incubating with human PAD4 (0.5 unit/μl, Cayman Chemical) for 3 hr at 37°C. Control lysates were prepared by diluting them in citrullination buffer and incubating under identical conditions without PAD4. To assess PAD4-specific activity, an additional control was included in which PAD4 was incubated alone for 3 hr at 37°C in the absence of bacterial lysate. Citrullinated or native bacterial lysates were dialyzed into PBS and biotinylated using EZ-Link NHS-PEG4-Biotin (Thermo Fisher Scientific) according to the manufacturer’s instructions.

### Flow cytometry and cell sorting

Cells were washed with PBS to prepare single-cell suspension after cell culture at 350 g for 5 min and resuspended with FACS stain buffer (BD Biosciences) with Fixable Viability Stain 510 (BD Biosciences) and Human TruStain FcX (BioLegend) for 15 min at 4°C. To detach monocyte-derived dendritic cells (MoDCs) from bottom of each well, MoDCs were washed with cold PBS, and treated with cold PBS with 0.02% EDTA for 15 min in 4°C. Cells were washed with stain buffer, and stained with fluorophore-conjugated antibodies targeting cell surface molecules in stain buffer for 30 min at 4°C. Monoclonal antibodies for flow cytometry used in this study are listed in table S6. For chemokine receptor staining, the cells were incubated with chemokine receptor-targeting antibodies at 37°C for 20 min, and processed with the next step. For intracellular staining, isolated or cultured cells were treated with eBioscience Protein Transport Inhibitor Cocktail (Thermo Fisher Scientific) for the last 5 hr of the cell culture. And the cells were washed and subsequently fixed and permeabilized using eBioscience Fixation/Permeabilization kit (Thermo Fisher Scientific). The cells were stained with intracellular molecule-targeting antibodies (table S6) for 30 min at 4°C. Cells were acquired with BD LSRFortessa flow cytometer (BD Biosciences), and flow cytometric data were analyzed with FlowJo software (BD Biosciences). To define memory B cell subset, we gated IgD^−^CD27^−^ as a double-negative (DN), IgD^+^CD27^+^ as a non-switched memory (NSM), IgD^−^CD27^+^ as a switched memory (SwMe), or IgD^+^CD27^−^ as a naïve (N) among CD19^+^ B cells. For defining the plasmablast, we gated CD19^+^CD20^−^IgD^−^CD27^+^CD38^high^. For ACPA^+^ B cells and CD8^+^ T cell sorting from ACPA^+^ RA blood, frozen PBMCs were thawed in 37°C, and washed with FACS stain buffer to make single-cell suspensions. The optimal concentration of ACPA multimers (table S1) was pre-titrated. PBMCs were stained with Fixable Viability Stain 510, and TruStain FcX for 15 min, and further stained with anti-CD19, CD3, CD4 and CD8 antibodies, and ACPA multimers in stain buffer. Samples were sorted on a BD FACSAria II (BD Biosciences) or Sony SH800 Cell Sorter (Sony Biotechnology) with the gating strategy (fig. S9A).

### *In vitro* NET generation and collection of NET components

Purified neutrophils were resuspended in no phenol red Roswell Park Memorial Institute 1640 medium (RPMI, Corning life science), plated at 2 × 10^6^ cells/well in 12 well plate and stimulated with 100 ng/ml of phorbol myristate acetate (PMA, Sigma Aldrich), 10 nM (low) or 1 μM (high) of ionomycin (R&D Systems), or 2 μg/ml of native *S. parasanguinis* (SP) lysates for 4 hr at 37°C to induce neutrophil extracellular traps (NETs). 200 μM of the PAD inhibitor Cl-Amidine (Cayman Chemical) was pre-treated before stimulation of neutrophils for 30 min. After 4 hr, supernatants were gently collected and the NETs in the wells were mildly washed twice with 1 mL cold PBS. To harvest the NETs, the NETs were digested with 10 U/mL of micrococcal nuclease for 15 min at 37°C, and centrifugated at 300 g for 5 min at 4°C. Cell-free supernatants were collected and stored at −20°C for cell stimulation experiments. Quantification of neutrophil elastase was measured using the Neutrophil Elastase Activity Assay kit (Cayman Chemical). The pellets were used to detect and quantify NETosis using flow cytometry. The pellets were washed with PBS, and incubated with TruStain FcX for 15 min, and subsequently stained with anti-MPO antibody (BD Biosciences) and Sytox Green Ready Flow Reagent (Thermo Fisher Scientific) according to the manufacturer’s instruction. To investigate whether ACPA binds to the NET products, the pellets were stained with 1 μg/mL of human ACPA that was previously shown(*88*) as strong binding to human citrullinated proteins and anti-human IgG Fc secondary antibody-allophycocyanin (APC) (BioLegend). After washing with PBS, Sytox Green^+^, MPO^+^, or ACPA^+^ cells were detected using BD LSR Fortessa analyzer.

### Immunofluorescence staining

To study NET formation, human neutrophils were plated at 10^5^ cells/well in chambered coverslip (ibiTreat, ibidi), and NETois was induced as previously described. Before fixing the cells, the cells were incubated with 1 μM of SYTOX Green (Thermo Fisher Scientific) at 37°C for 15 min. The cells were fixed with 4% paraformaldehyde for 15 min at room temperature, and washed twice with PBS. Fixed neutrophils were incubated with a blocking buffer containing 1% normal goat serum (Thermo Fisher Scientific) in 1X TBS (Bio-Rad), and stained with primary antibodies, including anti-citrulline monoclonal antibody (clone 1G9, Cayman Chemical, 1:50 dilution), anti-Biotin polyclonal antibody (Thermo Fisher Scientific, 1 μg/mL), or anti-citrullinated histone H3 (citrulline R2+R8+R17) polyclonal antibody (Abcam, 1 μg/mL), in the blocking buffer at 4°C overnight. Then, neutrophils were washed with TBST containing 0.1% Triton-X, and stained with secondary antibodies, including anti-Rabbit IgG (H+L) secondary antibody-Alexa Fluor 488 (Thermo Fisher Scientific, 1:500), anti-Mouse IgG (H+L) secondary antibody-Alexa Fluor 594 (Thermo Fisher Scientific, 1:500), or anti-Goat IgG (H+L) secondary antibody-Alexa Fluor 647 (Thermo Fisher Scientific, 1:500), for 1 hr at room temperature. Then, slides were washed with TBST, stained with Hoechst 33342 (Thermo Fisher Scientific, 1 μg/mL) for 1 min at room temperature and mounted with Prolong Glass Antifade Mountant (Thermo Fisher Scientific). NET formation or the presence of citrullinated proteins was observed under Zeiss LSM 880 or Nikon confocal microscopy. For analyzing colocalization of biotin and citrulline, confocal images for biotin and citrulline signals were processed in Fiji (ImageJ). Each channel was background-corrected and thresholded using Otsu’s method. Binary masks were generated to identify positive regions, and the overlap area was quantified as the fraction of biotin-positive pixels colocalized with citrullinated signal: PBS and Iono-High groups, which lacked biotin signal, were designated as ND (no-biotin control).

For detection of dendritic cells, monocytes were seeded on the chambered coverslip and differentiated into monocyte-derived dendritic cells (MoDCs) for 7 days. After the differentiation, MoDCs were treated with NET products derived from each stimulation in the presence or absence of TLR4 or IRAK4 inhibitor. MoDCs were fixed, and blocked as previously described. MoDCs were incubated with anti-Biotin, citrulline, or citrullinated H3 antibody, or anti-TLR4 polyclonal antibody (Thermo Fisher Scientific, 10 μg/mL), or anti-MyD88 monoclonal antibody (clone 9R10V8, Thermo Fisher Scientific, 1:200 dilution) in the blocking buffer at 4°C overnight. Then, MoDCs were washed with TBST, stained with secondary antibodies, and Hoechst 33342. Protein internalization or TLR signaling was acquired by a confocal microscopy. Relative mean intensity was quantified with Image J software.

For detection of immune cell population in ACPA^+^ RA synovium, the paraffin-embedded synovial tissues were cut into 5 μm sections, and deparaffinized in xylene and rehydrated with absolute ethanol and gradual 95, 70, 50% of ethanol. The slides were washed twice with deionized water, and antigens were retrieved with antigen retriever Citrate Buffer (Sigma Aldrich). Non-specific binding was blocked with 2% Bovine Serum Albumin (BSA) dissolved in TBS with 0.2% Triton X-100. After blocking, primary antibodies were incubated in the blocking buffer at 4°C overnight. Primary antibodies in this study include 5 μg/mL of CD8a monoclonal antibody (clone AMC908, Thermo Fisher Scientific, Alexa Flour 488-conjugated), 5 μg/mL of CD19 monoclonal antibody (clone 6OMP31, Thermo Fisher Scientific), 15 μg/mL of CXCR1 monoclonal antibody (clone 42705.111, Thermo Fisher Scientific, Alexa Fluor 594-conjugated), or 15 μg/mL of IL-8 monoclonal antibody (clone 6217, Thermo Fisher Scientific) in separate staining panels to avoid cross-reactivity. Slides were washed and stained with secondary antibodies for CD19 and IL-8, including anti-Mouse IgG (H+L) secondary antibody-Alexa Fluor 594, or anti-Rat IgG (H+L) secondary antibody-Alexa Fluor 488 (Thermo Fisher Scientific) for 1 hr at RT. After 1 hr, slides were washed and mounted with Vectashield mounting media with DAPI (Vector Laboratories). Images were obtained with Nikon confocal microscopy and analyzed with IMARIS software (Oxford Instruments).

### Western blot

To detect citrullinated proteins in NETs, 2 × 10^6^ neutrophils/well were plated in 12 well plate, and equal volumes of supernatants were collected after stimulation, and denatured with SDS sample buffer containing β-mercaptoethanol (Bio-Rad). Protein citrullination was detected by human ACPA immunoblotting. Briefly, transferred PVDF membrane was incubated with 1 μg/mL of purified human ACPA in TBST at 4°C overnight, and anti-human IgG HRP (Cell Signaling, 1:10,000 dilution) for 1 hr RT. SuperSignal West Pico PLUS chemiluminescent (Thermo Fisher Scientific) was used for chemiluminescence detection.

### Secreted embryonic alkaline phosphatase (SEAP) production by HEK-Blue hTLR cells

HEK-Blue^TM^ hTLR2 or hTLR4 cells were purchased from InvivoGen. The cell line was cultured in Dulbecco’s minimal essential media (DMEM) in the presence of L-glutamine, 10% heat-inactivated fetal bovine serum (FBS), 100 U/mL penicillin/streptomycin, and 100 μg/mL normocin in 75 cm^2^ flasks at 37°C in 5% CO_2_ incubator. The cell lines were used only under 10 passages following thawing. To stimulate the cells, 1.5 × 10^5^ cells/mL were seeded onto 96 flat-bottom wells, and treated with NET supernatants, native or citrullinated H3 or vimentin in a dose dependent manner for 24 hr at 37°C. LPS (Sigma Aldrich, 0.1 μg/mL) or FSL-1 (Pam2CGDPKHPKSF) (InvivoGen, 50 ng/mL) was used as a positive control of HEK-Blue TLR4 or TLR2 stimulation. SEAP activity was measured by HEK-Blue Detection at OD_650_ with BioTek synergy H1 microplate reader (Agilent).

### T cell proliferation assay

For the isolation and maturation of MoDCs, CD14^+^ monocytes were isolated using Human CD14 Positive Selection kit II (Stem Cell Technologies), and differentiated into immature dendritic cells with Mo-DC Differentiation Medium containing GM-CSF and IL-4 (Miltenyi Biotec) for 7 days. On day 7, human TNF alpha in fresh Differentiation Medium was added to generate mature MoDCs for an additional 3 days. Mature MoDCs were incubated with NET supernatants, 5 μg/mL of native histone H3 or vimentin (Cayman chemical), or 5 μg/mL of citrullinated histone H3 or vimentin (Cayman Chemical) for 12 hr. Small molecules, including TAK-242 TLR4 inhibitor (EMD Millipore, 5 μM), Zimlovisertib IRAK4 inhibitor (Sigma Aldrich, 5 μM) or Ibrutinib BTK inhibitor (Selleckchem, 1 μM), were pre-treated 30 min before the addition of each stimulation to block TLR or BCR signaling. Protein processing or trafficking inhibitors, including 5 μg/mL of Brefeldin A (BFA, Thermo Fisher Scientific), 10 μg/mL of primaquine (Sigma Aldrich), 10 μM of chloroquine (InvivoGen), or 10 μg/mL of lactacystin (Sigma Aldrich) were also pre-treated 30 min prior to the stimulation of citrullinated proteins to block the cross-presentation pathway. Inhibitor concentrations were determined based on dose-response titration experiments to ensure cell viability remained comparable to vehicle controls. Cell viability was assessed using Fixable Viability Stain 510. Autologous CD8^+^ T cells from the same patients were isolated using EasySep Human CD8^+^ T cell Isolation kit (Stem Cell Technologies), and labeled with eBioscience Cell Proliferation Dye eFluor 450 (Thermo Fisher Scientific). For the coculture, the antigen-loaded MoDCs were incubated with the labeled CD8^+^ T cells at 1:10 ratio in RPMI 1640 media containing recombinant IL-2 (Peprotech, 50 U/mL) and anti-CD28 antibody (BD Biosciences, 2 μg/mL) at 37°C for 7 days. To block the interaction of HLA class I complex and T cell receptor, anti-HLA-A,B,C antibody (BioLegend, 1 μg/mL) was added with each antigen. Fresh culture media was replenished every 3 days.

For B cell and CD8^+^ T cell coculture, pan CD19^+^ B cells and CD8^+^ T cells from ACPA^+^ RA PBMCs were isolated using EasySep Human Pan-B cell Enrichment kit and Human CD8^+^ T cell Isolation kit, separately. B cells were pre-treated with pharmacologic inhibitors, including 5 μM TLR4 inhibitor, 5 μM E6446 TLR7 inhibitor (Sellekchem), 5 μM IRKA4 inhibitor, or 1 μM Evobrutinib BTK inhibitor (MedChemExpress) 30 min before stimulation of B cells. Then, B cells were incubated with NETs, native or citrullinated proteins for 3 hr, and cocultured with Cell Proliferation Dye eFluor 450-labeled CD8^+^ T cells in Iscove’s Modified Dulbecco’s Medium (IMDM) containing anti-CD28 antibody, IL-2, IL-21 (Peprotech, 50 ng/mL), and recombinant BAFF protein (R&D Systems, 100 ng/mL) for 7 days. The ration of B cells and CD8^+^ T cells was 1:10. For ACPA^+^ or ACPA^−^ B cells and CD8^+^ T cells coculture, the cells were sorted as previously described, and cocultured in the same condition with pan B cell and CD8^+^ T cell coculture assay. After 7 days, proliferating or activated T or B cells were quantified using flow cytometry. The proliferating cells were calculated based on the proportion of Cell Proliferation Dye^low^ population among total cells.

### Citrullinated protein internalization with ACPA immune complex

Human ACPAs were purified as previously described (*88*). To generate ACPA immune complex with citrullinated proteins, 2.5 μg/mL of His-tagged citrullinated histone H3 and vimentin (Cayman chemical) was separately incubated with 25 μg/ml of 2 different types of ACPAs (ACPA 66, ACPA74), or isotype control at 37°C for 1 hr. 10 μg/ml of anti-Fc gamma receptor (FcγR) antibody (BD Biosciences) was pre-treated to MoDCs 30 min prior to the treatment of ACPA immune complex. After 4 hr of ACPA immune complex treatment, internalized His-tag in MoDCs was detected using flow cytometry.

### shRNA-mediated *TLR4* knockdown

Control or *TLR4* shRNA lentiviral particles, containing a pool of four *TLR4*-specific constructs to knockdown gene expression, were purchased from Santa Cruz Biotechnology. MoDCs were differentiated from CD14^+^ monocytes in 24 well plates, and transfected with control or *TLR4* shRNA lentiviral particles according to the manufacturer’s instruction. After 48 hr of post-transfection, TLR4 expression was assessed using flow cytometry. The shRNA-treated MoDCs were cocultured with autologous CD8^+^ T cells from the same patient.

### Generation of fluorophore-conjugated citrullinated peptide tetramers

The list of biotinylated citrullinated or native peptides is provided in table S1. These citrullinated peptides were validated as a strong binding to ACPAs in previous studies (*89, 90*). The citrullinated peptides were incubated with phycoerythrin (PE) or allophycocyanin (APC)-labeled streptavidin (Thermo Fisher Scientific), and native peptides were conjugated with Brilliant Violet 605 (BV605), as previously described (*91*). Optimal concentrations of the labeled tetramers were determined by titration with B cells from CCP-high RA patients (fig. S10). In this study, citrullinated peptide-PE and APC double-positive and BV605-negative population was used as ACPA^+^ B cells, and PE and APC double-negative population was used as ACPA^−^ B cells.

### Quantification of secreted ACPA level

To quantify the secreted level of ACPA, B cells were isolated from ACPA^+^ RA PBMCs, and stimulated with anti-IgM antibody (Jackson Immunoresearch), native or citrullinated proteins, or NETs for 10 days. The supernatants were collected and ACPA level was measured using Human Cyclical Citrullinated Peptide ELISA kit (MyBioSource).

### Transwell assay

Pan B cells were isolated from ACPA^+^ RA PBMCs, and seeded onto the bottom well of Transwell plate with 96 permeable supports with 0.4 or 5.0 μm pore polycarbonate membrane (Corning). B cells were pre-stimulated with anti-IgM antibody, native or citrullinated proteins for 1 day and autologous CD8^+^ T cells were seeded onto the Transwell insert with the ratio of 1:10 (B:CD8^+^) in complete RPMI 1640 medium containing IL-2. The cells were incubated for 12 hr in the presence or absence of pre-treated 1 μg/mL of anti-human IL-8 antibody (Thermo Fisher Scientific) and/or anti-human CXCR1 antibody (Thermo Fisher Scientific). The supernatant from B cell culture stimulated by anti-IgM antibody, native or citrullinated proteins for 7 days was added to investigate whether CD8^+^ T cells on the transwell insert migrate into the plate due to chemotaxis. The activation of CD8^+^ T cells in the plate bottom (migrated) or transwell insert (non-migrated), or CD19^+^ B cells in the plate bottom was measured using flow cytometry. The migrated cell ratio was calculated based on migrated cell counts in untreated wells.

### Quantification of chemokine levels

Pan B cells were isolated from ACPA^+^ RA PBMCs and stimulated with anti-IgM antibody, native or citrullinated proteins in IMDM containing IL-2, IL-21, and BAFF for 7 days. The cell culture supernatants were collected and stored at −80°C. Samples were sent to Eve Technologies and the level of chemokine was measured using the Human Cytokine/Chemokine 96-Plex Discovery Assay based on the Luminex 200 platform (Luminex). The assay was performed according to the manufacturer’s protocol. Using the Human Chemokine Antibody Array with 38 targets (Abcam), we semi-quantitatively measured chemokine levels from the cell culture supernatants following with the manufacturer’s instruction, and quantified the level of each chemokine on the membrane using Image J software.

### Statistical analysis

GraphPad Prism v10 or R v4.41 were used for the statistical analyses. Data are represented as means ± SEM. Specific statistical tests for each figure are indicated in the respective figure legends.

## Supplementary Materials

Figs. S1 to S20

Tables S1 to S6

## Acknowledgments

We thank R. Camille Brewer for providing bacterial lysates and protocols, Alejandro Gomez for providing recombinant ACPA mAbs, and Tobias Lanz for providing inputs for tetramer generation and insightful discussions. We also thank all the volunteers and patients in this study at Stanford hospital and VA Palo Alto healthcare system.

## Funding

The study was supported by following grants and funding sources:

National Institutes of Health grant R01 AR063676 (WHR)

National Institutes of Health grant U19 AI110491 (WHR)

National Institutes of Health grant R01 AR078268 (WHR)

Johnson and Johnson

## Author contributions

Conceptualization: JSM, WHR

Methodology: JSM, MZ, LSD, EKS, WHR

Investigation: JSM, MZ, WHR

Resources: OS, JC, DCR, MHS, LTD

Visualization: JSM, MZ

Funding acquisition: WHR

Project administration: JSM, OS, WHR

Supervision: JSM, MMD, WHR

Writing – original draft: JSM, MZ, MCH, WHR

Writing – review & editing: JSM, MZ, LSD, EKS, OS, JC, DCR, MHS, LTD, MCH, MMD, WHR

## Data and materials availability

The files of sequencing data will be deposited to SRA upon acceptance of the current work. All reagents and materials used in this study are commercially available.

