## Supplementary Materials for "Cross-presentation of citrullinated antigens drives cytotoxic CD8^+^ T cell responses in rheumatoid arthritis"

Figs. S1 to S20

Tables S1 to S6

16 **Supplementary Figure**

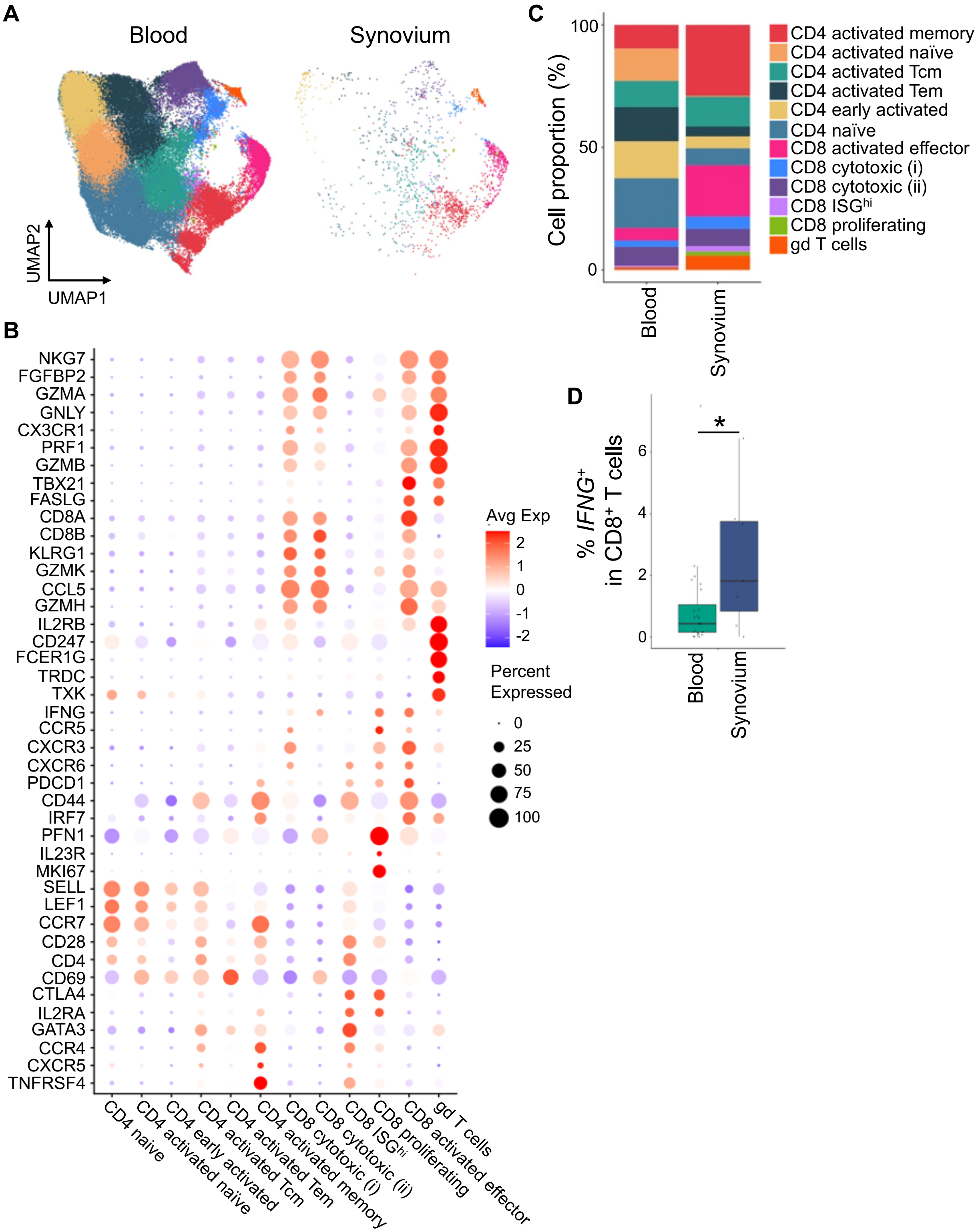


17

**Figure S1. Transcriptomic characterization of RA blood and synovial T cells.** (**A**) UMAP plots of T cells from RA blood and synovium shown separately (*n* = 93,086 cells for blood; *n* = 1,817 cells for synovium). (**B**) Heatmap showing expression of canonical marker genes used to annotate major T cell populations. (**C**) Proportion of annotated T cell clusters in RA blood and synovium. (**D**) Percentage of $IFNG^{+}$ $CD8^{+}$ T cells in RA blood and synovium, defined as cells within $CD8^{+}$ T cell clusters with *IFNG* > 0. Statistical analysis in cell percentage was calculated with sample level Bayes linear model in Limma. *$P < 0.05$

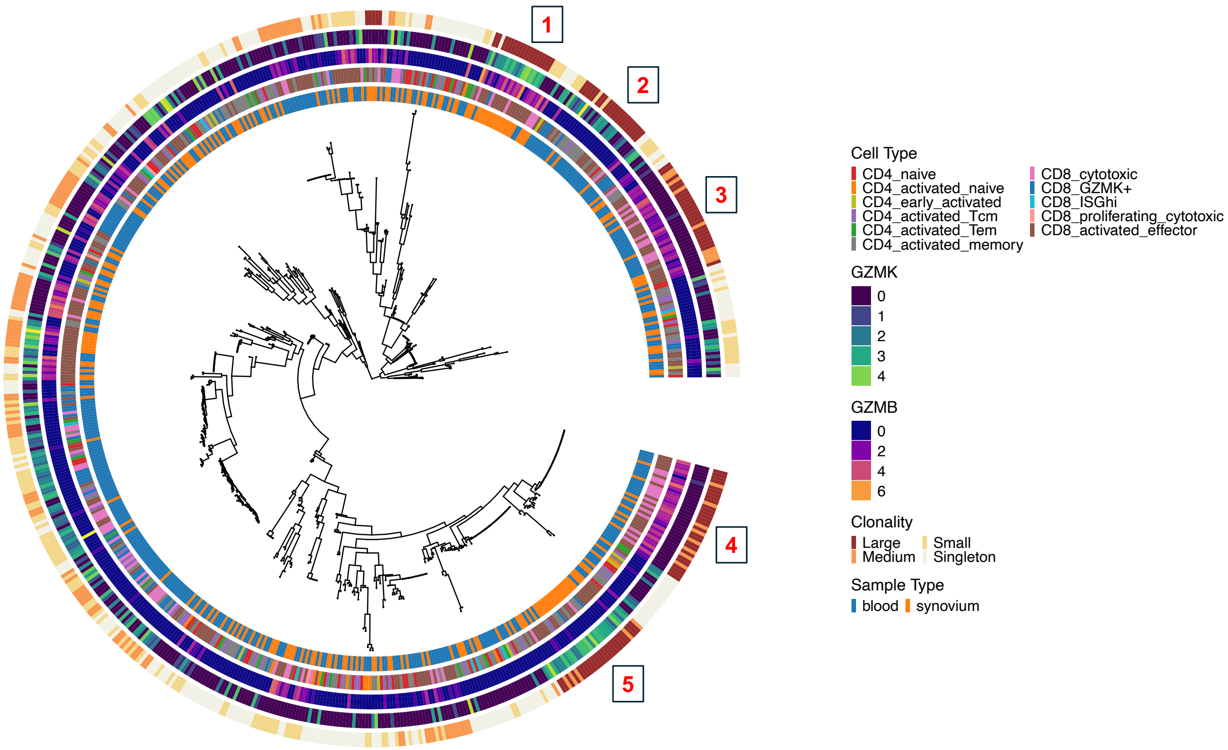


**Figure S2. TCR lineage tree of RA T cells highlights expanded clone families and their transcriptional states.** TCR lineage tree constructed from paired TCRαβ sequences obtained from RA blood and synovial T cell scTCR data ($n$ = 646 in 5 patients). Each node represents a single T cell, and edges connect clonotypes with shared sequence similarity. The tree is annotated by (i) cell cluster, (ii) *GZMK* expression level, (iii) *GZMB* expression level, (iv) TCR clonality/clone size, and (v) sample type. Largely expanded clone families are highlighted and labeled 1–5.

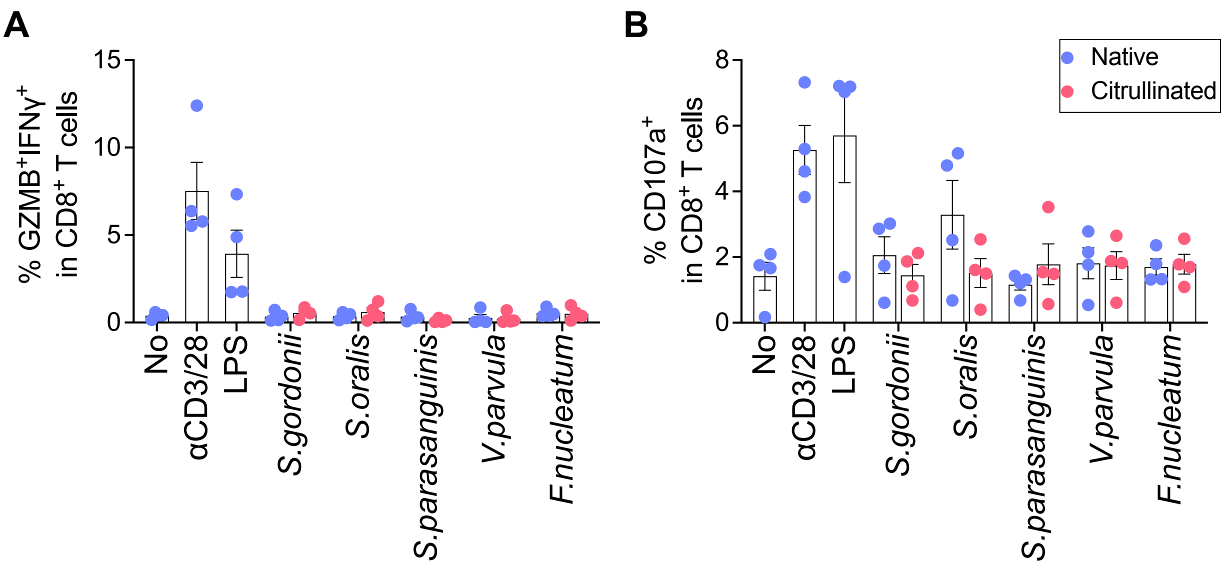


**Figure S3. Healthy control CD8+ T cells do not respond to citrullinated bacterial lysates. (A and B**) Proportion of GZMB$^{+}$IFNγ$^{+}$ (A) or CD107a$^{+}$ (B) CD8$^{+}$ T cells from healthy control PBMCs ($n$ = 4) stimulated by native (*blue*) or citrullinated (*red*) oral bacterial lysates for 16 hr. SG, *Streptococcus gordonii;* SO; *Streptococcus oralis;* SP; *Streptococcus parasanguinis;* VP, *Veillonella parvula;* FN, *Fusobacterium nucleatum.* Statistical analysis was determined using two-way ANOVA with Tukey's test.

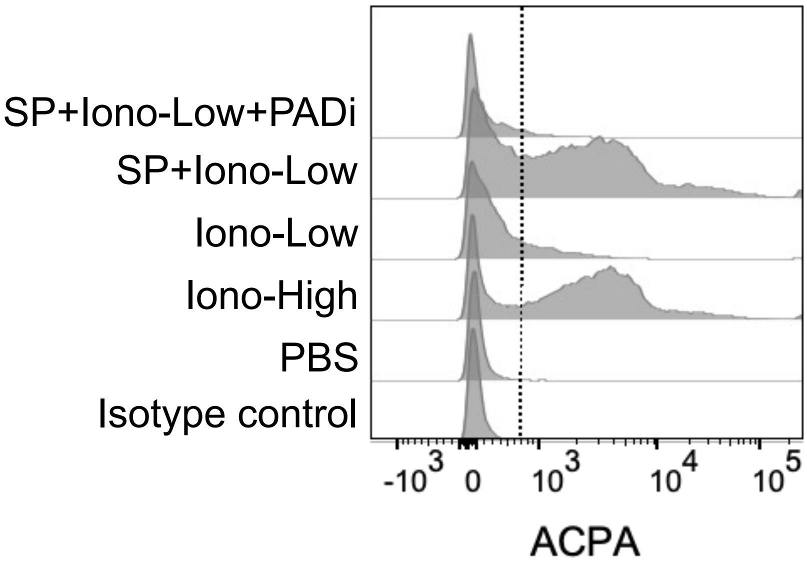


**Figure S4. ACPA binds to *S.parasanguinis*-mediated NETs-derived antigens.** Flow cytometry histrogram showing ACPA binding ability to NETs induced by ionomycin high concentration (1 μM, Iono-High), ionomycin low concentration (0.01 μM, Iono-Low), native SP lysates with low-dose ionomycin (SP+Iono-Low), or SP with low-dose ionomycin and PAD inhibitor (Cl-amidine) (SP+Iono-Low+PADi). SP; *S. parasanguinis*, PADi; PAD enzyme inhibitor.

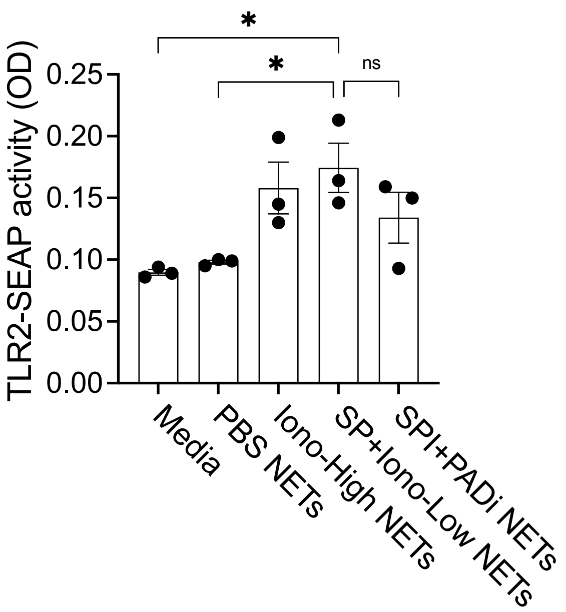


**Figure S5. TLR2 signaling is activated by oral bacterial NET components.** Quantification of TLR2-SEAP activity induced by treatment of media only, or PBS, ionomycin high, native *S. parasanguinis* (SP) lysates with low-dose ionomycin (SPI), or SPI with PAD inhibitor-induced NETs. n = 3. Statistical analysis was determined using one-way ANOVA with Tukey's multiple comparisons test. Data are plotted as means ± SEM. *P < 0.05.

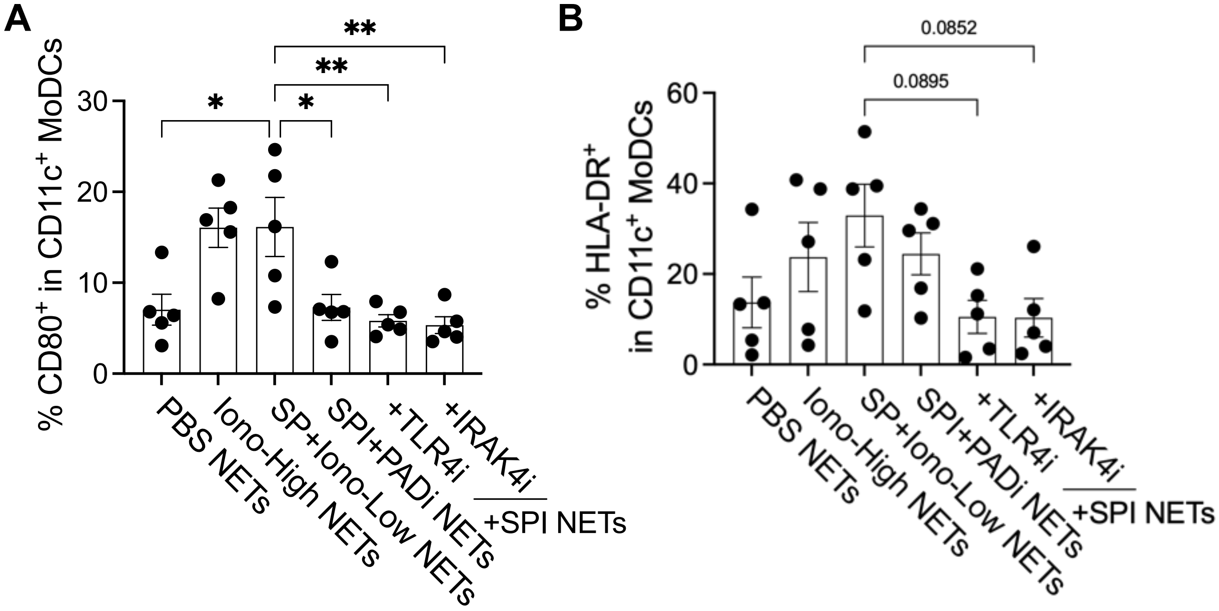


**Figure S6. *S. parasanguinis*-induced NETs stimulate dendritic cells to exhibit antigen-presenting activity.** (**A** and **B**) Quantification of CD80$^{+}$ (A, n = 5) or HLA-DR$^{+}$ (B, n = 5) on the surface of CD11c$^{+}$ MoDCs in response to PBS, high-dose ionomycin, native *S. parasanguinis* (SP) lysates with low-dose ionomycin (SPI), or SPI with PAD inhibitor-induced NETs in the presence or absence of TLR4, or IRAK4 inhibitor. Statistical analysis was determined using one-way ANOVA with Tukey's multiple comparisons test (A and B). Data are plotted as means ± SEM. P values are presented on the bar graph (B). *$P < 0.05$.

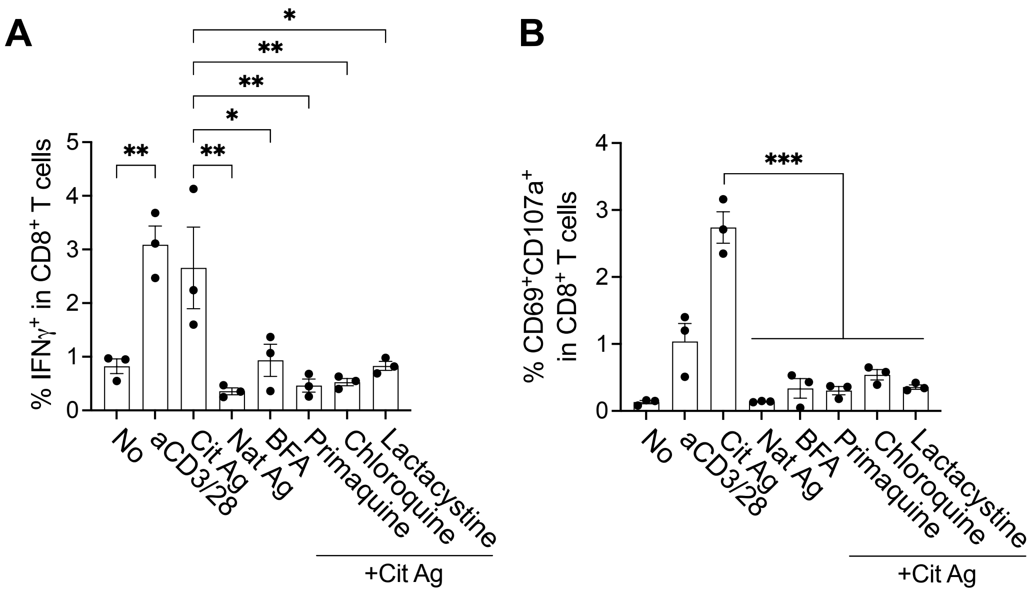


**Figure S7. Inhibitors of cross-priming decrease activation of CD8$^+$ T cells in response to citrullinated antigens.** (**A** and **B**) Monocyte-derived dendritic cells were pre-treated with cross-priming inhibitors for 30 min, and incubated with CD8+ T cells from the same donor in the presence or absence of native or citrullinated proteins for 7 days. Anti-CD3 antibody was used as a positive control for CD8$^+$ T cell activation. IFNγ$^+$ (A) or CD69$^+$CD107a$^+$ (B) CD8$^+$ T cells were quantified using flow cytometry. n = 3. Statistical analysis was determined using one-way ANOVA with Tukey's multiple comparisons test (A and B). Data are plotted as means ± SEM. *P < 0.05. **P < 0.01, ***P < 0.001.

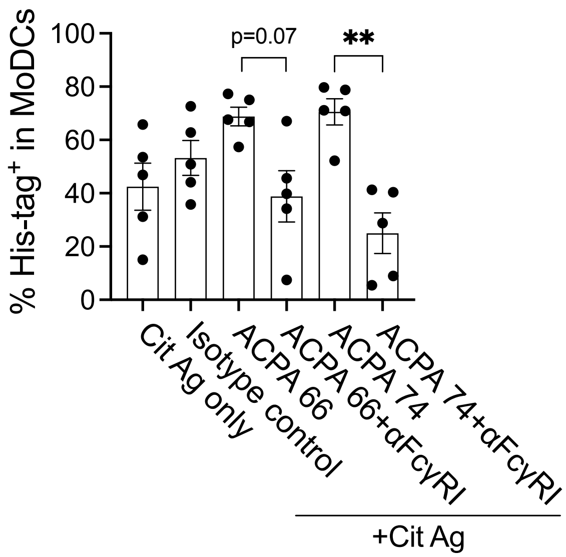


**Figure S8. ACPA immune complexes facilitate dendritic cell internalization of citrullinated antigens.** Bar graph representing the percentage of internalized His-tag$^{+}$ monocyte-derived dendritic cells ($n$ = 5). MoDCs were incubated with His-tagged citrullinated antigen only, or immune complexed with isotype control IgG, or two different types of ACPAs (ACPA 66, or 74) in the presence or absence of Fc receptor blocking antibody for 30 min. Internalized His-tag was detected by flow cytometry.

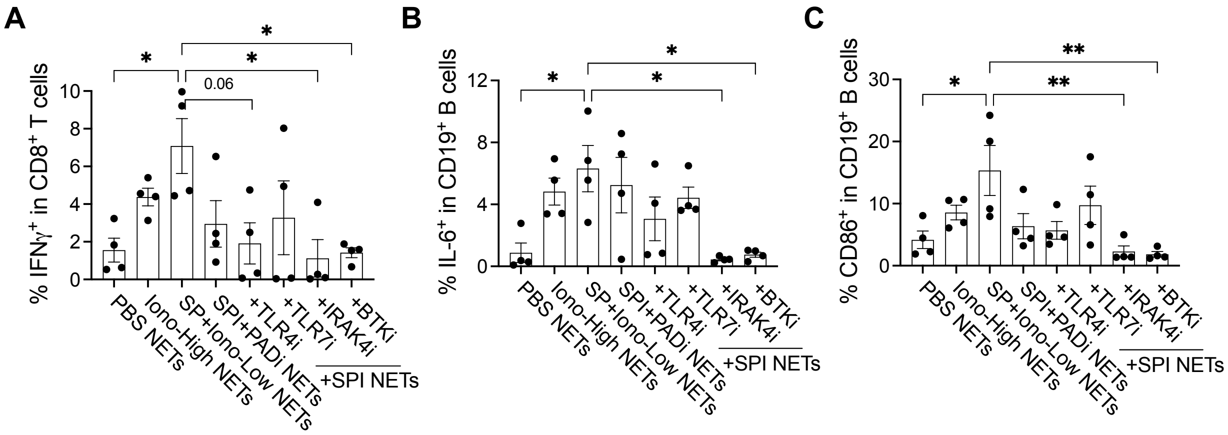


**Figure S9. *S. parasanguinis*-induced NETs activate B cell responses.** (**A** to **C**) $CD8^+$ T cells were co-cultured with pan B cells from the same $ACPA^+$ RA patient, treated with PBS, high-dose ionomycin, native *S. parasanguinis* (SP) with low-dose of ionomycin (SPI), SPI with PAD inhibitor-induced NETs in the presence or absence of TLR4, TLR7, IRAK4, or BTK inhibitor. $IFN\gamma^+$ $CD8^+$ T cells (A, $n = 4$), IL-$6^+$ (B, $n = 4$), or $CD86^+$ (C, $n = 4$) $CD19^+$ B cells were measured from the coculture by flow cytometry. Statistical analysis was determined using one-way ANOVA with Tukey's multiple comparisons test (A-C). Data are plotted as means ± SEM. $^*P < 0.05$.

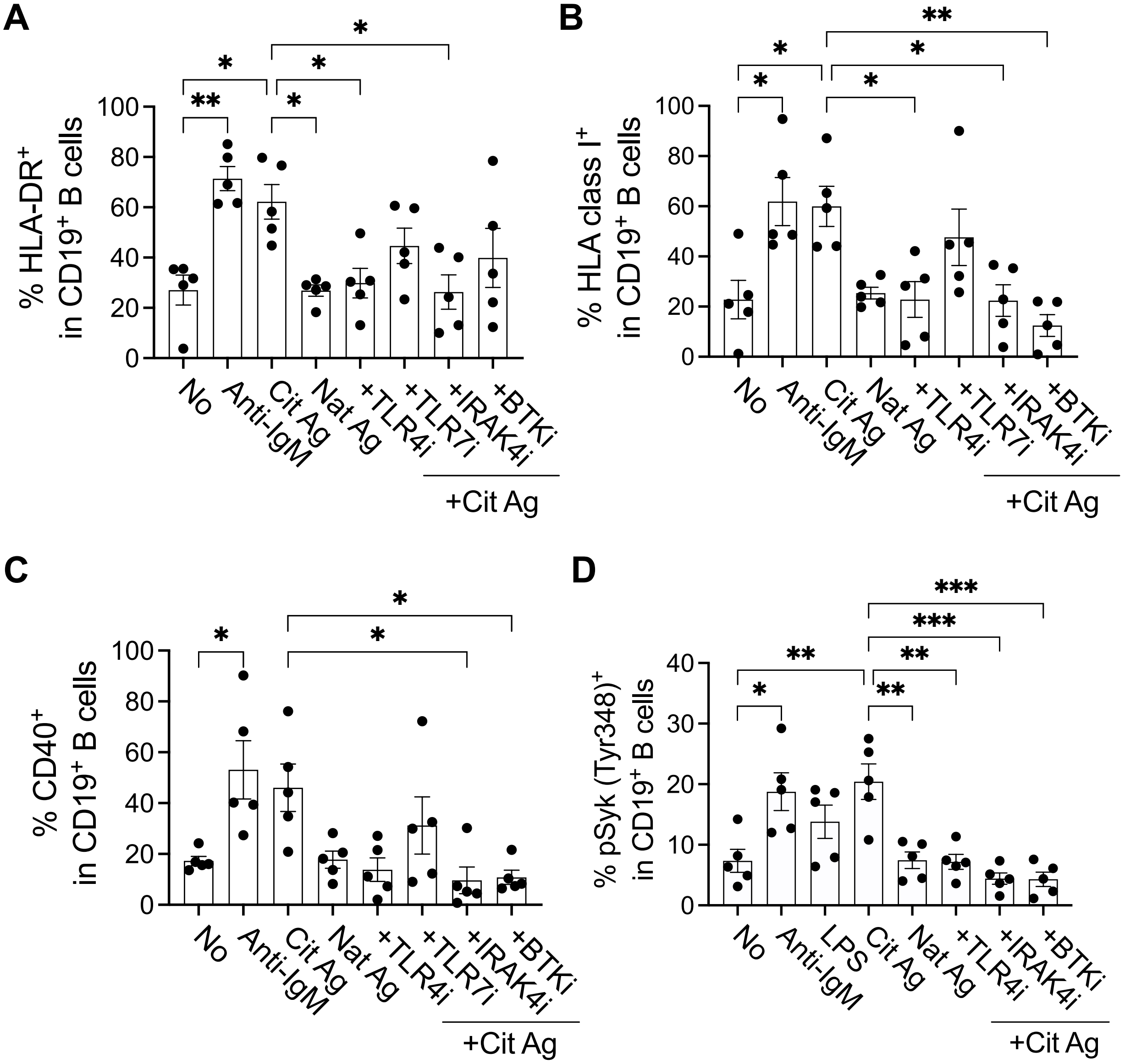


**Figure S10. Citrullinated antigens stimulate B cells in a TLR4-dependent manner to express antigen-presenting markers of B cells.** (**A** to **C**) Pan B cells were incubated with anti-IgM antibody, native (Nat) or citrullinated (Cit) antigens with or without TLR4, TLR7, or IRAK4 inhibitor for 24 hr. HLA-DR$^+$ (A, $n$ = 5), HLA-A, B, C$^+$ (B, $n$ = 5), or CD40$^+$ (C, $n$ = 5) CD19$^+$ B cells were quantified using flow cytometry. (**D**) Quantification of phospho-Syk (Tyr348)-expressing CD19$^+$ B cells in ACPA$^+$ RA patients in response to anti-IgM antibody, LPS, native or citrullinated antigens in the presence or absence of TLR4, IRAK4, or BTK inhibitor. $n$ = 5. Statistical analysis was determined using one-way ANOVA with Tukey's multiple comparisons test (A-D). Data are plotted as means ± SEM. *$P < 0.05$. **$P < 0.01$, ***$P < 0.001$.

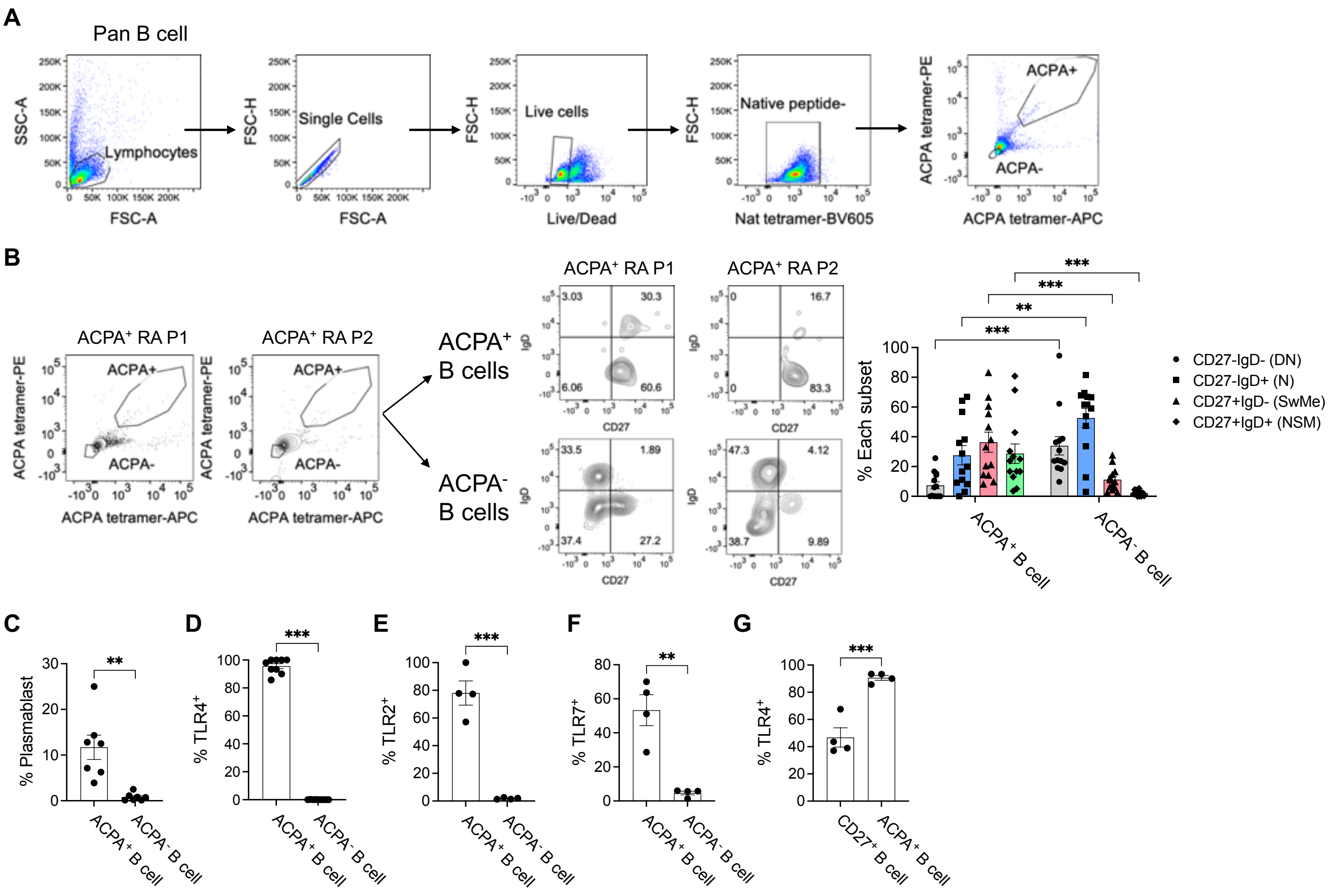


**Figure S11. Phenotype of ACPA$^+$ B cells.** (**A**) Gating strategy of ACPA-tetramer$^+$ or ACPA$^-$ B cells in total B cells from ACPA$^+$ RA blood. (**B**) Representative flow cytometry dot plots (*left*), or frequencies (*right*) of each B cell subset in ACPA-expressing or non-expressing B cells from 12 ACPA$^+$ RA patients. DN; double negative, N; naïve, SwMe; switched memory, NSM; non-switched memory. (**C**) Quantification of plasmablast ($n$ = 7) among ACPA$^+$ or ACPA$^-$ B cells. (**D** to **F**) Quantification of TLR4$^+$ B cells (D, $n$ = 9), TLR2$^+$ B cells (E, $n$ = 4), or TLR7$^+$ B cells (F, $n$ = 4) among ACPA$^+$ or ACPA$^-$ B cells. (**G**) Comparison of TLR4$^+$ B cells among CD27$^+$ or ACPA$^+$ B cells in four ACPA$^+$ RA bloods. Statistical analysis was determined using two-way ANOVA with Fisher's LSD test (B), or Student's unpaired t-test (C-G). Data are plotted as means ± SEM. $^{**}P < 0.01$, $^{***}P < 0.001$.

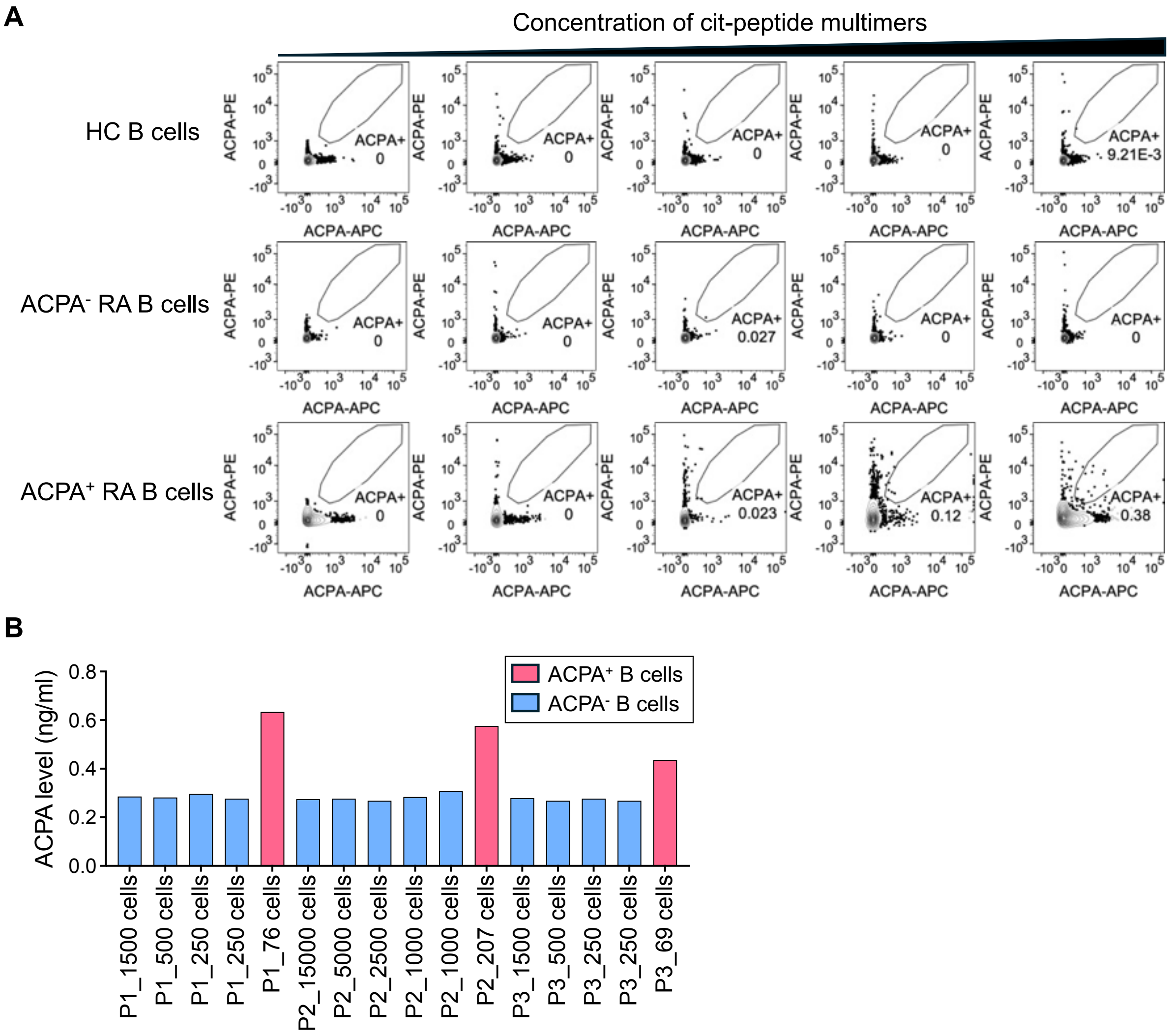


**Figure S12. Target specificity of citrullinated peptide-tetramers.** (**A**) Pan B cells from healthy control, ACPA$^-$ RA, or ACPA$^+$ RA patients were incubated with various concentration of citrullinated peptide-tetramers and detected APC and PE-double positive population as ACPA$^+$ B cells. (**B**) Different number of ACPA$^-$ (*blue*) or ACPA$^+$ (*red*) B cells from ACPA$^+$ RA blood (Patient 1-3) were sorted using staining of citrullinated peptide-tetramers, and cultured for 10 days. The level of secreted ACPA in culture supernatants from ACPA$^-$ or ACPA$^+$ B cells was measured using ACPA ELISA ($n$ = 3).

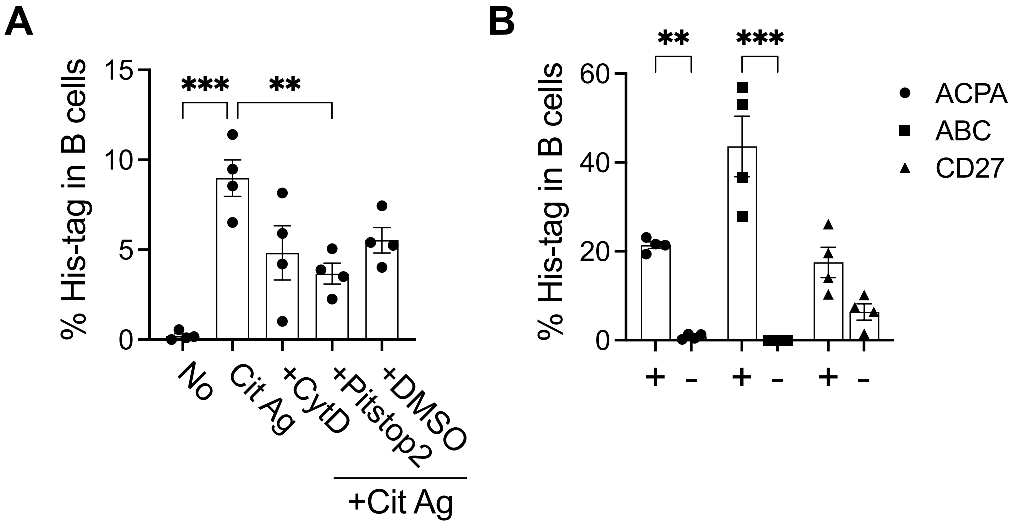


**Figure S13. Citrullinated antigens can be internalized by receptor-mediated endocytosis.** (**A** and **B**) Pan B cells were pre-treated with cytochalasin D (CytD), pitstop 2, or DMSO for 30 min, and incubated with His-tagged citrullinated proteins for 1 hr. Cells were fixed and permeabilized to detect internalized His-tagged protein in B cells. Expression of His-tag was measured in total B cells (A, $n$ = 4) or ACPA$^{+/-}$, age-associated B cells (ABC), or CD27$^{+/-}$ B cells (B, $n$ = 4). Statistical analysis was determined using one-way ANOVA with Tukey's multiple comparisons test (A), or two-way ANOVA with Tukey's multiple comparisons test (B). Data are plotted as means ± SEM. **P < 0.01, ***P < 0.001.

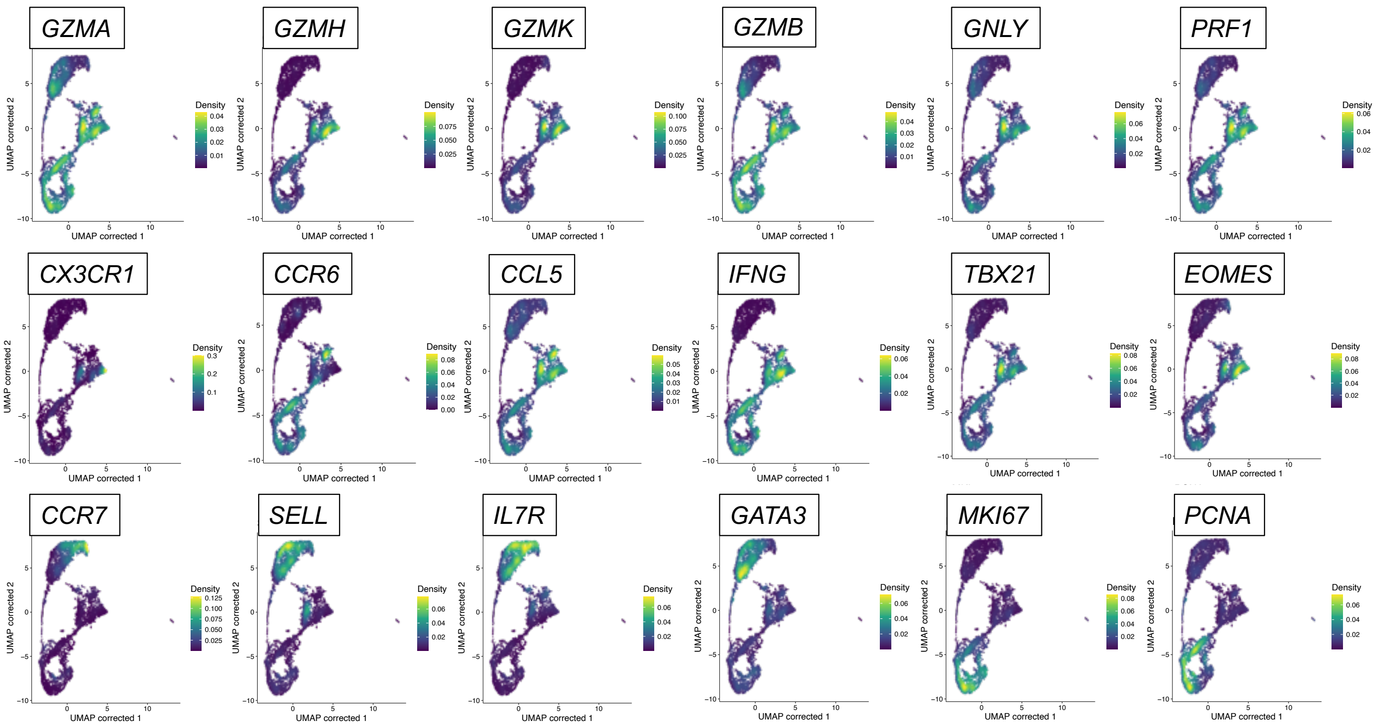


**Figure S14. Density plots of key markers of CD8$^+$ T cells.** Density plots with the levels of canonical markers to define CD8$^+$ T cell clusters.

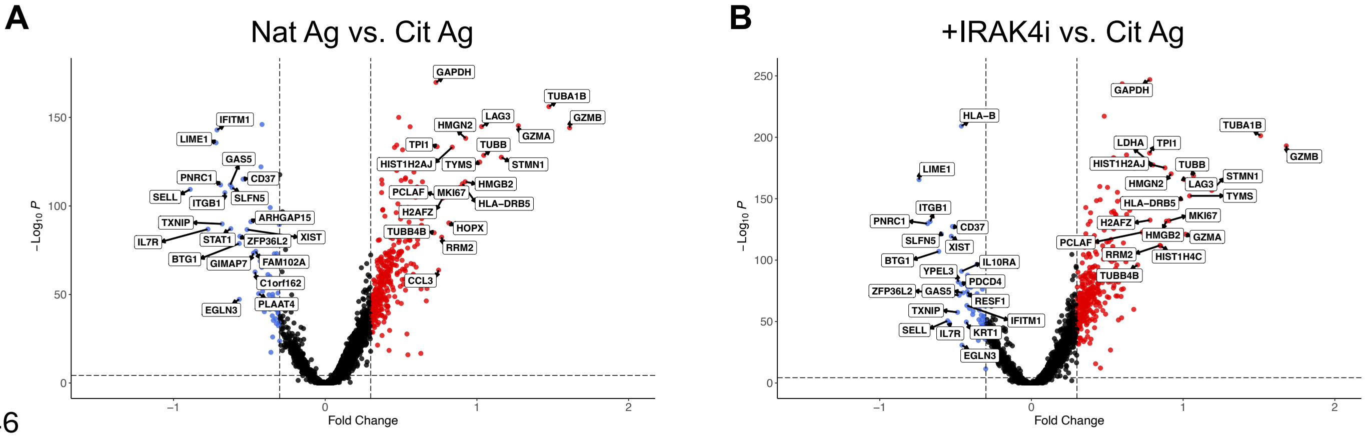


**Figure S15. Volcano plots of differentially expressed genes from scRNA-seq analysis of CD8$^+$ T and B cells.** (**A** and **B**) Volcano plots illustrating differentially expressed gene analysis contrasting native protein-treated CD8$^+$ T cells with citrullinated antigen-treated CD8$^+$ T cells (A), or citrullinated antigen-treated CD8$^+$ T cells with IRAK4 inhibitor and citrullinated antigen-treated CD8$^+$ T cells (B) from B and CD8$^+$ T cell co-culture assay. Statistical analysis was calculated using Wilcoxon rank-sum test to identify differentially expressed genes (DEGs).

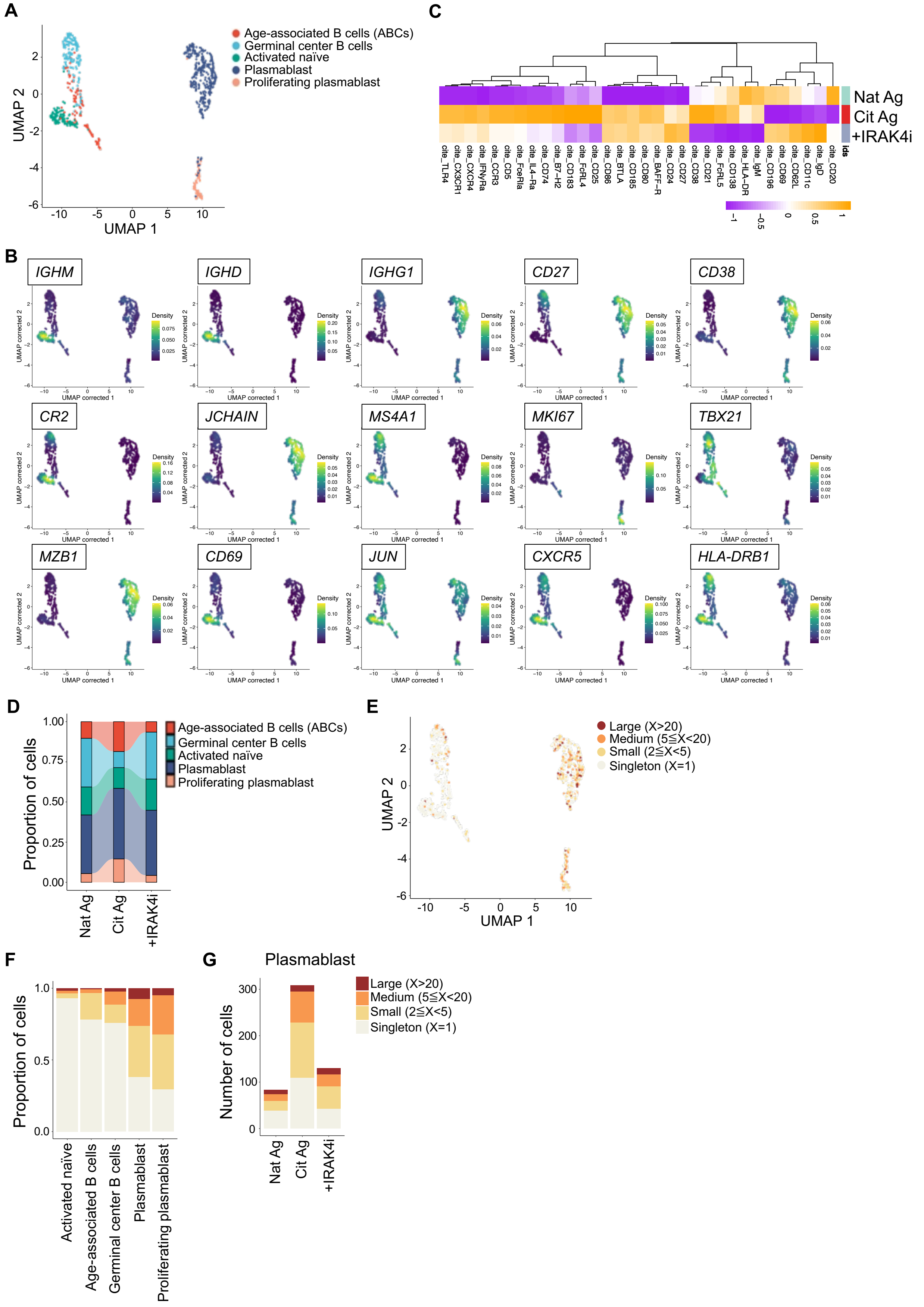
A
Age-associated B cells (ABCs)
Germinal center B cells
Activated naïve
Plasmablast
Proliferating plasmablast
UMAP 1
UMAP 2
C
Nat Ag
Cit Ag
+IRAK4i
ids
cite_TLR4
cite_CX3CR1
cite_CXCR4
cite_IFNyRa
cite_CCR3
cite_CD5
cite_FceRIa
cite_IL4−Ra
cite_CD74
cite_B7−H2
cite_CD183
cite_FcRL4
cite_CD25
cite_CD86
cite_BTLA
cite_CD185
cite_CD80
cite_BAFF−R
cite_CD24
cite_CD27
cite_CD38
cite_CD21
cite_FcRL5
cite_CD138
cite_HLA−DR
cite_IgM
cite_CD196
cite_CD69
cite_CD62L
cite_CD11c
cite_IgD
cite_CD20
B
IGHM
IGHD
IGHG1
CD27
CD38
CR2
JCHAIN
MS4A1
MKI67
TBX21
MZB1
CD69
JUN
CXCR5
HLA-DRB1
Density
UMAP corrected 1
UMAP corrected 2
D
Proportion of cells
Age-associated B cells (ABCs)
Germinal center B cells
Activated naïve
Plasmablast
Proliferating plasmablast
Nat Ag
Cit Ag
+IRAK4i
E
Large (X>20)
Medium (5≦X<20)
Small (2≦X<5)
Singleton (X=1)
UMAP 1
UMAP 2
F
Proportion of cells
Activated naïve
Age-associated B cells
Germinal center B cells
Plasmablast
Proliferating plasmablast
G
Plasmablast
Number of cells
Large (X>20)
Medium (5≦X<20)
Small (2≦X<5)
Singleton (X=1)
Nat Ag
Cit Ag
+IRAK4i

**Figure S16. Citrullinated antigens induce B cell differentiation and activation.** (**A**) UMAP plot of B cells (*n* = 1,330) in the coculture assay with B cells and $CD8^+$ T cells from 2 $ACPA^+$ RA PBMCs. (**B**) Density plots showing expression levels of canonical B cell markers to define clusters. (**C**) Heatmap showing protein expression level of key B cell markers in B cells from Cit Ag vs. Nat Ag vs. Cit Ag with IRAK4i stimulations based on the level of CITE-seq antibodies. (**D**) Alluvial plot representing comparison of proportion of each B cell cluster in each condition. (**E**) UMAP plot of B cells integrated with BCR clonality (*n* = 1,023 paired BCR sequences). Color indicates the groups by the frequency of clonotypes in total cells. X represents the frequency of each clonotype defined by its unique paired BCR sequence. Large-expanded clones (X>20), Medium-expanded clones (20>X ≧5), Small expanded clones (5>X≧2), and Singletons (X=1). (**F**) Proportion of clonal lineages based on clonal size in each cluster. (**G**) The absolute number of plasmablasts with BCR clonality in each condition.

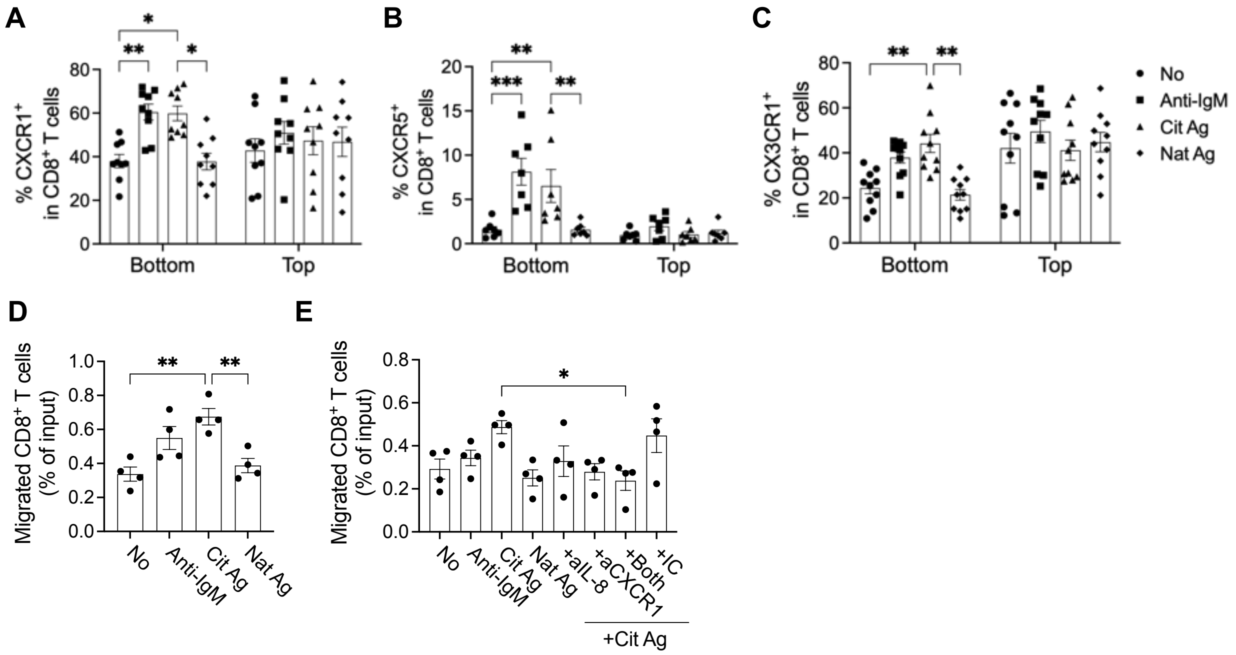


**Figure S17. Citrullinated antigen-stimulated B cells recruit inflammatory CD8$^+$ T cells.** (**A** to **C**) Quantification of CXCR1$^+$ (A, $n = 9$), CXCR5$^+$ (B, $n = 7$), or CX3CR1$^+$ (C, $n = 10$) CD8$^+$ T cells in the bottom or transwell insert of transwell assay. (**D**) Relative migration of CD8$^+$ T cells towards B cell supernatants. Data are presented as the fold change in migration relative to untreated B cell supernatants. $n = 5$. (**E**) Bar graph showing the percentage of migrated CD8$^+$ T cells (relative to total input) from the transwell insert into the bottom well containing B cells, in the presence or absence of indicated stimuli and blocking antibodies. $n = 4$. Statistical analysis was determined using two-way ANOVA with Tukey's multiple comparisons test (A-C) or one-way ANOVA with Tukey's multiple comparisons test (D, E). Data are plotted as means ± SEM. *$P < 0.05$. **$P < 0.01$, ***$P < 0.001$.

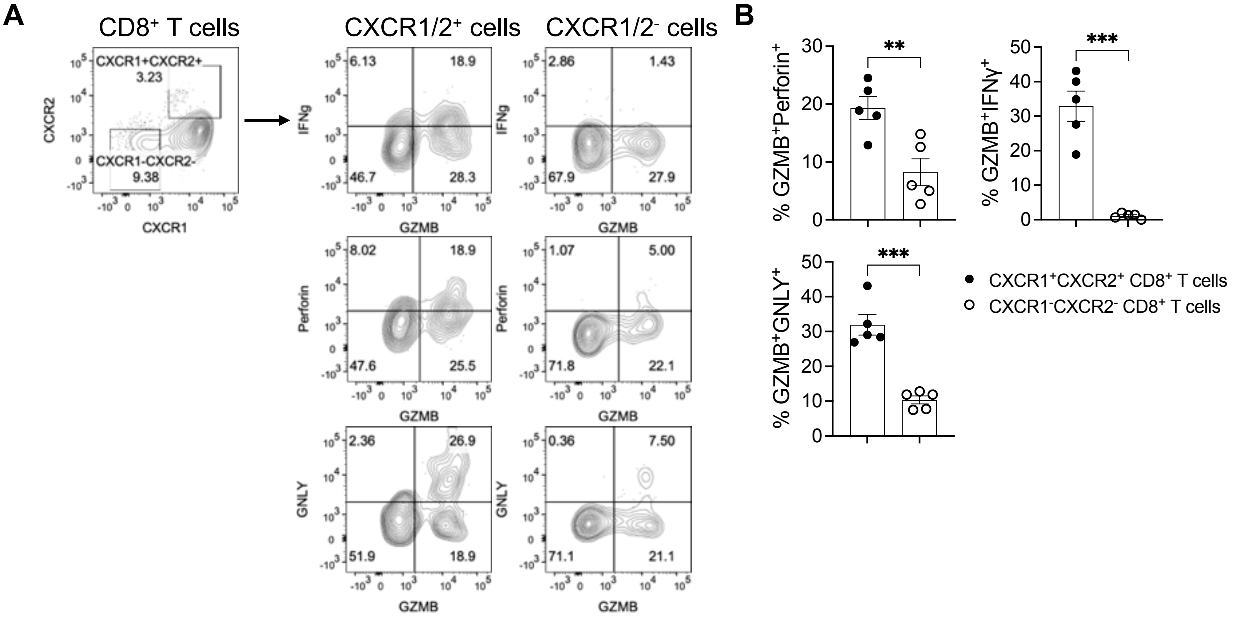


**Figure S18. CXCR1$^+$CXCR2$^+$ CD8$^+$ T cells are highly cytotoxic.** (**A**) Representative dot plots of the expressions of cytotoxic mediators comparing CXCR1/2$^+$ or CXCR1/2$^-$ CD8$^+$ T cells. (**B**) Quantification of GZMB$^+$IFNγ$^+$, GZMB$^+$Perforin$^+$, or GZMB$^+$GNLY$^+$ CD8$^+$ T cells in five ACPA$^+$ RA bloods. Statistical analysis was determined using Student's unpaired t-test (B). Data are plotted as means ± SEM. **P < 0.01, ***P < 0.001.

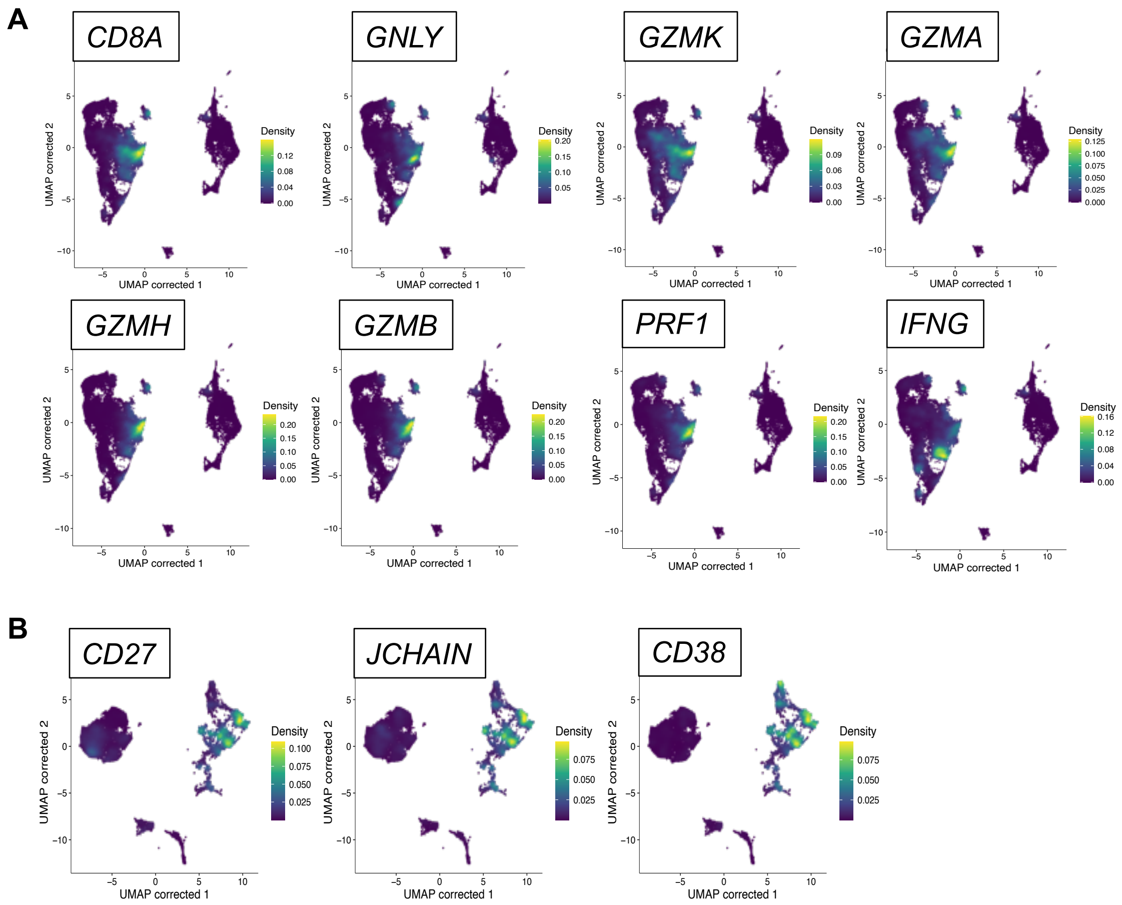


**Figure S19. Characterization of synovial CD8$^{+}$ T cells and B cells.** (**A**) Density plots with the levels of cytotoxic markers to define cytotoxic CD8$^{+}$ T cells from ACPA$^{+}$ RA and OA synovium. (**B**) Density plots of key markers of plasmablasts in from ACPA$^{+}$ RA and OA synovium.

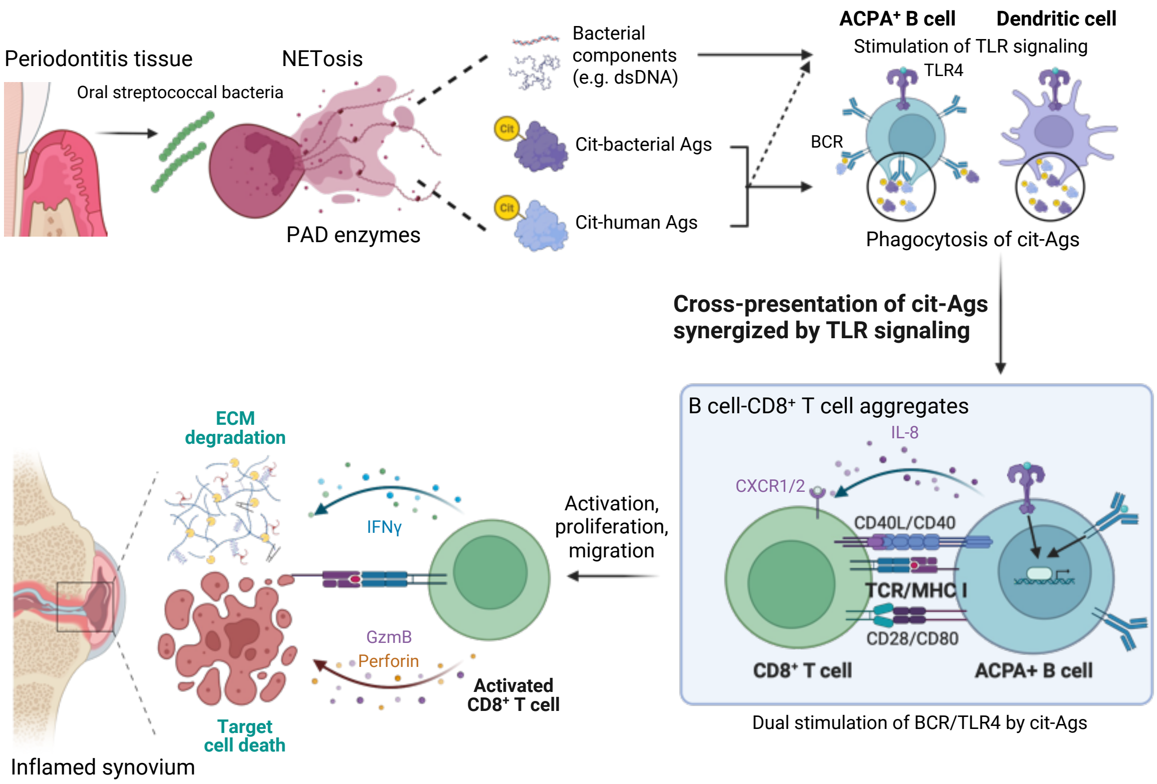


**Figure S20. Scheme of the proposed mechanism by which citrullinated antigens drive cytotoxic CD8$^{+}$ T cell activation through cross-presentation by MoDCs and ACPA$^{+}$ B cells to possibly mediate synovium inflammation and joint destruction in RA.** Figure created with BioRender. Cit; citrullinated, GzmB; Granzyme B, ECM; extracellular matrix, BCR; B cell receptor, TLR; Toll-like receptor.

**Supplementary Table**

| Peptide | Sequence |
|---|---|
| FibA 556-575 Cit sm Cyclic | Biotin-Ahx-NTKESSSHHPGCAEFPS[Cit]GKC-$CONH_2$ |
| FibA 616-635 Cit3 sm Cyclic | Biotin-Ahx-THSTK[Cit]CHAKS[Cit]PV[Cit]GIHTSC-$CONH_2$ |
| CFC 48-65 Cit2 Cyclic | Biotin-Ahx-CTIHAHPGSR[Cit]GG[Cit]HGYHHC-$CONH_2$ |
| Fibronectin 1029-1042 Cit2 | Biotin-Ahx-LTVGLT[Cit][Cit]GQPRQY |
| H2A/a-2 1-20 Cit3 | Biotin-Ahx-MSG[Cit]GKQGGKA[Cit]AKAKT[Cit]SS-$CONH_2$ |
| Clusterin 231-250 Cit3 | Biotin-Ahx-HFS[Cit]ASSIIDELFQD[Cit]FFT[Cit]-$CONH_2$ |
| H2A/a 1-20 Cit3 sm2 Cyclic | Biotin-Ahx-MSG[Cit]GKQGCKA[Cit]AKAKT[Cit]SSC-$CONH_2$ |
| Vim 58-77 Cit3 sm1 Cyclic | Biotin-Ahx-GGCVYAT[Cit]SSACV[Cit]L[Cit]SSVPGV-$CONH_2$ |

**Table S1. Peptide sequences of ACPA-tetramers.** Each peptide was conjugated with streptavidin-PE and APC to detect ACPA-expressing B cells. [Cit]; citrullination.

| Genes | logFC | pvalue | FDR |
|---|---|---|---|
| GZMB | 1.611231 | 1.63E-148 | 7.41E-145 |
| TUBA1B | 1.474714 | 4.47E-161 | 8.16E-157 |
| GZMA | 1.273434 | 6.60E-150 | 4.82E-146 |
| STMN1 | 1.160805 | 1.53E-131 | 3.10E-128 |
| TUBB | 1.045013 | 1.24E-132 | 2.66E-129 |
| LAG3 | 1.030657 | 2.68E-149 | 1.63E-145 |
| TYMS | 1.018758 | 8.84E-129 | 1.70E-125 |
| HMGN2 | 0.926125 | 1.72E-142 | 6.27E-139 |
| HMGB2 | 0.922488 | 1.84E-117 | 2.16E-114 |
| HLA-DRB5 | 0.904396 | 2.74E-116 | 3.03E-113 |
| MKI67 | 0.889119 | 2.82E-114 | 2.71E-111 |
| HIST1H2AJ | 0.837308 | 2.41E-137 | 6.27E-134 |
| HOPX | 0.815058 | 7.78E-94 | 4.11E-91 |
| H2AFZ | 0.774682 | 5.22E-108 | 3.74E-105 |
| RRM2 | 0.767928 | 1.03E-85 | 4.42E-83 |
| CCL3 | 0.747242 | 7.90E-67 | 1.70E-64 |
| TPI1 | 0.737546 | 1.20E-137 | 3.36E-134 |
| PCLAF | 0.730773 | 1.37E-109 | 1.02E-106 |
| GAPDH | 0.729891 | 4.55E-175 | 1.66E-170 |
| TUBB4B | 0.717837 | 3.20E-88 | 1.50E-85 |
| HIST1H4C | 0.670541 | 3.14E-56 | 4.62E-54 |
| CCL4 | 0.666309 | 3.83E-49 | 4.39E-47 |
| EBP | 0.641022 | 2.10E-117 | 2.39E-114 |
| LDHA | 0.635783 | 2.70E-96 | 1.54E-93 |
| ANXA2 | 0.635045 | 1.56E-92 | 7.92E-90 |
| LGALS1 | 0.631579 | 5.13E-56 | 7.48E-54 |
| H2AFV | 0.63142 | 1.24E-112 | 1.05E-109 |
| GNLY | 0.630414 | 5.68E-19 | 1.94E-17 |
| ENO1 | 0.614711 | 4.85E-113 | 4.42E-110 |
| NASP | 0.612705 | 8.85E-98 | 5.21E-95 |
| MCM7 | 0.608869 | 2.50E-84 | 1.04E-81 |
| TUBA1C | 0.608169 | 1.00E-81 | 3.66E-79 |
| SLC25A5 | 0.605588 | 2.03E-114 | 2.00E-111 |
| IL2RA | 0.599255 | 1.70E-68 | 3.84E-66 |
| PKM | 0.595468 | 1.81E-108 | 1.32E-105 |
| TXN | 0.592034 | 1.50E-110 | 1.17E-107 |
| CTSW | 0.589242 | 2.55E-80 | 9.13E-78 |
| EIF4A1 | 0.584743 | 4.71E-123 | 7.47E-120 |

| | | | |
|---|---|---|---|
| HIST1H1B | 0.5829 | 3.84E-70 | 9.41E-68 |
| PTTG1 | 0.582733 | 1.23E-88 | 5.84E-86 |
| CENPF | 0.580532 | 2.04E-64 | 4.03E-62 |
| UBE2S | 0.578163 | 6.61E-63 | 1.20E-60 |
| RAN | 0.57181 | 5.46E-113 | 4.86E-110 |
| CARHSP1 | 0.571734 | 2.97E-102 | 1.97E-99 |
| CXCR3 | 0.565145 | 1.46E-70 | 3.65E-68 |
| SNRPB | 0.562848 | 1.67E-117 | 2.03E-114 |
| HNRNPA2B1 | 0.559267 | 3.63E-149 | 1.89E-145 |
| FABP5 | 0.558316 | 6.36E-84 | 2.49E-81 |
| C12orf75 | 0.557723 | 4.58E-84 | 1.84E-81 |
| PGAM1 | 0.556454 | 1.76E-110 | 1.34E-107 |
| TESC | 0.55229 | 2.49E-94 | 1.34E-91 |
| LMNB1 | 0.549904 | 3.16E-84 | 1.30E-81 |
| ANP32E | 0.546109 | 7.46E-87 | 3.40E-84 |
| CCL5 | 0.545036 | 3.55E-18 | 1.16E-16 |
| PA2G4 | 0.544066 | 9.08E-103 | 6.25E-100 |
| UBALD2 | 0.543442 | 1.67E-79 | 5.93E-77 |
| NKG7 | 0.538601 | 3.55E-30 | 2.06E-28 |
| TNF | 0.536135 | 2.12E-69 | 5.03E-67 |
| MAD2L2 | 0.532143 | 6.72E-101 | 4.22E-98 |
| HMGB1 | 0.529482 | 9.16E-120 | 1.34E-116 |
| UBE2C | 0.529164 | 2.54E-55 | 3.62E-53 |
| PCNA | 0.527788 | 2.41E-66 | 5.11E-64 |
| DBI | 0.527659 | 6.04E-119 | 8.17E-116 |
| MCM5 | 0.527103 | 2.69E-83 | 1.02E-80 |
| H2AFX | 0.523704 | 2.23E-67 | 4.89E-65 |
| DUT | 0.522015 | 1.18E-61 | 2.07E-59 |
| TOP2A | 0.515223 | 1.08E-58 | 1.68E-56 |
| TPX2 | 0.509902 | 2.20E-67 | 4.87E-65 |
| ACTB | 0.509834 | 9.10E-136 | 2.07E-132 |
| SMC4 | 0.506955 | 2.23E-61 | 3.91E-59 |
| TMPO | 0.506435 | 1.90E-72 | 5.09E-70 |
| H2AFY | 0.505551 | 2.26E-90 | 1.11E-87 |
| RANBP1 | 0.503803 | 2.32E-77 | 7.62E-75 |
| TPM4 | 0.501158 | 1.13E-77 | 3.88E-75 |
| HIST1H1E | 0.501139 | 1.10E-47 | 1.21E-45 |
| EIF1AY | 0.500835 | 3.28E-118 | 4.13E-115 |
| STIP1 | 0.496654 | 8.57E-87 | 3.86E-84 |

| | | | |
|---|---|---|---|
| SRSF3 | 0.496551 | 4.00E-96 | 2.24E-93 |
| NUSAP1 | 0.49484 | 1.82E-57 | 2.74E-55 |
| DDX39A | 0.494825 | 1.45E-76 | 4.52E-74 |
| PTMS | 0.492335 | 4.30E-66 | 9.00E-64 |
| FDPS | 0.48889 | 8.62E-84 | 3.34E-81 |
| DEK | 0.488734 | 2.22E-77 | 7.38E-75 |
| TK1 | 0.488647 | 1.83E-75 | 5.47E-73 |
| PFN1 | 0.484557 | 8.12E-155 | 9.87E-151 |
| RPA3 | 0.484419 | 1.20E-82 | 4.45E-80 |
| SH2D2A | 0.48437 | 1.33E-87 | 6.13E-85 |
| ALYREF | 0.48303 | 1.41E-76 | 4.43E-74 |
| HSP90AA1 | 0.478993 | 9.02E-105 | 6.33E-102 |
| CHCHD2 | 0.478 | 3.66E-126 | 6.67E-123 |
| HIST1H1D | 0.477202 | 1.67E-48 | 1.88E-46 |
| MZT2A | 0.47638 | 7.90E-86 | 3.43E-83 |
| PGK1 | 0.475657 | 1.73E-74 | 5.05E-72 |
| HSP90AB1 | 0.474715 | 3.64E-65 | 7.29E-63 |
| CCND2 | 0.473701 | 2.98E-64 | 5.78E-62 |
| SMC2 | 0.472256 | 6.61E-73 | 1.81E-70 |
| TMSB10 | 0.471564 | 1.41E-138 | 4.29E-135 |
| COX8A | 0.47095 | 6.26E-124 | 1.04E-120 |
| CALR | 0.46942 | 2.00E-69 | 4.77E-67 |
| YBX1 | 0.467883 | 1.05E-99 | 6.40E-97 |
| FEN1 | 0.458769 | 8.34E-70 | 2.02E-67 |
| ASPM | 0.458482 | 1.29E-51 | 1.64E-49 |
| CFL1 | 0.456566 | 3.86E-137 | 9.40E-134 |
| GSTO1 | 0.452454 | 3.61E-81 | 1.30E-78 |
| C17orf49 | 0.450881 | 2.03E-77 | 6.80E-75 |
| IFNG | 0.449948 | 2.33E-55 | 3.34E-53 |
| FAM102A | -0.458119 | 1.26E-77 | 4.30E-75 |
| C1orf162 | -0.462886 | 9.51E-66 | 1.93E-63 |
| GIMAP7 | -0.472451 | 1.27E-76 | 4.03E-74 |
| ARHGAP15 | -0.490496 | 1.51E-94 | 8.25E-92 |
| XIST | -0.515785 | 4.83E-90 | 2.32E-87 |
| CD37 | -0.54178 | 5.57E-119 | 7.81E-116 |
| ZFP36L2 | -0.563201 | 3.13E-86 | 1.37E-83 |
| EGLN3 | -0.566437 | 4.69E-50 | 5.56E-48 |
| BTG1 | -0.566483 | 4.68E-82 | 1.72E-79 |
| SLFN5 | -0.614581 | 4.53E-114 | 4.23E-111 |

| | | | |
|---|---|---|---|
| STAT1 | -0.622469 | 1.37E-90 | 6.84E-88 |
| GAS5 | -0.625822 | 1.09E-115 | 1.13E-112 |
| ITGB1 | -0.661546 | 3.93E-111 | 3.12E-108 |
| TXNIP | -0.67967 | 2.81E-93 | 1.46E-90 |
| PNRC1 | -0.689702 | 1.31E-115 | 1.33E-112 |
| IFITM1 | -0.71378 | 3.17E-147 | 1.29E-143 |
| LIME1 | -0.719742 | 6.43E-140 | 2.13E-136 |
| IL7R | -0.773457 | 2.68E-90 | 1.30E-87 |
| SELL | -0.89022 | 5.63E-113 | 4.89E-110 |

**Table S2. Differentially expressed genes identified in $CD8^+$ T cells from citrullinated antigen-loaded B cell coculture as compared to those from native protein-loaded B cell coculture.** Differentially expressed genes were selected with average Log2 Fold change > 0.45. Wilcoxon rank sum test and the Benjamini–Hochberg (BH) method to adjust the *P* values for multiple testing. FC; fold change, FDR; false discovery rate.

| Genes | logFC | pvalue | FDR |
|---|---|---|---|
| GZMB | 1.680252 | 1.67E-197 | 8.69E-194 |
| TUBA1B | 1.510702 | 6.94E-206 | 5.07E-202 |
| STMN1 | 1.188824 | 1.08E-160 | 1.79E-157 |
| TUBB | 1.068337 | 9.48E-173 | 2.03E-169 |
| TYMS | 1.042696 | 3.18E-156 | 4.83E-153 |
| GZMA | 1.015152 | 1.15E-124 | 7.34E-122 |
| LAG3 | 1.006964 | 1.16E-169 | 2.35E-166 |
| HLA-DRB5 | 0.983214 | 9.86E-154 | 1.44E-150 |
| HMGN2 | 0.921186 | 2.34E-174 | 5.33E-171 |
| MKI67 | 0.907567 | 9.86E-136 | 8.37E-133 |
| HMGB2 | 0.889612 | 2.13E-135 | 1.77E-132 |
| HIST1H2AJ | 0.880285 | 2.77E-179 | 7.21E-176 |
| HIST1H4C | 0.852117 | 5.68E-115 | 3.05E-112 |
| RRM2 | 0.847094 | 1.83E-115 | 1.01E-112 |
| LDHA | 0.801128 | 1.09E-181 | 3.06E-178 |
| H2AFZ | 0.781895 | 2.85E-136 | 2.54E-133 |
| GAPDH | 0.780109 | 3.44E-252 | 1.25E-247 |
| TPI1 | 0.778896 | 1.67E-191 | 7.60E-188 |
| PCLAF | 0.729922 | 2.88E-126 | 1.95E-123 |
| TUBB4B | 0.700269 | 1.82E-99 | 6.92E-97 |
| ENO1 | 0.677933 | 5.69E-179 | 1.38E-175 |
| CCL3 | 0.674624 | 9.59E-63 | 1.19E-60 |
| EBP | 0.648116 | 2.59E-144 | 2.86E-141 |
| H2AFV | 0.627656 | 6.21E-141 | 5.96E-138 |
| HMGB1 | 0.62687 | 5.18E-190 | 2.10E-186 |
| MCM7 | 0.623114 | 6.17E-102 | 2.45E-99 |
| HIST1H1B | 0.619242 | 7.24E-89 | 2.05E-86 |
| PKM | 0.609685 | 9.37E-150 | 1.10E-146 |
| NASP | 0.606225 | 1.81E-117 | 1.02E-114 |
| CENPF | 0.605959 | 1.02E-79 | 2.12E-77 |
| HNRNPA2B1 | 0.605315 | 9.86E-199 | 6.00E-195 |
| TXN | 0.603407 | 8.28E-150 | 1.01E-146 |
| TMSB10 | 0.598053 | 1.54E-248 | 2.80E-244 |
| RAN | 0.594705 | 6.36E-153 | 8.60E-150 |
| HOPX | 0.594421 | 8.22E-56 | 8.74E-54 |
| UBE2S | 0.594038 | 3.40E-79 | 6.93E-77 |
| UBE2C | 0.591028 | 2.23E-73 | 3.75E-71 |
| TUBA1C | 0.581194 | 1.22E-90 | 3.74E-88 |

| | | | |
|---|---|---|---|
| NUSAP1 | 0.57439 | 3.07E-85 | 7.61E-83 |
| LGALS1 | 0.573358 | 5.15E-54 | 5.19E-52 |
| HSP90AB1 | 0.572378 | 1.73E-107 | 7.81E-105 |
| HSP90AA1 | 0.571401 | 4.51E-183 | 1.37E-179 |
| SNRPB | 0.567135 | 9.70E-152 | 1.22E-148 |
| IL2RA | 0.567061 | 9.37E-74 | 1.59E-71 |
| EIF4A1 | 0.566598 | 3.76E-159 | 5.97E-156 |
| H2AFX | 0.566552 | 9.86E-89 | 2.77E-86 |
| SLC25A5 | 0.563086 | 8.78E-135 | 6.81E-132 |
| ANXA2 | 0.562686 | 2.15E-99 | 8.10E-97 |
| TOP2A | 0.557433 | 8.04E-74 | 1.38E-71 |
| PA2G4 | 0.557146 | 1.42E-140 | 1.33E-137 |
| DUT | 0.555333 | 2.30E-85 | 5.74E-83 |
| FABP5 | 0.554857 | 2.01E-96 | 7.00E-94 |
| UBALD2 | 0.553473 | 1.51E-106 | 6.71E-104 |
| SRSF3 | 0.550498 | 1.49E-141 | 1.47E-138 |
| LMNB1 | 0.548845 | 3.32E-103 | 1.34E-100 |
| PCNA | 0.544232 | 3.86E-82 | 8.75E-80 |
| FDPS | 0.541941 | 6.00E-129 | 4.21E-126 |
| PGAM1 | 0.541555 | 1.25E-143 | 1.35E-140 |
| COX8A | 0.54149 | 1.01E-187 | 3.70E-184 |
| CTSW | 0.54076 | 2.10E-80 | 4.48E-78 |
| CARHSP1 | 0.53639 | 1.35E-114 | 7.15E-112 |
| RANBP1 | 0.534202 | 4.56E-105 | 1.91E-102 |
| TESC | 0.533389 | 3.64E-106 | 1.60E-103 |
| ANP32E | 0.53233 | 1.54E-105 | 6.51E-103 |
| TUBA4A | 0.528586 | 3.25E-102 | 1.30E-99 |
| PTMA | 0.527947 | 1.06E-142 | 1.10E-139 |
| STIP1 | 0.524876 | 2.40E-120 | 1.46E-117 |
| PTTG1 | 0.524212 | 1.59E-85 | 4.03E-83 |
| CCL4 | 0.521323 | 3.35E-29 | 1.47E-27 |
| CHCHD2 | 0.519488 | 1.41E-184 | 4.67E-181 |
| DEK | 0.515332 | 1.15E-103 | 4.73E-101 |
| SMC4 | 0.51043 | 1.53E-72 | 2.52E-70 |
| MAD2L2 | 0.510349 | 1.45E-118 | 8.67E-116 |
| HIST1H3B | 0.507592 | 3.41E-85 | 8.40E-83 |
| MCM5 | 0.507485 | 1.79E-88 | 4.87E-86 |
| ACTB | 0.501444 | 3.50E-163 | 6.08E-160 |
| TPX2 | 0.499801 | 3.54E-73 | 5.92E-71 |

| | | | |
|---|---|---|---|
| PGK1 | 0.499112 | 6.89E-111 | 3.49E-108 |
| RPA3 | 0.498567 | 3.41E-109 | 1.66E-106 |
| TMPO | 0.497396 | 5.10E-88 | 1.36E-85 |
| H2AFY | 0.496846 | 2.92E-118 | 1.69E-115 |
| MZT2A | 0.494981 | 6.01E-136 | 5.22E-133 |
| TK1 | 0.492992 | 1.42E-88 | 3.92E-86 |
| LINC01943 | 0.488737 | 7.54E-59 | 8.68E-57 |
| ASPM | 0.485107 | 4.25E-68 | 6.10E-66 |
| SMC2 | 0.482958 | 1.20E-88 | 3.35E-86 |
| PPIA | 0.479982 | 5.61E-222 | 6.83E-218 |
| EIF1AY | 0.479434 | 4.92E-128 | 3.39E-125 |
| DBI | 0.477801 | 3.49E-137 | 3.19E-134 |
| C12orf75 | 0.476857 | 2.65E-77 | 5.20E-75 |
| NME1 | 0.476488 | 4.25E-87 | 1.10E-84 |
| SNRPD1 | 0.475895 | 1.26E-111 | 6.48E-109 |
| CALR | 0.475253 | 1.22E-89 | 3.56E-87 |
| SIRPG | 0.473659 | 3.26E-104 | 1.35E-101 |
| TNF | 0.470591 | 7.94E-62 | 9.73E-60 |
| ANAPC11 | 0.469604 | 3.29E-115 | 1.79E-112 |
| ZWINT | 0.469388 | 4.43E-94 | 1.50E-91 |
| YBX1 | 0.469043 | 1.96E-134 | 1.49E-131 |
| FEN1 | 0.46723 | 9.15E-85 | 2.21E-82 |
| GNLY | 0.454986 | 2.90E-14 | 6.63E-13 |
| TPM3 | 0.453323 | 4.69E-153 | 6.59E-150 |
| IFNG | 0.452884 | 3.31E-65 | 4.31E-63 |
| JPT1 | 0.45185 | 3.63E-90 | 1.09E-87 |
| CFL1 | 0.450691 | 6.11E-164 | 1.11E-160 |
| CDKN3 | 0.450674 | 5.11E-75 | 9.13E-73 |
| GAS5 | -0.451008 | 2.26E-76 | 4.31E-74 |
| EGLN3 | -0.457861 | 3.01E-33 | 1.53E-31 |
| IL10RA | -0.460847 | 3.48E-94 | 1.19E-91 |
| PDCD4 | -0.461499 | 2.49E-83 | 5.71E-81 |
| HLA-B | -0.461781 | 8.78E-214 | 8.01E-210 |
| ZFP36L2 | -0.476181 | 2.43E-74 | 4.25E-72 |
| YPEL3 | -0.482795 | 2.20E-85 | 5.54E-83 |
| TXNIP | -0.487105 | 2.49E-60 | 2.94E-58 |
| CD37 | -0.520516 | 5.55E-131 | 3.97E-128 |
| XIST | -0.5284 | 5.49E-123 | 3.39E-120 |
| IL7R | -0.542038 | 1.04E-51 | 9.53E-50 |

| | | | |
|---|---|---|---|
| SELL | -0.552769 | 2.36E-53 | 2.29E-51 |
| SLFN5 | -0.594256 | 6.98E-125 | 4.55E-122 |
| BTG1 | -0.61098 | 1.55E-110 | 7.76E-108 |
| ITGB1 | -0.667915 | 4.67E-135 | 3.70E-132 |
| PNRC1 | -0.685151 | 2.69E-133 | 2.01E-130 |
| LIME1 | -0.74092 | 1.59E-169 | 3.05E-166 |

**Table S3. Differentially expressed genes identified in CD8$^+$ T cells from citrullinated antigen-loaded B cell coculture as compared to those from citrullinated antigen with IRAK4 inhibitor-stimulated B cell coculture.** Differentially expressed genes were selected with average Log2 Fold change > 0.45. Wilcoxon rank sum test and the Benjamini–Hochberg (BH) method to adjust the *P* values for multiple testing. FC; fold change, FDR; false discovery rate.

| | **RA patients** |
|---|---|
| Total Number | 51 |
| Gender, Female/Male | 8/43 |
| Age (years) - mean±SD | 66.55±10.41 |
| Smoking (%) | 67.5 |
| Disease activity | |
| ACPA positivity (%) | 100 |
| CCP value | 213.71±67.97 |
| RF positivity (%) | 88.24 |
| CDAI score - mean±SD | 15.70±12.51 |
| DAS28-CRP | 3.24±1.6 |
| DAS28-ESR | 4.17±2.04 |
| MTX use (%) | 41.18 |

**Table S4. Demographic characteristics of RA patients.**

| TotalSeq C-barcode | Marker | Sequence | Clone |
|---|---|---|---|
| 34 | CD3 | CTCATTGTAACTCCT | UCHT1 |
| 45 | CD4 | GAGGTTAGTGATGGA | SK3 |
| 138 | CD5 | CATTAACGGGATGCC | UCHT2 |
| 46 | CD8 | GCGCAACTTGATGAT | SK1 |
| 53 | CD11c | TACGCCTATAACTTG | S-HCL-3 |
| 81 | CD14 | CAATCAGACCTATGA | M5E2 |
| 50 | CD19 | CTGGGCAATTACTCG | HIB19 |
| 100 | CD20 | TTCTGGGTCCCTAGA | 2H7 |
| 181 | CD21 | AACCTAGTAGTTCGG | Bu32 |
| 180 | CD24 | AGATTCCTTCGTGTT | ML5 |
| 85 | CD25 | TTTGTCCTGTACGCC | BC96 |
| 154 | CD27 | GCACTCCTGCATGTA | O323 |
| 386 | CD28 | TGAGAACGACCCTAA | CD28.2 |
| 389 | CD38 | TGTACCCGCTTGTGA | HIT2 |
| 63 | CD45RA | TCAATCCTTCCGCTT | HI100 |
| 87 | CD45RO | CTCCGAATCATGTTG | UCHL1 |
| 84 | CD56 | TTCGCCGCATTGAGT | QA17A16 |
| 168 | CD57 | AACTCCCTATGGAGG | QA17A04 |
| 147 | CD62L | GTCCCTGCAACTTGA | DREG-56 |
| 146 | CD69 | GTCTCTTGGCTTAAA | FN50 |
| 5 | CD80 | ACGAATCAATCTGTG | 2D10 |
| 6 | CD86 | GTCTTTGTCAGTGCA | IT2.2 |
| 246 | CD122 | TCATTTCCTCCGATT | TU27 |
| 390 | CD127 | GTGTGTTGTCCTATG | A019D5 |
| 55 | CD138 | ACTCTTTCGTTTACG | MI15 |
| 140 | CD183 | GCGATGGTAGATTAT | G025H7 |
| 144 | CD185 | AATTCAACCGTCGCC | J252D4 |
| 143 | CD196 | GATCCCTTTGTCACT | G034E3 |
| 148 | CD197 | AGTTCAGTCAACCGA | G043H7 |
| 88 | CD279 | ACAGCGCCGTATTTA | EH12.2H7 |
| 828 | FcRL4 | CGATTTGATCTGCCT | 413D12 |
| 829 | FcRL5 | TCACGCAGTCCTCAA | 509f6 |
| 215 | BAFF-R | CGAAGTCGATCCGTA | 11C1 |
| 179 | CX3CR1 | AGTATCGTCTCTGGG | K0124E1 |
| 366 | CXCR4 | TCAGGTCCTTTCAAC | 12G5 |
| 159 | HLA-DR | AATAGCGAGCAAGTA | L243 |
| 384 | IgD | CAGTCTCCGTAGAGT | IA6-2 |

| | | | |
|---|---|---|---|
| 375 | IgG-Fc | CTGGAGCGATTAGAA | M1310G05 |
| 136 | IgM | TAGCGAGCCCGTATA | MHM-88 |
| 224 | TCRab | CGTAACGTAGAGCGA | IP26 |
| 139 | TCRrd | CTTCCGATTCATTCA | B1 |
| 89 | TIGIT | TTGCTTACCGCCAGA | A15153G |
| 7 | PD-L1 | GTTGTCCGACAATAC | 29E.2A3 |
| 9 | B7-H2 | GTGCATTCAACAGTA | 2D3 |
| 151 | CTLA-4 | ATGGTTCACGTAATC | BNI3 |
| 152 | LAG-3 | CATTTGTCTGCCGGT | 11C3C65 |
| 170 | BTLA | GTTATTGGACTAAGG | MIH26 |
| 171 | ICOS | CGCGCACCCATTAAA | C398.4A |
| 352 | FceRIa | CTCGTTTCCGTATCG | AER-37 (CRA-1) |
| 363 | IL4-Ra | CCGTCCTGATAGATG | G077F6 |
| 397 | CCR3 | ACCAATCCTTTCGTC | 5E8 |
| 935 | CD74 | CTGTAGCATTTCCCT | LN2 |
| 219 | IFNyRa | TGTGTATTCCCTTGT | GIR-208 |
| 405 | TLR4 | GCTTAGCTGTATCCG | HTA125 |

**Table S5. List of CITE-seq antibodies used in scRNA-seq.**

| Marker | Clone | Fluorescence | Titration | Company |
|---|---|---|---|---|
| **CD3** | SK7 | FITC/APC | 1:100 | BD Biosciences |
| **CD4** | RPA-T4 | BV605 | 1:100 | BioLegend |
| **CD8** | RPA-T8 | Alexa Fluor 700 | 1:100 | BD Biosciences |
| **CD69** | FN50 | PerCP-Cy5.5 | 1:50 | BD Biosciences |
| **MPO** | 5B8 | PE | 1:10 | BD Biosciences |
| **TLR4** | HTA125 | APC/BV421 | 1:25 | BioLegend |
| **IRAK4** | L29-525 | PE | 1:25 | BD Biosciences |
| **CD11c** | 3.9 | BV605/BV421 | 1:100 | BioLegend |
| **Granzyme B** | GB11/QA16A02 | PE/PerCP-Cy5.5 | 1:50 | BD Biosciences/BioLegend |
| **IFNγ** | 4S.B3/B27 | APC-Cy7/PE | 1:25 | BioLegend |
| **CD107a** | H4S3 | APC-Cy7 | 1:100 | BioLegend |
| **IL-6** | MQ2-13A5 | PE-Cy7 | 1:25 | BioLegend |
| **CD80** | L307.4 | AF700 | 1:50 | BioLegend |
| **CD86** | BU63 | FITC | 1:50 | BioLegend |
| **CD19** | HIB19 | BV421/APC-Cy7 | 1:50 | BioLegend |
| **IL-8** | 8CH | AF488 | 1:25 | Thermo Fisher Scientific |
| **CD27** | O323 | PerCP-Cy5.5 | 1:50 | BioLegend |
| **T-bet** | 4B10 | PE-Cy7 | 1:50 | BioLegend |
| **CD38** | S17015A/HIT2 | AF700/APC-Cy7 | 1:50 | BioLegend |
| **CXCR1** | 8F1/CXCR1 | PE-Cy7 | 1:50 | BioLegend |
| **CXCR2** | 5E8/CXCR2 | PerCP-Cy5.5 | 1:25 | BioLegend |
| **HLA-DR** | L243 | AF488 | 1:100 | BioLegend |
| **HLA-A,B,C** | W6/32 | APC-Cy7 | 1:100 | BioLegend |
| **CD40** | 5C3 | PE-Cy7 | 1:50 | BioLegend |
| **pSyk (Tyr348)** | moch1ct | PE | 1:50 | Thermo Fisher Scientific |
| **IgD** | IA6-2 | BUV737 | 1:100 | BD Biosciences |
| **TLR2** | W15145C | PE-Cy7 | 1:25 | BioLegend |
| **TLR7** | S18024F | FITC | 1:25 | BioLegend |
| **Perforin** | dG9 | BV421 | 1:25 | BioLegend |
| **Granulysin** | RB1 | AF488 | 1:25 | BD Biosciences |
| **His-tag** | J095G46 | FITC | 1:25 | BioLegend |
| **CXCR5** | J252D4 | BV421 | 1:50 | BioLegend |
| **CX3CR1** | 2A9-1 | FITC | 1:50 | BioLegend |

**Table S6. List of antibodies used in flow cytometry.**
